# Three-Dimensional Molecular Atlas of Octopus Arm Neuroanatomy Highlights Spatial and Functional Complexity

**DOI:** 10.1101/2024.04.14.589438

**Authors:** Gabrielle C. Winters-Bostwick, Sarah E. Giancola-Detmering, Caleb J. Bostwick, Robyn J. Crook

**Affiliations:** Department of Biology, San Francisco State University, San Francisco, CA, USA; Jupiter Bio LLC, Daly City, CA, USA

## Abstract

Octopus arms, notable for their complex anatomy and remarkable flexibility, have sparked significant interest within the neuroscience community. However, there remains a dearth of knowledge about the molecular and functional identities of various cell types in the arm’s nervous system. To address this gap, we used hybridization chain reaction (HCR) to identify distinct neuronal types in the arms of the pygmy octopus, *Octopus bocki*, including putative dopaminergic, octopaminergic, serotonergic, GABAergic, glutamatergic, cholinergic, and peptidergic neurons. We obtained high-resolution multiplexed fluorescent images at 0.28x0.28x1.0 μM voxel size from 10 arm base and arm tip cross sections (each 50 μM thick) and created three-dimensional reconstructions of the axial ganglia, illustrating the spatial distribution of multiple neuronal populations. Our analysis unveiled anatomically distinct and molecularly diverse scattered neurons, while also highlighting multiple populations of dense small excitatory neurons that appear uniformly distributed throughout the cortical layer. Our data provide new insights into how different types of neurons may contribute to the ability of an octopus to interact with its environment and execute complex tasks. In addition, our findings establish a benchmark for future studies, allowing pioneering exploration of octopus arm molecular neuroanatomy, and offering exciting new avenues in invertebrate neuroscience research.

## Introduction

Octopuses are recognized for their complex behavior and remarkable cognitive abilities, captivating scientists across diverse disciplines. These enigmatic molluscs display dynamic camouflage, advanced hunting strategies, and problem-solving skills unparalleled among invertebrates. Perhaps the most striking feature of an octopus is its set of eight autonomous arms, each executing both decentralized and coordinated functions with remarkable proprioceptive abilities, fine motor control, and advanced tactile sensation.

Along the entire length of each arm lies an axial nerve cord (ANC), a strand of bead-like ganglia that houses neuronal networks responsible for supporting sensory, motor, and integrative functions within and across arms. Each ganglion, associated with a single sucker, possesses a repeated general cytoarchitecture with a dense cortical layer (CL) of neuronal somata surrounding a neuropil (NP) of projections and synapses. Dorsal to the CL and NP is a tract of axons (the cerebrobrachial tract or CBT) that runs the length of the cord, spanning the ganglia^1,2^. Within the cortical layer, each ganglion houses a dense population of small (∼8-10 μM diameter) morphologically homogenous neurons, as well as less numerous and anatomically diverse large neurons (up to ∼25 μM)^3^

Even though approximately 60-70% (300-350 million out of about 500 million total) of the neurons in an octopus’s nervous system are located in the arms^1,3^,our understanding of the cellular composition and neurotransmission along the axial nerve cord is limited^2^.Therefore, the task of characterizing cellular diversity at multiple locations along the arm is not just intriguing but a fundamental step in broadening our grasp of nervous system evolution and function. With this in mind, we endeavor to define molecular identities of cell populations within this morphologically diverse complement of neurons, a pivotal step in understanding these animals’ impressive capabilities. This molecular study is an integral component of our overarching goal to characterize the neural connectome of octopus arms.

To date, the prevailing model of neurotransmission in the axial nerve cord is characterized by the employment of three distinct types of cholinergic neurons that innervate the surrounding muscle fibers^4^. This model is bolstered by the ubiquitous immunohistochemical localization of peripheral type choline acetyltransferase (pChAT) in the axial nerve cord and other neuronal arm structures, as demonstrated by Sakaue et al. in 2013^5^. This evidence emphasizes acetylcholine’s prominent role in the octopus’s neuromuscular junctions, highlighting its involvement in a range of functions from motor control to sensory processing. In addition to acetylcholine, serotonin has been identified in neurons and the surrounding tissues of the arm and proposed as a neuromodulator of synaptic transmission, as opposed to a primary neurotransmitter^6^.

Nevertheless, the neurochemical framework of cephalopod nervous systems, including that of the octopus, likely extends beyond acetylcholine and serotonin. In an effort to create a comprehensive molecular map of these systems, our study has incorporated a diverse array of molecular markers. We selected these markers to identify representatives of key functional classes of neurons: excitatory (e.g., glutamatergic), inhibitory (e.g., GABAergic, possibly glycinergic), monoaminergic (e.g., dopaminergic, serotonergic, octopaminergic), and peptidergic. This selection is grounded in the pivotal roles that these molecules play in essential neurological processes

Glutamate, primarily an excitatory neurotransmitter, is strongly implicated in the vertical lobe memory circuit of the octopus brain, as a facilitator of synaptic transmission at key neural junctions and playing a critical role in visual learning and spatial memory plasticity. GABA, in contrast, serves primarily as an inhibitory neurotransmitter across diverse taxa^7^, forming a dynamic balance essential for rapid synaptic transmission and modulation^8^. This balance may therefore also be vital for the octopus arm’s rapid response mechanisms in prey capture and sensory integration.

Dopamine and serotonin, traditionally linked with neuromodulation of reward-motivated behavior and mood regulation across vertebrate and invertebrate taxa^9,10^, are hypothesized to have unique functional roles in cephalopods. Dopamine may be involved in octopus feeding behaviors, as evidenced by its role in molluscan feeding circuits^11–13^ and in the octopus vertical lobe memory circuitry^3,14^. Serotonin is also posited to play a part in learning processes, specifically long-term potentiation (LTP)^15^ and possibly in the control of chromatophore expansion for camouflage^16^. Octopamine, a neuromodulator found across diverse taxa, is also implicated in octopus learning and memory, specifically short-term facilitation and suppression of LTP^17^ as well as a variety of other functions in invertebrates such as modulation of motor control^18^ and aggression^19^, which may suggest its potential role in the orchestration of arm movements for predatory behaviors in octopuses.

The functions of neuropeptides such as FLRIamide and bradykinin-like neuropeptide within octopus tissues remain less elucidated. FMRFamide-like peptides^20^ like FXRIamides (specifically FLRIamide in Octopus) are believed to exert a neuromodulatory influence in cephalopod brains^21,22^, impacting neural functions beyond simple neurotransmission, and FLRIamide specifically has been localized to the large efferent neurons of the vertical lobe^14^. These peptides may play roles in a spectrum of physiological processes across cephalopod nervous systems, ranging from pain modulation to behavior and metabolic regulation ^15,23,24^. Bradykinin-like neuropeptide, in this case representing a potential innovation in molluscan neurochemistry, presents an intriguing frontier for future neurobiological research, with its function yet to be fully understood.

Our study aims to characterize the neural connectome of octopus arms using multiplexed Hybridization Chain Reaction (HCR) probes. This approach has enabled us to map the expression of mRNAs associated with both established neurotransmitters and cephalopod-specific neurosecretory molecules. We have successfully created 3D reconstructions of the axial nerve cord ganglia tip and base, revealing the spatial distribution of various cell types. Utilizing these data, we have formulated hypotheses regarding the functions and dynamics of different neurochemicals within the octopus arm’s nervous systems. By integrating cellular, molecular, anatomical, and neurochemical insights, our research aims to significantly enhance the understanding of cephalopod neurobiology, shedding light on its broader implications for the study of nervous system evolution and function in marine organisms. This research not only enriches our comprehension of octopus neurobiology, but also underscores the remarkable complexity of one of nature’s most advanced biological systems.

## Results

### Identification and cloning of transcripts used for mapping neuronal populations

We used eight molecular markers to identify and classify diverse types of neurons in octopus axial nerve cords. Of these, six were low molecular weight neurotransmitters (glutamate, GABA, octopamine, serotonin, dopamine and acetylcholine) and two were putative neuropeptide precursors (FLRIamide, and bradykinin-like neuropeptide). In contrast to the neuropeptides for which an mRNA sequence is translated into a neuropeptide precursor (prepropeptide), low molecular weight neurotransmitters cannot be directly localized by *in situ* hybridization. *In situ* hybridization (including HCR used in this study) works by localizing mRNA transcripts in fixed cells or tissues using a detectable probe. Therefore, to visualize cells containing these low molecular weight markers, we used probes designed against mRNAs encoding proteins associated with the synthesis and/or transport of these markers. For example, to identify putative glutamatergic cells, we used probe sets designed against two different mRNAs encoding proteins associated with the transport of glutamate: vGluT (vesicular glutamate transporter) and slc6a15/18 (a solute carrier protein involved in glutamate precursor transport^25^ and previously used as a glutamatergic neuron marker in octopus^26^). A complete list of proteins whose mRNA sequences were used to identify neuronal markers cells in this study can be found in Table 1, and in Table S1, which also contains *Octopus bocki* mRNA sequences used to create HCR probes and lot numbers for Molecular Instruments HCR probes.

**Table 1:** Cell types and markers examined in this study.

| Cell Type (Neurotransmitter) | Transcript name | Shorthand HCR probe |
| --- | --- | --- |
| **Cholinergic cells (Acetylcholine)** | Choline acetyltransferase (ChAT) | ChAT B1 |
| **Dopaminergic cells (Dopamine)** | Tyrosine Hydroxylase (TyH) | TyH B4 |
| **Excitatory cell (Maybe Glutamate)** | Excitatory Amino Acid Transporter 1 (EAAT1) | EAAT1 B4 |
| **GABAergic/Inhibitory cells (GABA)** | vesicular GABA Transporter/vesicular Inhibitory Amino Acid Transporter (vGAT) | vGAT B1 |
| **Glutamatergic cells (Glutamate)** | vesicular Glutamate Transporter (vGluT) | VGluT B3 |
| **Glutamatergic or GABAergic cell (Glutamate or GABA)** | Solute Carrier Family 6 Member 15/18 (slc6a15/18) | slc6a15/18 B5 |
| **Glycinergic Cell (Glycine)** | Glycine Transporter (GlyT) | GlyT B4 |
| **Octopaminergic cells (Octopamine)** | Tyramine $\beta$ hydroxylase (TβH) | TβH B5 |
| **Peptidergic cells (FLRlamide)** | FLRlamide peptide precursor (FLRI) | FLRI B2 |
| **Peptidergic cells (Bradykinin-like)** | Bradykinin-like peptide precursor (Brady) | Bradykinin-like B1 |
| **Serotonergic cells (Serotonin)** | Tryptophan Hydroxylase (TpH) | TpH B3 |

All mRNA sequences were identified from our *O. bocki* neuronal transcriptomes (SRA accession number PRJNA1050799) and their conserved evolutionary identity was confirmed by multiple sequence alignments and gene trees (see supplementary materials for trees and aligned sequences). We also cloned most of the selected transcripts from *O. bocki* arm tissues (with the exception of slc6a15/18) in order to verify their sequence and tested them using chromogenic digoxygenin labelled *in situ* hybridization assays before moving on to HCR. Representative images of chromogenic assays for each transcript’s expression in octopus arms can be found in the supplement. All chromogenic assay results were consistent with those of HCR assays.

One putative secretory molecule in this study is referred to as “bradykinin-like” neuropeptide, consistent with recent publications^14^. It loosely aligns with other molluscan (gastropod and bivalve) sequences annotated as “bradykinin-like neuropeptide precursor”. However, we cannot confirm that the cephalopod bradykinin-like neuropeptide precursor shares any evolutionary origin or function with that of other molluscs, as the e-values for all non-cephalopod BLASTp results were 0.02 or greater when we used the *O. bocki* protein sequence as a query. It is therefore possible that this putative secretory molecule is a cephalopod innovation with unknown function. The amino acid sequences for *Octopus bimaculoides* and *Octopus sinensis* used in our multiple sequence alignments (see supplementary materials) are both manually translated from transcripts annotated as ncRNA despite the presence of a signal peptide in the amino acid sequence.

The transcript referred to throughout this manuscript as “tyramine β hydroxylase” (TβH) is homologous to sequences annotated as both “tyramine β hydroxylase” and “dopamine β hydroxylase” (DβH) (see supplementary materials for multiple sequence alignments). Multiple sequence alignments of invertebrate homologs did not clarify the identity of this molecule, as annotations in closely related species are inconsistent, and functional domains were not distinguishable based on available protein sequences. This distinction is important for determining whether the neurotransmitter produced by cells expressing this molecule is octopamine (from TβH) or norepinephrine (from DβH). Therefore, we used expression data to deduce the molecule’s likely identity and putative function. Here we make the assumption that a cell expressing dopamine β hydroxylase would also exhibit a mechanism for dopamine synthesis. The absence of any colocalization between DβH/TβH and TyH (which would indicate the presence of dopamine) suggests that there is no known dopamine synthesis in cells expressing DβH/TβH (see expression profiles in Figures 2 and 3 below). Although it is possible that dopamine is synthesized by a pathway other than that involving TyH, for this study we refer to this ambiguous sequence as tyramine β hydroxylase (TβH) and interpret its expression as an indication of the presence of octopamine.

**Figure 1:**
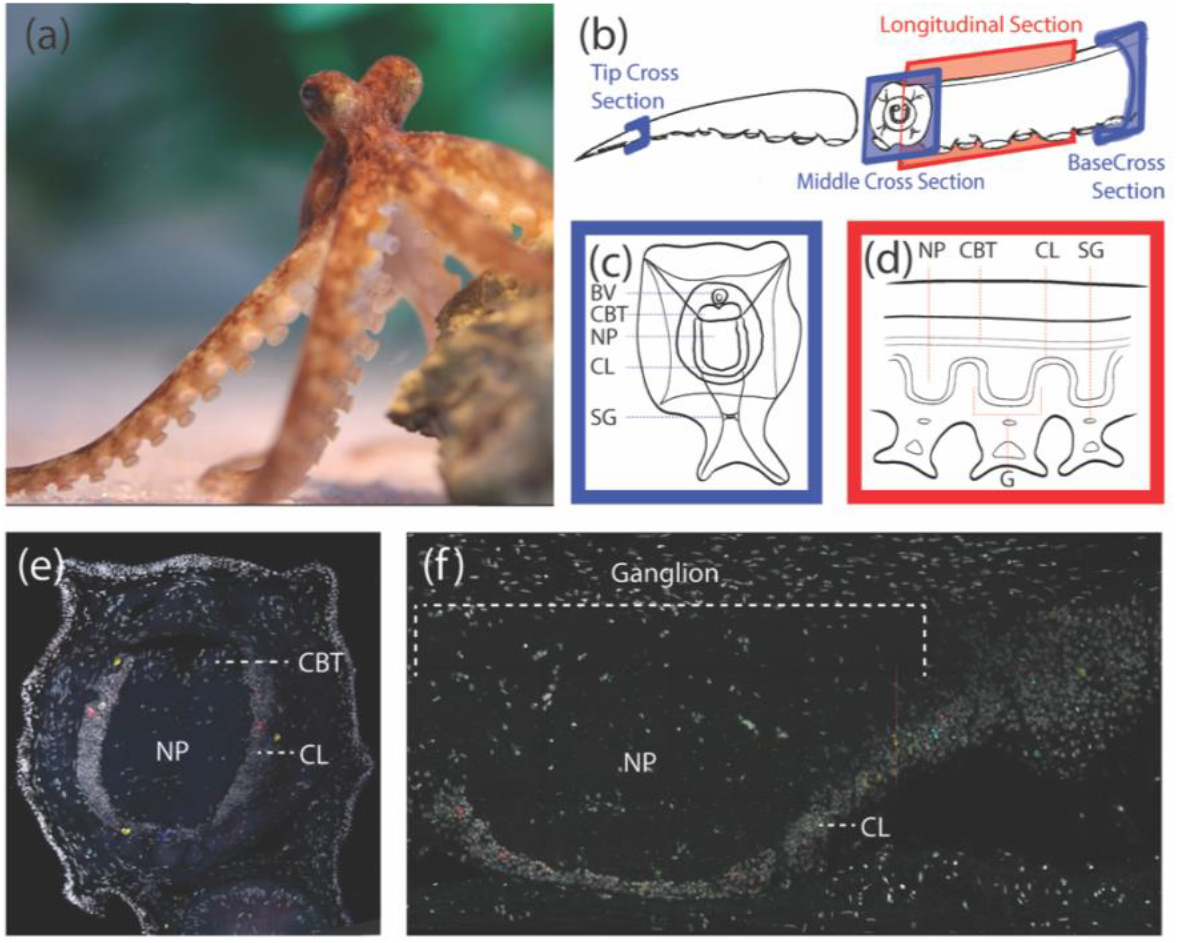
Arm neuroanatomy of *Octopus bocki*. a: Adult female *O. bocki*. b: Line schematic of an intact arm with longitudinal and cross-sectional plane locations indicated in red and blue (respectively). c-d: Line schematics illustrating neuroanatomy of octopus arms in cross (c) and longitudinal (d) section. e-f. Confocal microscopy images of cell nuclei (gray= DAPI) in a whole arm tip cross section (e) and a longitudinal section of one axial nerve cord ganglion. BV: Blood Vessel. NP: Neuropil. CL: Cortical Layer. G: Ganglion. CBT: Cerebro-Brachial Tract. SG: Sucker Ganglion

**Figure 2:**
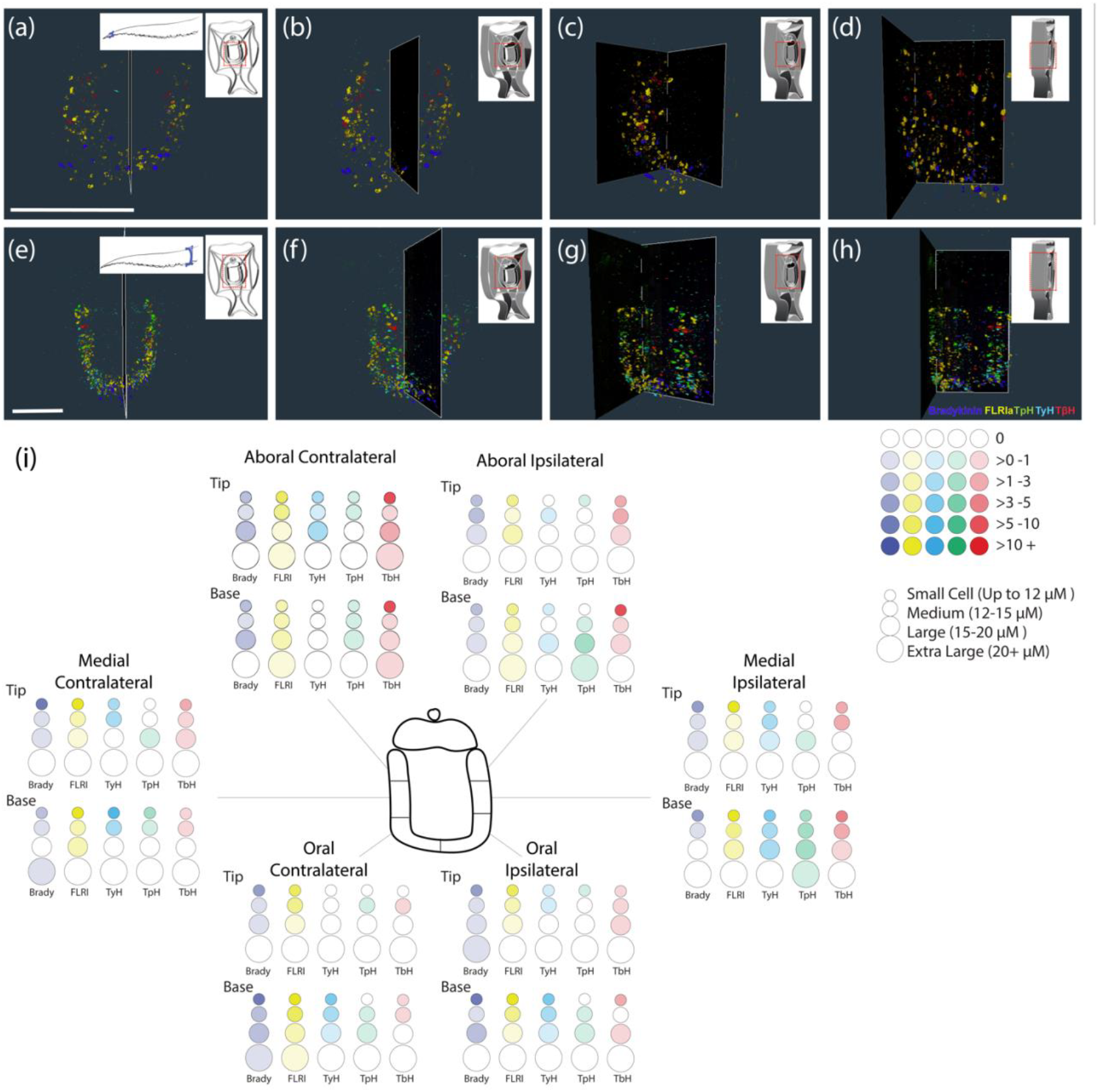
Three-dimensional reconstruction of neurochemical expression in diverse scattered cortical neurons in *O. bocki* ANC at the arm tip (a-d) and base (e-h). Each series (a-d and e-h) shows a molecular map of five distinct cell-type markers progressively rotated around a Y-axis. Reconstructions, composed of ten serial 50 μM cross sections, represent 500μM of the ANC. Scale bars in panels a and e are 350 μM. Panel i shows the breakdown of count averages (n=3) for each cell type in arm base and tip slices. The ANC cortical layer is divided into six locations based on the oral/medial/aboral and ipsilateral/contralateral position. Within each location the number of cells of each size (small, medium, large and extra large) were counted in a maximum intensity projection of a 50 μM z-stack for each neurotransmitter. The opacity of the filling for each cell size in the schematic indicates the number of cells of that size in that section (see Tables S2-S6 for counts and analyses). Colors - blue: bradykinin-like neuropeptide; yellow: FLRIamide neuropeptide; green: tryptophan hydroxylase; cyan: tyrosine hydroxylase; red: tyramine β hydroxylase. See supplementary materials (animation S1) for animated videos of rotating reconstructions.

**Figure 3:**
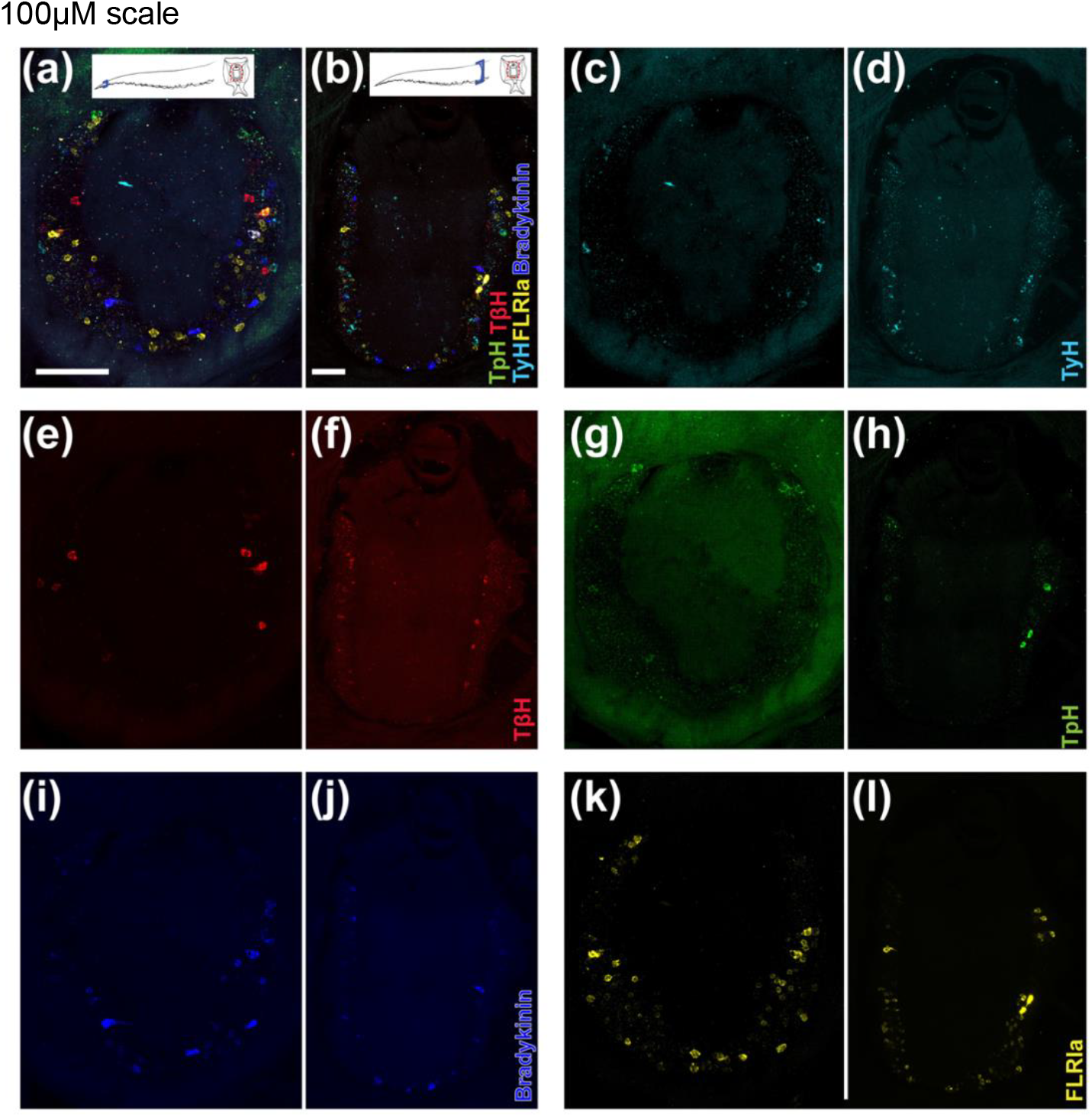
Cross section slices (50 μM thick) from arm tip (a, c, e, g, i, k) and base (b, d, f, h, j, l) reveal expression maps for distinct scattered cortical neuron populations using five cell type markers. Panels a and b show a composite image of cells expressing all five markers and the remaining panels show expression profiles for tyrosine hydroxylase in cyan (c-d), tyramine β hydroxylase in red (e-f), tryptophan hydroxylase in green (g-h), bradykinin-like neuropeptide in blue (i-j), and FLRIamide in yellow (k-l). Scale bars are 100 μM.

### Expression profiles of distinct neuronal populations

The neurons detected using our eight HCR probes were generally found in one of three broad expression patterns/profiles that we defined based on location, density, and (to some extent) soma size. These profiles are “scattered cortical neurons” (SCNs), “dense small cortical neurons” (DSCNs), and “arborizing neuropil neurons” (ANNs). Although, most neuronal types identified in this study fall primarily into one of these three designations, there are exceptions that are also described below.

#### Scattered cortical neurons (SCNs) and 3D reconstruction of the axial nerve cord

The first expression profile, referred to as “scattered cortical neurons” (SCNs) can be described as distinctly labelled, typically morphologically large (>10 μM, but up to 25 μM) neurons scattered throughout the axial nerve cord cortical layer. SCNs are typically dispersed, such that individual neuronal somata (and in many cases their projections) are easily visually distinguished. The transcripts whose expression demonstrated the SCN profile were FLRIamide neuropeptide, bradykinin-like neuropeptide, tyrosine hydroxylase (TyH-indicating dopamine expression), tyramine β hydroxylase (TβH-indicating octopamine expression), and tryptophan hydroxylase (TpH-indicating serotonin expression) (Figures 2 and 3). With few exceptions (see below), each of these transcripts appears to be expressed in a distinct subset of SCNs.

Figure 2 shows three-dimensional reconstructions of the arm tip (a-d) and base (e-h) using SCN transcript expression. Each stack is composed of ten 50 μM thick slices, allowing us to reconstruct 500 μM of the ANC at each position along the arm at a resolution of 0.28x0.28x1.0 μM voxel size (see supplementary animation S1). Still images of these reconstructions are shown in Figure 2 in a series as the arm is progressively rotated around a vertical (oral-aboral) axis, facilitating visualization of both regional specificity and heterogeneity for each type of SCN identified.

Some of the more faintly labelled and/or smaller cells are difficult to visualize in the 3D reconstruction, so we also compared cell counts (averaged across three 50 μM slices each of arm tip and base - see Tables S2a and S2b for raw counts and averages) of different sized soma labelled by each SCN transcript in six regions of the ANC CL from the arm base and tip to reveal expression trends. Figure 2i illustrates the distribution of the mean value for each cell count per 50 μM slice using a simplified schematic. We compared the average total number of neurons (all sizes in all regions) labelled by each SCN marker in the arm base vs arm tip (Table S3) as well as the number of cells of each size (“Small”: under 12 μM soma diameter; “Medium”:12-15 μM soma diameter; “Large”: 15-20 μM soma diameter; “Extra Large”: greater than 20 μM soma diameter) for each marker in the arm base vs arm tip (Table S4). We also compared the number of neurons in each vertical (oral/medial/aboral) and lateral (ipsilateral vs contralateral to the sucker) stratum for each transcript in the base and also in the tip (Tables S5 and S6).

The numbers of bradykinin-like neuropeptide-positive neurons and octopaminergic (TβH) neurons were not significantly different overall nor when comparing per cell size category between the arm base and the tip. However, the horizontal stratification of these two markers along the oral-aboral axis is immediately apparent in the 3D reconstruction of the arm (Figure 2 and supplementary animation S1). According to cell count analyses (Table S5), octopaminergic neurons, labelled in red by TβH, are most abundant in the medial and aboral region of the CL and nearly absent in the oral/ventral stratum. There are more octopaminergic neurons on the ipsilateral (to the sucker) side than the contralateral side of the ANC in the arm base, but not the tip (Table S6). Bradykinin-like neuropeptide-positive neurons in blue appear to cluster toward the medial and oral area (Table S5). Note that the stratification of bradykinin-like neuropeptide is statistically significant in the arm base, but not in the tip, possibly due to small sample sizes or the presence of faintly labelled small cells in the aboral regions that are difficult to visualize in the 3D reconstruction.

The density of FLRIamide-positive neurons appears consistent between the base and tip in the 3D reconstructions and individual slices (Figure 2, 3, and supplementary animation S1). Based on cell count analyses, there were no significant differences between base and tip for any specific cell size (Table S4), but the tip has significantly fewer FLRIamide-positive neurons overall than the base (Table S3). This may be a reflection of the fact that there are fewer cells altogether in the tip than in the base. Cell count data suggests that FLRIamide expressing neurons are most numerous in the Medial CL (Table S5) and are more abundant on the contralateral side (than the side ipsilateral to the sucker) in the tip, but not in the base (Table S6).

There were significant differences between the overall tip and base cell counts for dopaminergic (TyH) and serotonergic (TpH) neurons (Table S3) and the magnitude of this divergence is clearly visible in the 3D reconstruction (Figure 2, Table S4, and supplementary animation S1). Furthermore, dopaminergic neurons have distinct stratification patterns between the arm base and tip (Figure 2, Figure 3, and Table S5). In the arm base they can be seen in cyan throughout the CL, but are primarily clustered toward its lower medial and oral regions (Figure 2e-h and Figure 3a-d). In the tip dopaminergic cell labelling is faint, making them nearly undetectable in the 3D reconstruction, but they are most numerous in the aboral region of the CL. Despite the presence of especially large serotonergic neurons in the aboral region of the arm base (and in few slices of the tip), visually skewing our models to suggest aboral stratification (Figures 2-3 and supplemental animation S1), according to cell count analyses there are no significant differences between vertical (Table S5) or lateral (Table S6) distribution in these neurons (except for one comparison in the arm base where there are slightly more cells in the medial region than in the oral region - Table S5).

Representative images of each transcript’s expression profile in arm base and tip slices (Figure 3) highlight the remarkable size range for the SCNs. All six markers are expressed in cell somata ranging from in size from small to Extra Large except for TyH, which is found in Small, Medium, and Large but not Extra Large somata. For all transcripts except TpH, the most abundant cell size is small in the base and tip sections. For TpH, there are more Medium and Large cell bodies than small ones in the tip, but not in the base.

#### Dense Small Cortical Neurons: Excitatory and inhibitory cells

The numerous “dense small cortical neurons” (DSCNs) appear to be morphologically homogeneous (generally 8-10 μM diameter and nearly spherical) and very densely packed with a high nucleus to cytoplasm ratio. The compact nature of these smaller neurons, in contrast to that of the SCNs, presents a challenge when it comes to distinguishing individual cell somata. Nevertheless, our HCR study has revealed multiple types of neurons within this population.

The most abundant neurons in the DSCN layer are cholinergic (labelled by choline acetyltransferase - Figure 4b and c) and glutamatergic (labelled by vesicular glutamate transporter (vGluT) and slc6a15/18-Figure 4a, c, and g-i). Note that vGluT indicates glutamate secretion by a cell, while slc6a15/18 is a solute carrier protein implicated in glutamate synthesis but not in secretion). Nearly all of the observed cells that express slc6a15/18 also expressed vGluT. While the vast majority of the DSCNs appear to be either exclusively glutamatergic or cholinergic and these neurons are homogenously and evenly mixed throughout the cortical layer, there are some less frequent exceptions to this observation. One notable exception is the occasional colocalization of excitatory amino acid transporter 1 (EAAT1) with the glutamatergic markers (Figure 4c) but not with ChAT (Figure 4b).

**Figure 4:**
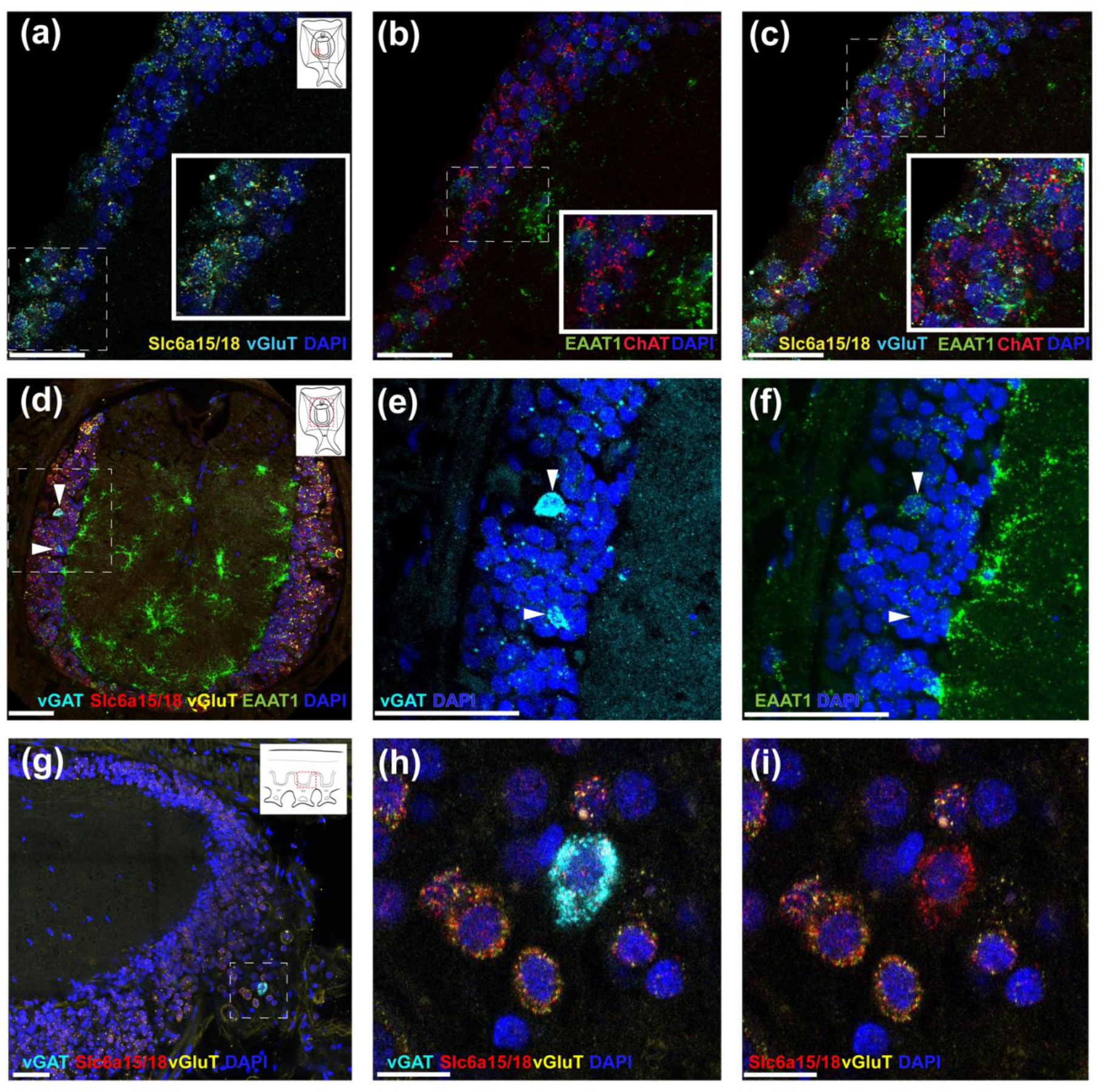
Densely packed morphologically homogenous small cortical neurons (DSCNs) predominantly express excitatory cell markers in addition to extremely scarce inhibitory cell markers. Panel a shows colocalization of vGluT and slc6a15/18 (putative glutamatergic neuron markers), panel b shows localization of ChAT and EAAT1 in distinct cell bodies, and panel c is a composite of the markers in panel b and b illustrating distinct expression profiles for the vGluT/slc6a15/18 cells, the ChAT-positive cells, and those expressing EAAT1. Panel d is a composite image of expression profiles for EAAT1, vGluT, slc6a15/18 and the GABAergic neuron marker vGAT (white arrows). Panels e-f depict further magnified regions of panel d (dashed line box) showing coexpression of vGAT (e) and EAAT1 in some, but not all vGAT and EAAT1-positive neurons. Putative glutamatergic (vGluT and slc6a15/18) and GABAergic (vGAT) markers are shown in panels g-i. Scale bars are 50 μM in a-g and for 15 μM in h-i.

An additional exception to exclusive dominance of cholinergic and glutamatergic cells in the DSCN layer is the very scarce presence of putative GABAergic (putative inhibitory) neurons. We identified these by detecting the expression of a transcript whose protein sequence is homologous to those annotated as vesicular inhibitory amino acid transporter (vIAAT) in *Octopus sinensis* (XP 029641102.1) and in *Octopus bimaculoides* (XP 014787249.1). This sequence has also been referred to as a vesicular GABA transporter (vGAT) in a recent publication^27^ and appears to be indicative of GABA expression, suggesting inhibitory activity. This transcript, henceforth referred to as “vGAT”, is one of the extremely scarce indicators for the presence of known inhibitory neuronal markers in the axial nerve cord (Figure 4d-h).

While it is clear that the majority of slc6a15/18-expressing neurons also express vGluT, indicating a glutamatergic identity, we observed that the vGAT transcript also colocalizes with slc6a15/18, but not with vGluT (Figure 4g-i). EAAT1 is also coexpressed in some but not all GABAergic neurons (Figure 4d-f). This suggests that EAAT1 does not exclusively label excitatory neurons as its name suggests, or that these cells may be both excitatory and inhibitory. The initial design of this study also included an additional putative inhibitory neuronal marker - a glycine transporter (GlyT2), to identify inhibitory glycinergic neurons. Despite clear GlyT2 expression in the DSCNs using the chromogenic assay (see supplementary chromogenic images), we did not detect convincing expression of the same molecule using HCR.

#### Arborizing Neuropil Neurons

The third expression profile identified in the axial nerve cord is found in the neuropil - the central portion of the axial nerve cord, surrounded by the cortical layer below the cerebrobrachial tracts. Our study revealed a morphologically striking and ubiquitous cell type throughout the neuropil: EAAT1-positive neurons with long branching dendrites (Figure 4b, d, f and Figure 5a, g). Based on their morphology, these cells appear to be neurons housed entirely within the neuropil, as opposed to neuronal processes projecting from cells whose somata reside in the cortical layer. In addition to the ubiquitous expression of EAAT1 in the neuropil cells, we also identified scarce FLRIamide-positive neuronal somata in the neuropil (not pictured).

**Figure 5:**
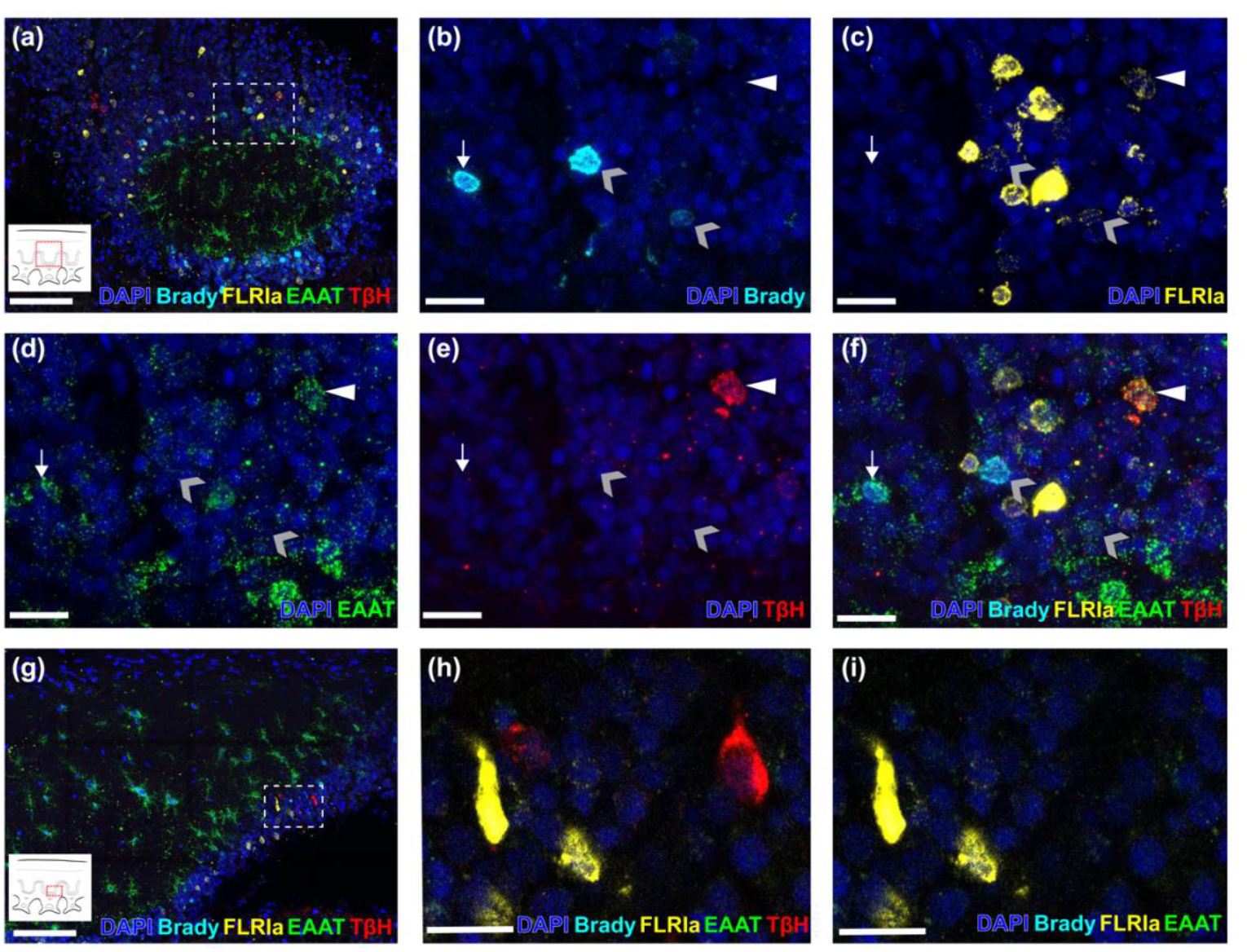
Neuronal markers overlap in unique combinations in scattered cortical neurons. Panel a depicts a composite image of bradykinin-like neuropeptide, FLRIamide, EAAT1, and TβH in the lateral region of a longitudinal section of a ganglion. Panels b-f are further magnified images of the region in the dashed line box in a. The white arrowhead in the upper right of panels b-f depicts a cell expressing FLRIamide, EAAT1, and TβH, but not bradykinin. The gray arrowheads in b-f point to two bradykinin-like neuropeptide -expressing cells that also express FLRIamide. The downward pointing arrow on the left side of panels b-f depicts a bradykinin-like neuropeptide expressing cell that also expresses EAAT1 only. Panel f shows a composite of panels b-e. Panel g depicts a different region of the tissue slice shown in a, exhibiting the same markers. Panels h and i show that unlike the TβH-positive cell in panels a-f, not all TβH cells coexpress FLRIamide and/or EAAT1. Scale bar is 100 μM in a and g and 20μM b-f and h-i.

#### Variability across multiple profiles

As in most biological systems, there appear to be some exceptions to neat classifications of neurons in the octopus arm based on morphology and transcript expression. One exception is the distribution of the EAAT1 transcript, as it can be identified in representatives from all three broad classifications of axial nerve cord neurons. While the EAAT1-positive cells are most striking in the neuropil of the ANC (Figure 4b, d, f and Figure 5a, g), we identified EAAT1 in some of the DSCNs (Figure 4b, c, and f), as well in some SCNs (Figure 5d, f).

Figure 5 depicts an example of multiple instances of neurotransmitter coexpression in scattered cortical neurons. Here we see a single neuron (Figure 5b-f white arrowhead) that appears to be octopaminergic (TβH), peptidergic (FLRIamide), and excitatory (EAAT1). In contrast, Figure 5 panels g-i (as well as expression profiles in Figures 2-3) clearly show that these three markers do not always overlap. We also see peptidergic neurons that express abundant bradykinin-like neuropeptide in combination with relatively low levels of FLRIamide (5b-f gray arrowheads), as well as bradykinin-like neuropeptide-positive cells with EAAT1 (5b-f white arrow pointing down).

One final example of overlapping expression profiles is the colocalization of dopaminergic (TyH) and glutamatergic (vGluT and slc6a15/18) markers (Figure 6). This is consistent with observations of such cells in the developing^27^ and adult^26^ optic lobes. While the majority of glutamatergic neurons are not dopaminergic, we observed that the majority of dopaminergic neurons in the arm did also express both glutamatergic markers (vGluT and slc6a15/18). However, consistent with findings in the optic lobe, there appears to exist a third small population of dopaminergic neurons without vGluT (Figure 6a-b white arrowhead). There did not appear to be any correlation between the size of a dopaminergic neuron and whether or not it was also glutamatergic.

**Figure 6:**
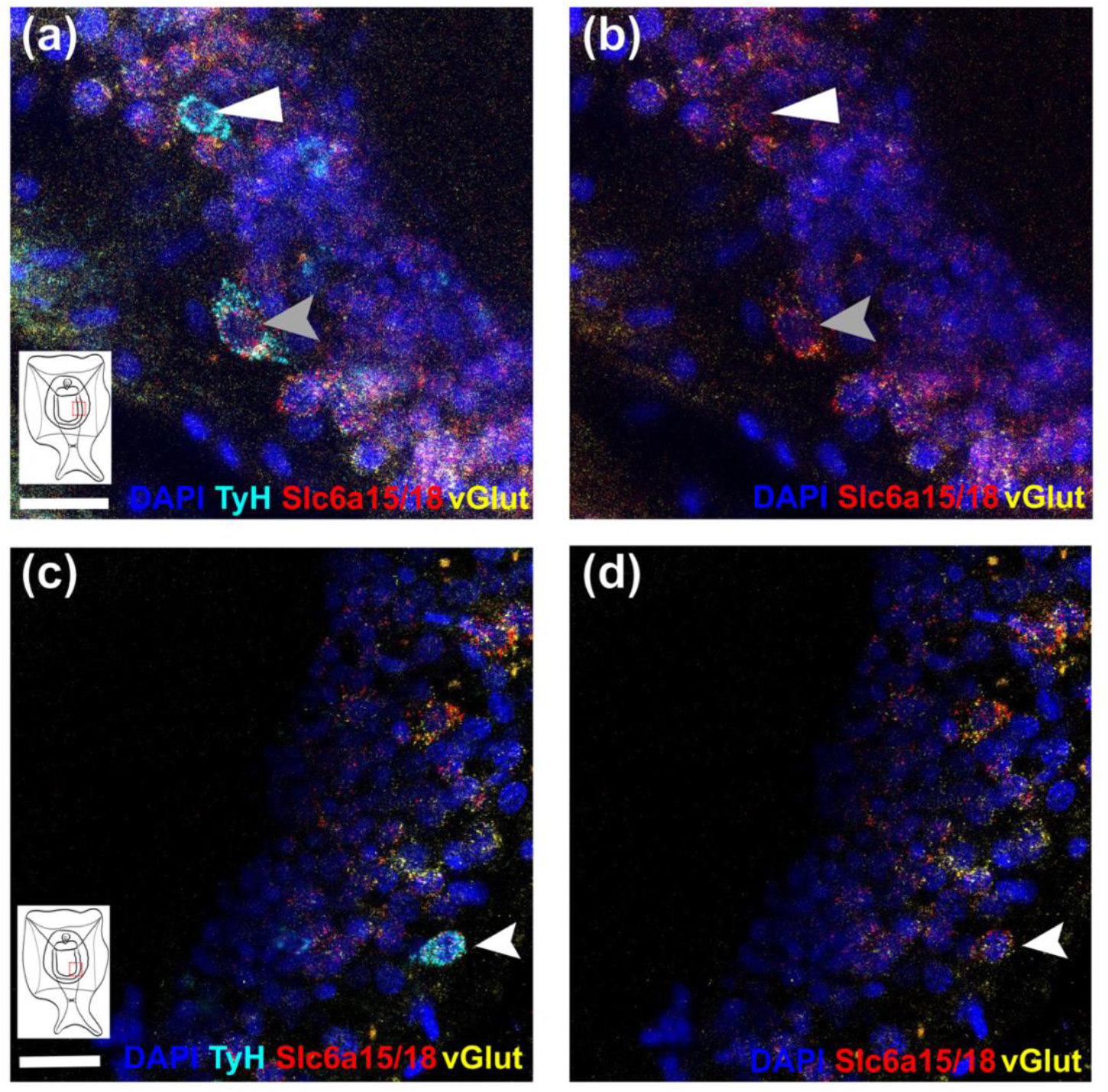
Dopaminergic and glutamatergic neuronal populations overlap in some (but not all) cases. Panels a-d depict glutamatergic neuron markers (vGluT and slc6a15/18) with (a and c) and without (b and d) labeling for dopaminergic neurons (TyH). Panel a depicts two dopaminergic neurons in cyan, the larger of which also expresses vGluT and slc6a15/18 (gray arrows in a and b), and the smaller of which does not (white arrows in a and b). Panels c and d provide an additional example of a relatively small cell expressing both TyH (c) and vGluT and slc6a15/18 (c and d). Scale bars are 20 μM.

## Discusssion

The goal of this study was to begin characterizing the molecular component of the octopus arm connectome. We have successfully produced a three-dimensional map of five different neurotransmitters, neuromodulators, and neuropeptides and compared these maps between sections taken from the arm’s tip and its base. Additionally, we have characterized distinct neuronal populations within the morphologically homogenous and densely packed small neurons of the ANC cortical layer. Finally, we identified interesting sub-populations of neurons with potential cotransmission of multiple molecules.

### 3D Reconstruction and Scattered Cortical Neurons

To date, our grasp of the cytoarchitecture and molecular characteristics within the axial nerve cord has been notably sparse. Our study begins to address this gap by delineating a more comprehensive molecular landscape of individual ganglia. We chose to use the markers that label the scattered cortical neuron (SCN) populations (FLRIamide, bradykinin-like, TβH, TyH, and TpH) to generate our first 3D models because their scattered distribution facilitated visualization, unlike the dense small cortical neurons (DSCNs), whose dense concentration makes it difficult to map individual neurons. The rotating animations provided in this manuscript’s supplementary materials (supplementary animation S1) reveal regions of relatively high as well as lower concentrations of cells labelled by each molecular marker along the proximal-distal axis. From these animations we can also detect and more confidently characterize oral/aboral and ipsilateral/contralateral distribution trends for each marker.

One striking observation we can make using these 3D models is that there are fewer dopaminergic and serotonergic neurons visible in the arm tip than in the base. Although cell count data reveal that these neurons are not entirely absent in the arm tip, they are certainly less abundant and less intensely labeled than those in the arm base. Arm tip ANCs are “newer” tissues developmentally than those in the base, so it is possible that these younger networks are establishing cells that secrete primary neurotransmitters (like acetylcholine and glutamate) first, while neurons containing molecules like serotonin and dopamine that typically play a neuromodulatory role may differentiate and establish connections later. Specifically, serotonin has been proposed to modulate transmission and integration of multiple primary sensory neurons converging on afferent neurons via presynaptic inhibition^6^. One could extrapolate that the necessity of this type of modulation increases with more numerous primary sensory neurons as a tissue grows, thus bolstering the need for increased neuromodulation at the larger arm base than the tip.

Interestingly, the distribution and density of octopamine, the third neuromodulator that we characterized in this study, does not vary dramatically between the tip and the base overall or for any size range except for the small size neurons, which are difficult to visualize in the 3D model. Bradykinin-like neuropeptides cell numbers do not vary significantly from base to tip in any comparison, showing remarkable consistency. Such uniformity along the arm’s proximal-distal axis may suggest a more persistent role in a secretory molecule’s function over time and/or along the length of the cord.

FLRIamide-positive neuron density varies significantly between base and tip (Table S3), but these neurons are exceptionally abundant in the base and the tip, suggesting an important role for FLIRamide peptide throughout the physical length and temporal lifetime of an octopus’s arm. This secretory molecule is also expressed in numerous morphologically diverse neurons in the brain^14,28^, notably in the large efferent neurons of the vertical lobe, which are GABAergic inhibitory neurons that receive excitatory cholinergic input from the amacrine interneurons and project from the memory circuit to premotor centers^14^. FLRIamide-positive neurons are clearly not all GABAergic in the ANC, but it is plausible that their role in modulating motor control is conserved in the arm.

Our study also provides some of the first data about the oral/aboral and lateral molecular architecture of the ANC. Distinct stratification of cell types is easily visualized in along the vertical axis of the cortical layer, with neurons expressing bradykinin-like neuropeptide more abundant toward the oral surface, and octopaminergic neurons located primarily along the medial and aboral or dorsal margins of the walls of the ganglion (Figures 2 and 3 and supplementary animation S1) Curiously, the vertical location of the dopaminergic neuron stratum varies significantly between the arm’s tip and base. We also found differences in distributions in the wall ipsilateral and contralateral to the sucker ganglion/sucker complex. Octopaminergic neurons were significantly more abundant on the same side as the sucker in the arm base, perhaps implying a neuromodulatory role in sucker sensory integration or motor control, and in contrast, FLRIamide-positive cells were more abundant on the contralateral side in the arm tip.

Despite clear stratification along the aboral/oral axis for some markers, neuropeptides and monoamines (as well as primary excitatory neurotransmitters like glutamate and acetylcholine discussed below) were all identified in all six delineated regions of the ANC’s cortical layer (Figure 2i). This information in conjunction with future physiological experiments to characterize the exact functions of each neurotransmitter could reveal functional domains of the arm cord.

Another notable observation from this study is the morphological diversity of previously described “large cells”^1^ of the cortical layer. Although some of the SCN soma are as small as those of the DSCNs, our experiments revealed a gradient of cell body sizes and a variety of shapes. None of the SCN markers was restricted to a single morphology, with soma sizes ranging from 8 to 20 μM for all, and up to 22-25 μM for most (bradykinin-like, FLRIamide, TpH, and TbH), suggesting heterogeneity of identity and function even within these populations.

### Excitatory and Inhibitory Markers

The majority of neurons in the cortical layer are morphologically homogenous cells with small somata. This study revealed that most of these cells fall into two sub-populations of putative excitatory cells: cholinergic and glutamatergic neurons (Figure 4a-c). These sub-populations appear to be completely intermingled and distributed equally and evenly throughout the CL. The abundance of cholinergic neurons is consistent with previous studies^2,4,5^ that implicate acetylcholine in neuromuscular control of the fibers surrounding the ANC. Conversely, we know little about the specific function of the numerous glutamatergic neurons in the ANC CL, but their role as excitatory neurons has been demonstrated in other octopus neural tissues like the vertical lobe^14,29–31^, thus we suggest that the ANC glutamatergic neurons may also be excitatory.

Perhaps one of the most curious findings of our study is the scarcity of presumed inhibitory markers in the ANC. Only approximately one out of every four 50 μM arm slices treated with probes for vesicular GABA transporter (vGAT) showed any labelling, and the maximum number of vGAT-positive neurons we identified in a single slice was three. All vGAT-positive neurons also expressed Scl6a15/18, a marker previously used to identify glutamatergic neurons in octopus^26^ due to its role in transporting glutamate precursors for glutamate synthesis^25^. Cells positive for vGAT and slc6a15/18 do not also express vGluT (Figure 4g-i), suggesting that glutamate produced in slc6a15/18-positive neurons can either be secreted by vGluT-positive (glutamatergic) neurons or decarboxylated and converted to GABA for secretion by vGAT-positive (GABAergic) neurons.

We also noted that a transcript annotated as excitatory amino acid transporter 1 (EAAT1), was coexpressed in some, but not all, GABAergic neurons (Figure 4d-f), glutamatergic neurons (Figure 4a-c), peptidergic neurons (FLRI and bradykinin-like in Figure 5b-f), and octopaminergic neurons (Figure 5d-f). In addition to these myriad cells in the CL, the most striking expression profile of the EAAT1-positive cells is in the arborizing neuropil neurons (Figures 4 and 5) whose broad projections extend in all directions up to 100 μM. The clear abundance of EAAT1 mRNA in these processes indicates that translation may be occurring at the synapse, suggesting a need for rapid and continuous production of protein encoding this potentially important transporter. Unfortunately, the role and identity of the specific molecule(s) transported by EAAT1, as well as the potential excitatory nature of EAAT1-positive neurons, is unconfirmed in octopus without further examination. Therefore, we cannot definitively say that cells labelled by EAAT1 are excitatory.

This study has revealed additional instances of colocalization between molecular markers, like FLRIamide and TbH (indicating an overlap between octopaminergic and peptidergic neurons) and FLRIamide and bradykinin-like (showing potential cotransmission of multiple neuropeptides by the same cell) (Figure 5), but colocalization is not found in the majority of cells for each cell type. This suggests that there may be multiple sub-populations within each cell type, for which molecular diversity is just as (or perhaps more) complex as the morphological and anatomical diversity described above. Another example of this is a partial overlap between dopaminergic and glutamatergic neurons that has also been identified in the inner granular layer of adult^26^ and developing^27^ octopus optic lobes. The ANC cortical layer contains cells that appear to be both glutamatergic (vGluT and slc6a15/18-positive) and dopaminergic (TyH-positive), as well as cells that produce glutamate or dopamine, but not both (Figure 6). Figure 6 also illustrates the fact that each sub-population of glutamatergic and/or dopaminergic neurons is not restricted to a certain size, as markedly different cell sizes are labelled by each marker, even within the same tissue. In fact, the cell depicted in Figure 6a-b expressing vGluT, slc6a15/18, and TyH is decidedly larger than most of the other glutamatergic neurons in the CL, suggesting heterogeneity in the excitatory DSCNs that we had initially overlooked.

### Conclusions

Exploratory, descriptive studies such as these tend to be hypothesis-generating, rather than hypothesis-testing, and as such, our data raise numerous new questions about the structure and function of the octopus arm. Our three-dimensional model provides some of the first molecular maps of the neuronal populations within the octopus arm, revealing previously undescribed anatomical features. Expression patterns along proximal-distal, oral-aboral, and lateral axes may lead to a better understanding of the roles for neuropeptides and neuromodulators in sensory and/or motor control of the arm. We also found an abundance of at least two kinds of putative excitatory neurons (cholinergic and glutamatergic) in the ANC, but saw very few inhibitory neurons, suggesting that inhibitory control may be more centralized than in the periphery. Finally, we highlighted the complexity of these networks by identifying multiple sub-populations of neurons that may produce more than one signaling molecule.

Although we did not include the DSCNs in our three-dimensional analysis of the cortical layer because their high density did not reveal obvious variation in their abundance throughout the cord, it is possible that their distribution is not completely consistent throughout the ganglion. We also did not include analysis of the peripheral neuronal structures of the arm like the sucker ganglion or the intramuscular nerve cords. We plan to characterize these tissues in future studies, to shed more light on the entire arm connectome and help us to better understand arm function.

## Methods

### Animals

Pygmy octopus (*Octopus bocki*) (Figure 1) were purchased from the Sea Dwelling Creatures (Los Angeles, California, USA), in August 2022, March 2023, and June 2023. Octopuses were temporarily housed individually within tanks containing recirculating seawater (1600 L) held at 23.5– 25.5°C and filtered via physical, chemical, and biological filtration. Enrichments such as plastic plants, shells, rocks, PVC pipes, and sand beds and were included in their temporary enclosures to mimic a naturalistic environment. Each octopus (n = 8) was fed 1-2 live grass shrimp (*Paeneus spp*.) per day. Octopuses were monitored daily for general health and euthanized for experiments within two weeks of their arrival. Experiments were conducted between August 2022 and January 2024.

Ethical Note: Octopuses are invertebrates and therefore are excluded from regulatory oversight in the USA; thus, no IACUC protocol was required for this study. However, we adhered to Directive 2010/63/EU and ARRIVE guidelines for characterizing standards of care, humane endpoints, and experimental procedures.

### RNA extraction & cDNA synthesis

To create a preliminary transcriptomes of *Octopus bocki* neural tissues we sequenced RNA from various tissues. We first extracted RNA from *O. bocki* arms, brains (supra and sub-esophageal regions), stellate ganglia, and optic lobes using the Qiagen RNeasy Micro kit (Cat # 74007) and RNeasy Mini kit (Cat # 74104) (See “Animal dissection and tissue preparation” below for euthanasia protocols). cDNA cloning libraries were created using SMARTScribe™ Reverse Transcriptase (Takara/Clontech Cat # 639537) and the Advantage 2 PCR kit (Takara/Clontech Cat # 639207). Libraries were checked for quality and approximate concentration on a 1% agarose gel.

### RNA sequencing

Libraries were prepared with PolyA selection and sequencing was carried out by Azenta Life Sciences (South Plainfield, NJ, USA). *Octopus bocki* RNA samples were quantified using Qubit 2.0 Fluorometer (Life Technologies, Carlsbad, CA, USA) and RNA integrity was verified using the Agilent TapeStation 4200 (Agilent Technologies, Palo Alto, CA, USA). Next, RNA sequencing libraries were prepared using the NEBNext Ultra II RNA Library Prep for Illumina using manufacturer’s instructions (NEB, Ipswich, MA, USA). Briefly, mRNAs were enriched with Oligo(dT) beads and subsequently fragmented for 15 minutes at 94 °C. Next, first strand and second strand cDNA were synthesized. cDNA fragments were end repaired and adenylated at 3’ ends, and universal adapters were ligated to cDNA fragments, followed by index addition and library enrichment by PCR with limited cycles. The sequencing libraries were validated on the Agilent TapeStation (Agilent Technologies, Palo Alto, CA, USA), and quantified by using Qubit 2.0 Fluorometer (Invitrogen, Carlsbad, CA) as well as by quantitative PCR (KAPA Biosystems, Wilmington, MA, USA). The sequencing libraries were multiplexed and clustered onto a flowcell. After clustering, the flowcell was loaded onto the Illumina instrument (HiSeq 4000 or equivalent) according to manufacturer’s instructions. The samples were sequenced using a 2x150bp Paired End (PE) configuration. Image analysis and base calling were conducted by the Control Software (CS). Raw sequence data (.bcl files) generated from the Illumina instrument was converted into fastq files and de-multiplexed using Illumina bcl2fastq 2.20 software. One mismatch was allowed for index sequence identification. Transcripts were assembled from raw paired-end reads using default settings for SeqMan NGen (DNASTAR.17 Lasergene Inc., Madison, WI, USA). These data are available in the Sequence Read Archive (SRA) under accession number PRJNA1050799.

### Sequence homolog identification

*Octopus bocki* transcripts encoding putative neuronal diversity markers (Table 1) were identified using tBLASTn searches of *O. bimaculoides* or *O. sinensis* protein sequences against our *O. bocki* transcriptomes (PRJNA1050799). These *O. bocki* hits were then translated and verified for sequence similarity using BLASTp against the NCBI database. Homologous proteins from selected organisms were included in multiple sequence alignments and gene trees and can be found in supplementary materials. Multiple sequence alignments were created in MEGA 11^32^ using Muscle default parameters (UPGMA cluster method with -2.9 gap open and 0 gap extension penalties) and maximum likelihood trees were generated in MEGA 11 using the Whelan and Goldman model^33^. The percentage of trees in which the associated taxa clustered together was calculated and is shown next to each branch.

### Molecular cloning

Primers (detailed in Table S1) were designed to amplify 300-1000 nucleotides of target sequences and synthesized by IDT (Coralville, Iowa, USA). PCR products were amplified from neuronal tissue cDNA libraries (see above) using TaKaRa LA Taq® DNA Polymerase (Takara Cat # RR002A) PCR products were purified using the MinElute PCR Purification Kit (Qiagen, Cat # 28004), then ligated into vectors and subsequently transformed into *E. coli* cells using the TOPO™ TA Cloning™ Kit for Sequencing with One Shot™ TOP10 Chemically Competent *E. coli* (ThermoFisher, Cat # K4575J10). Following transformation, plasmids were isolated with the QIAprep Spin Miniprep Kit (Qiagen, Cat # 27104). Insertion and orientation of sequences were confirmed by Sanger sequencing using the M13 Forward primer (Elim Bio, Hayward, CA, USA).

### Probe preparation

Plasmids were linearized using Pme1 (NEB Cat # R0560S) or NotI (NEB Cat # 189S) restriction endonuclease (depending on insert orientation). Next, digoxygenin (DIG)-labeled antisense probes were prepared using the Roche DIG labeling kit (Sigma Cat # 11277073910 and T7 (Sigma Cat # 10881767001) or T3 (Sigma Cat # 11031163001) RNA polymerase. RNA robes were purified using RNeasy MinElute Cleanup Kit (Qiagen Cat No./ID: 74204), and 1 μL was visualized on a 2% agarose gel.

### Animal dissection and tissue preparation

Adult *Octopus bocki* were euthanized in cold, isotonic MgCl_2_ solution (330mM MgCl2.6H20 in RODI water) and observed until respiration had ceased for more than 5 minutes. Upon confirming that the octopus had no reflexive movement to a noxious stimulus (arm pinch) it was dissected into individual arms and neural tissues. Initial fixation and sample preparation were modified from established mollusc and octopus colorimetric *in situ* hybridization protocols^28,34,35^ and fixed overnight in 4% paraformaldehyde (PFA) in phosphate-buffered saline (PBS) at 4°C. The following day tissues were rinsed in PBS x3, followed by 100% PTW (1xPBS, 0.1% Tween 20) for 10 minutes. Tissues were then dehydrated stepwise (PTW: MeOH - 3:1, 1:1, 1:3 - 10 minutes each) and stored at -20°C for up to one month. Two days before initiation of HCR protocols, arm tissues were rehydrated (PTW: MeOH - 1:3, 1:1, 3:1 - 10 minutes each) and rinsed in PTW. Next, tissues were post-fixed for 1 hour in 4% PFA in PBS at 4°C to reinforce structural rigidness of the tissue before slicing and rinsed in PBS before being transferred to 30% sucrose in PBS for 1-3 hours (until tissues sunk) for cryoprotection. Intact segments from the base and tip of the arms were embedded in O.C.T. medium (Tissue-Tek® Cat # 4583) and frozen. For HCR only, a minimum of ten serial sections (50 μM thick) were collected from each arm. For HCR and chromogenic *in situ* hybridization 20-30 μM thick sections were also collected on slides in cross and longitudinal section. Sections were allowed to dry for 20-40 minutes before proceeding to the next step. During this time, the area of the slide containing tissue slices was encircled using an ImmEdge Hydrophobic Barrier (PAP) Pen (Vector Laboratories Cat # H-4000) and allowed to dry. Slides were then transferred to lie flat in a humidified chamber, and for the duration of the experiment, solutions were carefully added and removed using pipettes and Kimwipe (Kimberly-Clark Professional™ Cat. # 34120) wicks, so as not to disturb tissue slices. Slides were then washed in PTW for 10 minutes, followed by 10 minutes in 0.3% TritonX 100 in PBS and one final wash in PTW before proceeding with the HCR or chromogenic *in situ* hybridization protocol.

### Chromogenic in situ hybridization

Protocols for chromogenic *in situ* hybridization were adapted from previous studies on *Octopus*^14,28,35^ and *Aplysia*^34,36,34,37^ nervous systems. After fixation, slicing, and initial PTW washes (see “Animal dissection and tissue preparation”), and subsequent permeabilization steps^34^, probes were dissolved in hybridization buffer at 1 μg/μL and enough of this solution was added to slides to cover tissues. Slides were incubated overnight in the humidified chamber at 50°C and washed the following day according to the protocols above. Slides were then incubated overnight at 4°C in 0.1% goat serum with 0.05% alkaline phosphatase-conjugated DIG antibodies (Roche Cat # 11093274910) and washed before development was carried out using detection buffer containing 20 μL/mL of NBT/BCIP. After development was complete (2-24 hours) samples were fixed in 4% PFA in MeOH for 60 minutes, washed in 100% EtOH, and embedded under a cover slip after treatment with methyl salicylate and permount.

### Hybridization Chain Reaction (HCR)

Hybridization chain reaction (HCR) *in situ* hybridization was completed according to Molecular Instruments protocol for “generic sample on a slide” Revision Number 9 from 2023-02-13 (https://files.molecularinstruments.com/MI-Protocol-RNAFISH-GenericSlide-Rev9.pdf) and all probes (see Table S1 for sequences submitted and associated amplifiers) were designed using their website (www.molecularinstruments.com). Probe concentrations of 16 nM were used to increase hybridization yield. Additionally, 100 μg/mL of salmon sperm DNA was added to the amplification buffer for the pre-amplification and amplification steps and the pre-amplification step was extended from 30 minutes to 1 hour to reduce nonspecific background fluorescence. All other reaction steps were completed precisely according to the manufacturer’s protocol. After final washes, tissues were embedded in Fluoromount-G™ Mounting Medium, with DAPI (Thermo Fisher Cat # 00-4959-52).

### Imaging

Fluorescent images were acquired using a Leica Stellaris DMI8 Confocal microscope (HC PL APO CS2 40x/1.30 OIL objective) and Las X 4.6.0.27096 software. Diode 405 was used to acquire DAPI images and WLL (85% of max power) was used to acquire all other channels. All images were taken in a 1024x1024 pixel format with a pixel dwell time of 0.6 μS. Bidirectional scanning was used for efficiency. Images were imported into FIJI^38^ and channels were merged before concatenating all sequential stacks (where applicable).

### Cell Counts

Cell counts were acquired by measuring the soma diameter of each individual cell in maximum intensity projections from selected slices in tip (“Tip 1”, “Tip 4”, “Tip 7”) and base (“Base 3”, “Base 6”, “Base 9”) using Las X 4.6.0.27096 software. Counts were aggregated across replicates. This aggregation was performed to sum counts across all sizes and locations for specified replicates. Generalized linear mixed models (GLMMs) were fitted to the cell count data to estimate the fixed effects of cell location (ipsilateral vs contralateral and oral vs medial vs aboral) and cell size while treating variation among individual slices of arm tissue as random effects. Each model was fitted using maximum likelihood estimation, with the ‘glmer’ function from the ‘lme4’ R package. The cell counts were modeled using a Poisson distribution with a logarithmic link function. Model selection and validation involved examining the significance of fixed effects, assessing model fit using Akaike’s Information Criterion (AIC), and ensuring the absence of overdispersion. Negative binomial (NB) models were also examined to account for overdispersion detected in some of the comparisons but comparing AIC between NB and Poisson models did not clearly favor the choice of one distribution over the other. Fitting NB models resulted in nearly identical fixed effect estimates as Poisson, so the Poisson distribution was selected since overdispersion was not always present. The “cell location” variable was examined in a variety of ways by fitting multiple models. Consistent in each model was the fixed effect “type”, corresponding to either the base of the arm or the tip. The following comparisons were subsequently made (see Tables S3-6 for values): 1) Tip vs base: total number of cells per cell marker, 2) Tip vs base: number of cells of each size classification per cell marker, 3) Oral/Aboral axis comparison: total number of cells per region (oral, medial, or aboral) in all preparations (tip and base for all markers), and 4) Lateral comparison: total number of cells per region (ipsilateral vs contralateral to sucker) in all preparations (tip and base for all markers). Once model parameters were estimated, each comparison was evaluated, generating a p-value. Post-hoc correction was applied to p-values to account for multiple comparisons where appropriate using hard Bonferroni adjustments. Specifically, adjusted p-values were calculated by multiplying the original p-values by the number of comparisons conducted within a data set (three for the oral/aboral axis comparison and four for the tip vs base cell size comparison).

### 3D reconstructions

Z-stack images of each sequential arm slice were acquired at 1 μM depth intervals (0.28x0.28x1.0 μM voxel size) in seven channels. Due to tissue shrinkage during the HCR treatment, this amounted to 22-31 images per tissue slice. For the arm base slices, we acquired 5x5 (25 total) tiled images for each section of every z-stack in order to ensure that the entire axial nerve cord cross section was captured. For the smaller arm tip slices we acquired 3x3 (9 total) tiled images for each section of the z-stacks. After concatenation and manual alignment in FIJI^38^, channels were split and despeckled using a 3 pixel median filter and individual files for each channel were imported into Amira^39^ for 3D reconstruction and animation.

## Supporting information

Animation S1

## References

1. Young, J. Z. The Anatomy of the Nervous System of Octopus Vulgaris. (Clarendon Press, Oxford, 1971).

2. Olson, C. S. & Ragsdale, C. W. Toward an Understanding of Octopus Arm Motor Control. Integr Comp Biol 63, 1277–1284 (2023).

3. Young, J. Z. The number and sizes of nerve cells in Octopus. Proc. Zool. Soc. Lond. 140, 229–254 (1963).

4. Matzner, H., Gutfreund, Y. & Hochner, B. Neuromuscular System of the Flexible Arm of the Octopus: Physiological Characterization. J Neurophysiol 83, 1315–1328 (2000).

5. Sakaue, Y. et al. Immunohistochemical localization of two types of choline acetyltransferase in neurons and sensory cells of the octopus arm. Brain Struct Funct 219, (2013).

6. Bellier, J.-P. et al. Immunohistochemical and biochemical evidence for the presence of serotonin-containing neurons and nerve fibers in the octopus arm. Brain Struct Funct 222, 3043–3061 (2017).

7. Miller, M. W. GABA as a Neurotransmitter in Gastropod Molluscs. Biol Bull 236, 144–156 (2019).

8. Lunt, G. G. GABA and GABA receptors in invertebrates. Seminars in Neuroscience 3, 251–258 (1991).

9. Bacqué-Cazenave, J. et al. Serotonin in Animal Cognition and Behavior. Int J Mol Sci 21, (2020).

10. Barron, A., Søvik, E. & Cornish, J. The Roles of Dopamine and Related Compounds in Reward-Seeking Behavior Across Animal Phyla. Front Behav Neurosci 4, (2010).

11. Brembs, B., Lorenzetti, F. D., Reyes, F. D., Baxter, D. A. & Byrne, J. H. Operant Reward Learning in Aplysia: Neuronal Correlates and Mechanisms. Science (1979) 296, 1706–1709 (2002).

12. Lechner, H. A., Baxter, D. A. & Byrne, J. H. Classical Conditioning of Feeding in Aplysia: II. Neurophysiological Correlates. The Journal of Neuroscience 20, 3377 (2000).

13. Lechner, H. A., Baxter, D. A. & Byrne, J. H. Classical Conditioning of Feeding in Aplysia: I. Behavioral Analysis. The Journal of Neuroscience 20, 3369 (2000).

14. Stern-Mentch, N., Bostwick, G. W., Belenky, M., Moroz, L. & Hochner, B. Neurotransmission and neuromodulation systems in the learning and memory network of Octopus vulgaris. J Morphol 283, 557–584 (2022).

15. Shomrat, T., Feinstein, N., Klein, M. & Hochner, B. Serotonin is a facilitatory neuromodulator of synaptic transmission and reinforces long-term potentiation induction in the vertical lobe of Octopus vulgaris. Neuroscience 169, 52–64 (2010).

16. Messenger, J. B. Neurotransmitters of cephalopods. Invertebr. Neurosci. 2, 95–114 (1996).

17. Turchetti-Maia, A. et al. A Novel NO-Dependent ‘Molecular-Memory-Switch’ Mediates Presynaptic Expression and Postsynaptic Maintenance of LTP in the Octopus Brain. (2018). doi:10.1101/491340.

18. Selcho, M., Pauls, D., el Jundi, B., Stocker, R. F. & Thum, A. S. The Role of octopamine and tyramine in Drosophila larval locomotion. Journal of Comparative Neurology 520, 3764–3785 (2012).

19. Andrews, J. C. et al. Octopamine Neuromodulation Regulates Gr32a-Linked Aggression and Courtship Pathways in Drosophila Males. PLoS Genet 10, e1004356. (2014).

20. Price, D. A. & Greenberg, M. J. The Hunting of the FaRPs: The Distribution of FMRFamide-Related Peptides. Biol Bull 177, 198–205 (1989).

21. Di Cosmo, A. & Di Cristo, C. Neuropeptidergic control of the optic gland of Octopus vulgaris: FMRF-amide and GnRH immunoreactivity. Journal of Comparative Neurology 398, 1–12 (1998).

22. Zheng, L., Cao, H., Qiu, J. & Chi, C. Inhibitory Effect of FMRFamide on NO Production During Immune Defense in Sepiella japonica. Front Immunol 13, (2022).

23. Walker, R. J., Papaioannou, S. & Holden-Dye, L. A review of FMRFamide- and RFamide-like peptides in metazoa. Invertebrate Neuroscience 9, 111–153 (2009).

24. Kobayashi, M. & Muneoka, Y. Functions, Receptors, and Mechanisms of the FMRFamide-Related Peptides. Biological Bulletin 177, 206–209 (1989).

25. Bröer, A. et al. The orphan transporter v7-3 (slc6a15) is a Na+-dependent neutral amino acid transporter (B0AT2). Biochemical Journal 393, 421–430 (2005).

26. Songco-Casey, J. O. et al. Cell types and molecular architecture of the octopus visual system. Current Biology 32, 5031–5044 (2022).

27. Styfhals, R. et al. Cell type diversity in a developing octopus brain. Nature Communications 2022 13:1 13, 1–17 (2022).

28. Winters, G. C. Molecular Mapping of the Octopus Brain. (Gainesville, FL, 2018).

29. Hochner, B., Shomrat, T. & Fiorito, G. The Octopus: A Model for a Comparative Analysis of the Evolution of Learning and Memory Mechanisms. Biol Bull 210, 308–317 (2006).

30. Hochner, B., Brown, E. R., Langella, M., Shomrat, T. & Fiorito, G. A Learning and Memory Area in the Octopus Brain Manifests a Vertebrate-Like Long-Term Potentiation. J Neurophysiol 90, 3547–3554 (2003).

31. Shomrat, T. et al. Alternative Sites of Synaptic Plasticity in Two Homologous Fan-out Fan-in Learning and Memory Networks. Current Biology 21, 1773–1782 (2011).

32. Tamura, K., Stecher, G. & Kumar, S. MEGA11: Molecular Evolutionary Genetics Analysis Version 11. Mol Biol Evol 38, 3022–3027 (2021).

33. Whelan, S. & Goldman, N. A General Empirical Model of Protein Evolution Derived from Multiple Protein Families Using a Maximum-Likelihood Approach. Mol Biol Evol 18, 691–699 (2001).

34. Jezzini, S. H., Bodnarova, M. & Moroz, L. L. Two-color in situ hybridization in the CNS of Aplysia californica. J Neurosci Methods 149, 15–25 (2005).

35. Winters, G. C., Polese, G., Di Cosmo, A. & Moroz, L. L. Mapping of neuropeptide Y expression in Octopus brains. J Morphol 281, 790–801 (2020).

36. Antonov, I., Ha, T., Antonova, I., Moroz, L. L. & Hawkins, R. D. Role of Nitric Oxide in Classical Conditioning of Siphon Withdrawal in Aplysia. The Journal of Neuroscience 27, 10993 (2007).

37. Moroz, L. L. et al. Neuronal Transcriptome of Aplysia: Neuronal Compartments and Circuitry. Cell 127, 1453–1467 (2006).

38. Schindelin, J. et al. Fiji: an open-source platform for biological-image analysis. Nat Methods 9, 676–682 (2012).

39. Stalling, D., Westerhoff, M. & Hege, H.-C. Amira: A Highly Interactive System for Visual Data Analysis. in Visualization Handbook (eds. Hansen, C. D. & Johnson, C. R. 749–767 (Butterworth-Heinemann, Burlington, 2005).

