## Supplemental Materials for "Three-Dimensional Molecular Atlas of Octopus Arm Neuroanatomy Highlights Spatial and Functional Complexity"

Table S1.

| Cell Type (Neurotransmitt | Transcript name | HCR Amplifier used | Sequence submitted for HCR probe (FASTA format) | Fluorophore 1 used | Fluorophore 2 used | Shorthand HCR probe | Clone number for chromogenic | Cloned sequence (probe template) (***indicates the entire sequence was not read) | Primer 1 | Primer 2 |
|---|---|---|---|---|---|---|---|---|---|---|
| Cholinergic cells | Choline acetyltransferase (ChAT) | B1 | >Obo.ChAT.B1 [illegible] | 647 | 514 | ChAT B1 | 32 | [illegible] | CGTTCCAAGC GAGCAGTACG | TTGTGTTTTCAT GGCCGCCC |
| Dopaminergic cells | Tyrosine Hydroxylase (TyH) | B4 | >Obo.TyH.B4 [illegible] | 647 | 546 | TyH B4 | 42 | [illegible] | ATCTCGCCT GGAAGTGG | CAGGCCAA ATTCCACAG |
| Excitatory cell (Maybe | Excitatory Amino Acid Transporter 1 | B4 | >Obo.EAAT1.B4 [illegible] | 546 | 647 | EAAT1 B4 | 39 | [illegible] | ATGAAAAGA GCGGCCAC | ACAAGTGTA CGTACCCAA |
| GABAergic/Inhibitory | vesicular GABA Transporter/ | B1 | >Obo.vIAAT/vGAT.B1 [illegible] | 647 | 514 | vGAT B1 | 6 | [illegible] | CCTGTTATG GACGCAGT | GAGGCGAC AGTTCTCGT |
| Glutamatergic cells | vesicular Glutamate Transporter | B3 | >Obo.VGluT1.B3 [illegible] | 488 | 543 | VGluT B3 | 9 | [illegible] | CGCCATGGAA GAAATTCTTTA | GCGATGAGAA ACACGTGTTC |

| | | | | | |
|---|---|---|---|---|---|
| Serotonergic cells | Peptidergic cells | Peptidergic cells | Octopaminergic cells | Glycinergic Cell | Glutamatergic or GABAergic cell |
| Tryptophan Hydroxylase (TpH) | Bradykinin-like peptide precursor | FxRIamide peptide precursor | Tyramine β hydroxylase (TβH) | Glycine Transporter (GlyT) | Solute Carrier Family 6 Member 15/18 |
| B3 | B1 | B2 | B5 | B4 | B5 |
| >Obo.TpH.B3<br>ATGGATGCCCGTTCAATTAGCTTCGAAAGAGCAGAGGTGATGAGGAGACAACAATCAACGGATTTTGACGATCACATTGCCGTTGTACCAGAGGAAACGGAAAATGGAAGAGGTGTTCAAGTCGTGAAAAAGTGGGCTCTATATTGCTGTCACTGAAGAATCAAGTTGGCTGTTTGTCAAGTGCTTTRAAAGTATTTCAGGACACAAAGTCAATGTCACACACATCGAATCGCGCAGATCGCAGATGGAAGACAGTGAATATGACATTTTCATCAACGTAGAGTCMAGTAAACTGGCTATAGAAGACCTTCTCGAAAAGCTTCAACTCCAAATCCTGATGATGTAGGAAATGTGAAGGCATTTCCCACCTGTCCTGAAGAACCAGATGATGATATGTGTGGTTCCTCGCAAAATAGCGGAATGGACAAATTTGCCAACCGTGTACTAATGTATGGTCGCGAACTAGATGCCGATCACCCTGGCTTCAAGGATCCAGTATATAGACAACGTAGGATAAAATTTGCCGAAATTGCTTATAATTATAAACATGGCCAACCAATACCAAGAGTTGAATACACAGCAGAAGAAGTAAAACACTTGGGGCACTGTCTTCAGAGAACTAAACAAGCTTTACCCTACCTACTCCTGCAAGCAGCATCTGAAGAACTTACCATTACTAATGAAATATTGCGGATACAGAGAGAACAATATTCCACAACTCGAGGATGTTTCAAAATTTCTCAAAGAGAGAACCGGGTTTACTGTTCGCCCTGTGGCGGGATATTTGTCTTCCAGAGACTTTCTAGCCGGGCTAGCATTCCGTGTGTTTCATTGCACCCAATATATTCGCCCATAGTTCCGATCCTTTCTACACACCAGAACCAGACTGTTGCCACGAATTACTGGGACATGTACCATTGTTTGCTGATCCTGATTTTGCTCATTTTTCTCATGAGTTAGGGTTAGCCTCCCTGGGGGCTTCAGATTTCGAAGTTGCCAAATTAGCCAGTTGCTATTTCTTCACAATTGAATTCGGATTGTGCCGAGAGAATGGTCAGCTCAGAGTGTATGGAGCTGGTCTCTTGTCTTCGATTGGCGAACTCAAGCATGTTCTCACAGACGCAGCCATTAAAAAGCCGTTCGATCCTGACAAAGTTATTGAGCAAGAATGTCAGGTTTCCACATACCAGGATGCCTATTTTGTCAGTGAATCTTTCAAAGAGGCCCAAAATCAAATCAGGGAATTTGCTTCGAACATCAAACGACCCTTCACTGTACGCTATGATCCTTATCACAGTCGGTATGGTTCTCGAAAGTTTGTCAACCATTAGCCAAGTAGTCAGTTTTGTTCGAGCCGATCTTTGTCTCATCTCTGATGCCATCAACAAACTCCACCAGTCGAAGATGAAGAAAACGAGACGAGCACGAACAAGGTTACTCAATGACAAA | >Obo.Bradykinin-like.B1<br>TTTCACTTGCAACTGTGACCCCACTCAACCGTTACATATATAGGCCTGCGATACGGCATCGACACTTCAGTTGAAAGATTCACGCCGGCAACTGCAAAACTCAGCGACTTCCGATAACAACGTGTTTCTACTAAAGAAACAAAAAACGATGGGAATAAATATAATCATGTGCTCTGTCTCGTCGCCGTGTTTGCTTCAAGCGTATATTCATTGCCGAAGCGGAATGACCATGTTATTACTCTGCGTTCGTTACGACAAACCGGCCTCAGTGATTCTGACAGTCGAGCATTACTTCAAGCATACATTCTCGGAAAGCTATCCAACGTAGACATTGCTGGTAACAAGGAAGCAGACAATTCAGATTATACTACCATTAAGAGGAAAGCATTTGGAGACCTATGGGTTACCTTCCTTTGATAACCATGGCGGTAGTGGCGCTAGCAGCAGCAACGATAACTCTCCCAGTGGAGGTACAGGATCTGCTGTATTCAGATATGGTTAAGTGACTCTGAGaAAAAAAAAATCAAATATATTATGAGGAGAGAATTAATGAGTATCAGTATGAACTAAGTAAAGACCTAACGAATTGGTTATTGTCTTCAGCGAGGTTGATTCAATTAAAATAGTATTCTTCTTC | >Obo.FLRI.B2<br>TTGTTAAAGAAAATGGCCGGTCTTTGGAGAATCGTTCTTTTAGCAACAATTACTTTGATGTCTGTGACATCGCAATGGGTTATCTATGTCCAGGCTGAAGAAGCTTCTAACAATGACGAATTGGTGAACAGTGCGAATCAAGCTGACCCTGAAATGGATAGCGAAAATGATCTGAAACGGGCAAATACATTTCTTCGAATAGGTAAAGCAAACGGAGTTTACGTTTAGCTCGAAGCCCTAGTTCTTTTCTGAGAATTGGTCGTCGTAACCAATTGCGCTTTATAAGAATCGGTAAAGCTCCATCAAGCATGTTTCTAAGAATTGGCAAAAGAGTGGATGATAGTGAAAATGATGATCTAAATGAATACGATGCCGATAAAATGTCAGGGCGTGATATCAGAGCCAGTCCATCATCGTTTCTTAGAATTGGAAAATCAGGAAATGAAGTAGAAGATGAAATTGATGACACGGATGAAATTGCTAAACGCGTTAACGCATTTCTACGCATCGGTCGACAAAATGATCCATCGTCGTTCCTGAGAATTGGTAAAAGTTTGAATAATGAAGATCTCTCTAAGGACAAACGTACCAACGCTTTCCTCAGAATCGGAAAAATACCTACCTCATCTTTCATTCGCTTAGGTCGTGGACCTTTCACAGAAGATAATGGCATCAACACTCGTGGATTTCGTGGTCCTACACATGGCTTTTTAAGGATTGGTAAACGCGCAGCGAACACCGACGGCAGTCATACTGATATTCAGTGATCTGAACGTCAAA | >Obo.TBH.B5<br>TAGCTGCTCCCAAGGTCACAGGTCGTCAGCATTATAAATATAATGTTTATCTGGACAGCCAGCGTCGTTTCCATTTATGGTGGGATGTTGACTATAAACGAAATGTGGTAAACTTCCGCCTTTTGGCGGTCGTGACGTTAGATGAATGGTTCGGTGTTGGTTACTCAGCGTATGGAAATCTTCAAATGCCGATTTCTTCGTTCACTGGGTCGACGAAAACTACGTCCAGCACTGTAAGGATGCATGGACGGACAGCAAAGGTATACTCCACATTGATAATCAACAAGATTACAAAATACTGTCTGGTAAGTTGAACAAGCATGCATTGGTCATTGAATATGAAAGACCTTTCGACACCTGTGACAATATGATTTTCCTTTTGATAACGGAACCCTTCACGTGGTATTCTTCATCGGTGATAACACGTTTCACTATCCGGAAGGGGTCAATGTGACTTCCTTATCGCCGGGACTTCAGCGTGTGCTTCTTCTGGAAGCCGATGTTCCGGTCCCTAGACATCCTAAAGACACCTGGACTTTTTCGGTTCTAGCTCCCGGCGTTACAATACCAGCAAAAGAAACGACAAACTGGTGGTACACAACAAAACTTCCAGCACTTAACGAACGACAGCATATAATCCAGTTTGAAAGCGCCATAACAAAAGGTAATAAGAATTTTGTTCATCACATCGAAGTGTTCCTCTGTGAGTGGAAACGTGGAATGAAAATACCCTATTTTAACGGTCCAAGTATGGTAGGCGAGAAACCTAAGGAACTAGTGTTTGTCGACATGTCATAGGAGCATGGGCTATGGGTGCGTCTGCCTTACATTTACCAGCAGANGCAGGCTGTGCAATTGGAGGCAACATCTCACGTGATATATTACTGGAAGTTCATTATATAATATCCTGAAATGAAACACGGAGTTAANGACACTTCAGGAATAAGTNCTATGTTACNCCTACACGAAGGAAGTACGATTCTGGAGTCATGGAAATTGGNTTGG | >Obo.GlyT.B4<br>aatgtttataaggatgcacttattgtccctatgatcaattgtgctacaagtatcttcgcaggtttgtcatcttctcagttcttggatttatggctttgaaactggtgttAGTGTTGACAACGTCGTCCGACAAGGTCCCGGTCTGACTTTCGTTGCTTATCCAGAAGCCCTTTCTCGTTTACCGATATCTTCGTTGTGGCCGTTTTATTCTTCCTATGCTTTTATAATCGGTCTGGACAGTCAGTTCGGAATGTTCGAGACAATGCTCAGTGCCTTCATGGATGAATATCCAGAATACCTCAGAGGCAAAAAAACCCTAATTGCGCCATTGCCTGCTTTGTGGAACTCCTCTCGGCCTGCCCTGCATCACACAGGGCGGAATCTACGTATTACAAATCATGGATTGGTACTGTGCAAGTTTTTCCCTTATGTTAATCTCTTGGCAGAGTGTGTTGTTATAGCATGGATTTACGGGGTCGATCGGTTCTACAAGGACATTGAACTTATGATTGGTTATCAGCCGTGTCGATGGTGGAAATTTCCTGGTGTTTTATCACGCCTGCGGTTATCTTGTTCATTTGGCTTTTCAGCGTTTCCACATTAGGACCTGTAACATACGGTGAAATCAGTACCCACCATGGGCTATAAGATTTGGCTGGATTCTTGGCTAGTTTCATTGGTCCCAATACCTTTAGTGATGATATATTCGATTTACAGAGCTGAGGGAACATTCATGGAAAGAGTAAAGTCGTTAATTAAACCAGCTCCGAACTGGGGTCCCGTCCTACCGGAAAACCGCAAACTTTATTTGGCTTCTCTCTAAATGGTAAAATGGAAAAGTATGTAATACAGCAACGTGGAGAAGAACAAGAAAGAAACCAAAGCGGAACACTGAAAATGGAATCTCAAATCTGATTGGTCAATGAGATCTGTTAATATTGAAGACGTCAGAGTGAGACTCGTAATCGAAAATGAGGACAAGATACGAAAGAAAACTATAAATAACTGGAAATTACGTTAAAATAGATACGGTGTATATTATTAATAGTCCTGGAGTAGAGAAACAGCTGTCGGGTTTCCGAG | >Obo.Slc6a15.B5<br>ATGGAACGTATTGAAATGAATCATTTGAATATACCGAGCCCACATTCGGACGAGAATGGTGATGGAAATAGCAACACTTTGTATGGAAGTACAATGACGCTGGGCGCAAGCTCTCCAGGAGTTGGCGCATCCACAGCTAATTTACTTAAGATTGATTTGGAGAAAAAAATCGATCGACATACGGGTGAAGAAAGAGAAACCTGGGATAATCGTGTCCAATTCATACTCTCATTGGCTGGTTTTGCTGTAGGTCTCGGAAATGTTTGGCGTTTCCCATATTTGACTCAGAAAAATGGAGGAGGGGCATTTCTTATACCATACGCTGTAATGCTGTTTGTGGAAGGAATCCCTATATTCTATTTGGAGTTAGCTATAGGTCAAAGATTACGCAAAGGTGCTATTGGTTGCTGGAATCAAGTATCACCTTATTTAGGTGGTCTTGGTATAGCTGCCGCAGTTGTCTCCTTCAACGTGGCGCTATATTACAACACAATCATGGCATGGTGCATGTACTATCTAGTCGAAAGTTTTCAGAGTCCTCTACCTTGGTCTTCCTGTCCATCTGAAACAGTGGGTAATATTACGACATTTAACCAAGAATGTGAAAAAAGTGGTTCGACAACCTACTTCTGGTTTCGTGTCGCTCTCAACAGTGCTCCGGCGATCGATTCAACACCTATAATAAGCTGGAAGATATCAGCCGCATTAATATGCGCTTGGTTATTGGTTTTCTGTTGCATGATCAAAGGCATCAAATCTTCTGGAAAGGTTGTCTATGTTACAGCAACTTTCCCATATTTTGTCCTGGTCATTTTTTCTTTCGTGGAGTAACTTTACATGGATTTGAGGAAGGTGTAAATACTTATTATTCCAAAGTGGGACAAGTTATTCGAACCTGCTGTTTGGTTAGACGCAGCAACTCAAATATTCTATCATTTGGCTTAGCATTCGGATGCTTGGTAGCTTTGGCTTCATATAATCCTGTCCGCAGTAACTTTCTAAGTGAAGCCCTTATGGTCAGTGTCGGCGACTCTTCACTTCCTTATTGACTGCCACAGTGGTTTTCTCAGTTCTCGGTTTCAAAGCAACAATGGCATACGAGAAATGCCTTGAGAAATTTGGCAGCACAAATCACACAAAGAATCCTAACATCACAGCAGAAAAAGAATGTGATTTTGAAAAACTAATGACAGAGTCAGCCTCTGGGACCGGACTAGCATTCATTGCCTTCACAGAAGCAATTGATCAGTTTCCTGTTGCGCCATTGTGGGCCGTTCTCTTCTTTCTCATGCTGCTCACACTTGGATTGGATACAATGTTTGGTACTCTTGAAGGTGCCATTACATCTATAAATGATATGTTATTGTTTCCTGCAATCAGAAAAGAAGTGTTTGTGGCATTGTATGTTTCGTATCAATGTTAATATCATTTTGCTTTGCAACGTCAACTGGTCCATATGTTTTGACCTGTTCGACATGTATGTGCAAATATCCCTCTCTAGTCATTGCCTCATTGAAGTCATTGCAGTTCATACATATACGGACTGAAACAGTTTAGTGATGACGTAGAAATGATGGTTGGAATACGTCCCAGTTACTTTTGGATGTTTATTTGGCGTTATCTAGCACCTCTAGTTATGATAGTCATTTTCGTTGCTAGTTTGTTCGAAATTTAGAGAATGGATCTATTATATGGCATGGGATGCAGACAAGGGTCACACTGTAATGCAACCCGACCATGGTGGGCGCAATTTATAGCTGCACTTCTGATCCTGTCCTCAGTTGCATGGATTCCATTTGTAGCCATTACACGTTACTTCAACTTAGTTAAATGGGAACCTGAGACGCCAGCTTCTTTCCAGCCGAAGAACTACGGGAAGAACGAAACATCAAGCCTCATCAAGCTACATTTATTGAAAAATATTTCTTTGGATTTAAAGGC |
| 543 | 514 | 594 | 488 | 546 | 594 |
| 488 | 647 | 514 | 594 | 647 | 488 |
| TpH B3 | Bradykinin-like B1 | FxRI B2 | TβH B5 | GlyT B4 | slc6a15/18 B5 |
| 51 | 2 | 16 | 25 | 45 | NA |
| ACACTTGGGGCACTGTCTTCAGAGAACTAAACAAGCTTTACCCTACCCACTCCTGCAAGCAGCATCTGAAGAACTTACCATTACTAATGAAATATTGCGGATACAGAGAGAGCAGTATTCCACAACTCGAGGATGTTTCAAAATTTCTCAAAGAGAGAACCGGGTTTACTGTTCGCCCTGTGGCGGGATATTTGTCTTCCAGAGACTTTCTAGCCGGGCTAGCATTCCGTGTGTTTCATTGCACCCAATATATTCGCCCATAGTTCCGATCCTTTCTACACACCAGAACCAGACTGTTGCCACGAATTACTGGGACATGTACCATTGTTTGCTGATCCTGATTTTGCTCATTTTTCTCATGAGTTAGGGTTAGCCTCCCTGGGGGCTTCAGATTTCGAAGTTGCCAAATTAGCCAGTTGCTATTTCTTCACAATTGAATTCGGATTGTGCCGAGAGAATGGTCAGCTCAGAGTGTATGGAGCTGGTCTCTTGTCTTCGATTGGCGAACTCAAGCATGTTCTCACAGACGCAGCCA | CATGTGCTCTGTCTCGTTGCCGTGTTTGCTTCAAGCGTATATTCATTGCCGAAGCGGAATGACCATGTTATTACTCTGCGTTCGTTACGACAAACCGGCCTCAGTGATTCTGACAGTCGAGCATTACTTCAAGCATACATTCTCGGAAAGCTATCCAACGTAGACATTGCTGGTAACAAGGAAGCAGACAATTCAGATTATACTACCATTAAGAGGAAAGCATTTGGAGACCTATGGGTTACCTTCCTTTGATAACCATGGCGGTAGTGGCGCTAGCAGCAGCAACGATAACTCTCCCAGTGGAGGCA | CCGGTCTTTGGAGAATCGTTCTTTTAGCAACAATTACTTTGATGTCTGTGACATCGCAATGGGTTATCTATGTCCAGGCTGAAGAAGCTTCTAACAATGACGAATTGGTGAACAGTGCGAATCAAGCTGACCCTGAAATGGATAGCGAAAATGATCTGAAACGGGCAAATACATTTCTTCGAATAGGTAAAGCAAACGGAGTTTACGTTTAGCTCGAAGCCCTAGTTCTTTTCTGAGAATTGGTCGTCGTAACCAATTGCGCTTTATAAGAATCGGTAAAGCTCCATCAAGCATGTTTCTAAGAATTGGCAAAAGAGTGGATGATAGTGAAAATGATGATCTAAATGAATACGATGCCGATAAAATGTCAGGGCGTGATATCAGAGCCAGTCCATCATCGTTTCTTAGAATTGGAAAATCAGGAAATGAAGTAGAAGATGAAATTGATGACACGGATGAAATTGCTAAACGCGTTAACGCATTTCTACGCATCGGTCGACAAAATGATCCATCGTCGTTCCTGAGAATTGGTAAAAGTTTGAATAATGAAGATCTCTCTAAGGACAAACGTACCAACGCTTTCCTCAGAATCGGAAAAATACCTACCTCATCTTTCATTCGCTTAGGTCGTGGACCTTTCACAGAAGATAATGGCATCAACACTCGTGGATTTCGTGGTCCTACACATGGCTTTTTAAGGATTGGTAAACGCGCAGCGAACACCGACGGCAGTCATACTGATATTCAGTGATCTGAACGTCAAATGATTCTACAACTAATTAATAAAAGACACCGGTTAAAGGGAACAATTAGAGTTAAAAGAAAAGATTACCATTTGGAAATGATTCTGAAACGAATAAGGACTATCACCGACGCAA | AGCTGCTCCCAAGGTCACAGGTCGTCAGCATTATAAATATAATGTTTATCTGGACAGCCAGCGTCGTTTCCATTTATGGTGGGATGTTGACTATAAACGAAATGTGGTAAACTTCCGCCTTTTGGCGGTCGTGACGTTAGATGAATGGTTCGGTGTTGGTTACTCAGCGTATGGAAATCTTCAAATGCCGATTTCTTCGTTCACTGGGTCGACGAAAACTACGTCCAGCACTGTAAGGATGCATGGACGGACAGCAAAGGTATACTCCACATTGATAATCAACAAGATTACAAAATACTGTCTGGTAAGTTGAACAAGCATGCATTGGTCATTGAATATGAAAGACCTTTCGACACCTGTGACAATATGATTTTCCTTTTGATAACGGAACCCTTCACGTGGTATTCTTCATCGGTGATAACACGTTTCACTATCCGGAAGGGGTCAATGTGACTTCCTTATCGCCGGGACTTCAGCGTGTGCTTCTTCTGGAAGCCGATGTTCCGGTCCCTAGACATCCTAAAGACACCTGGACTTTTTCGGTTCTAGCTCCCGGCGTTACAATACCAGCAAAAGAAACGACAAACTGGTGGTACACAACAAAACTTCCAGCACTTAACGAACGACAGCATATAATCCAGTTTGAAAGCGCCATAACAAAAGGTAATAAGAATTTTGTTCATCACATCGAAGTGTTCCTCTGTGAGTGGAAACGTGGAATGAAAATACCCTATTTTAACGGTCCAAGTATGGTAGGCGAGAAACCTAAGGAACTAGTGTTTGTCGACATGTCATAGGAGCATGGGCTATGGGTGCGTCTGCCTTACATTTACCAGCAGANGCAGGCTGTGCAATTGGAGGCAACATCTCACGTGATATATTACTGGAAGTTCATTATAATAATCCTGAAATGAAACACGGAGTTAANGACACTTCAGGAATAAGTNCTATGTTACNCCTACACGAAGGAAGTACGATTCTGGAGTCATGGAAATTGGNTTGG*** | GCATGTCCTGGGGTGGTCTCATCACAATGGCCAGTTACAATAAATTCCACAACAACGTTTACAGAGATGCTATGTTAATCCCAGTTATTGGTTGTACAACTAGTGTTTATGCCGGATTAGTTATTTTCTCTGTTCTTGGATTCATGTCTTATGAAACTGGTATCCCAATTGAAGATGTTGTCACCCAAGGTCCTGGTCTCACATTTGTTGCTTACCCAGAAGCTGTAACTAAGTTACCAGTGTCCCCTTTGTGGGCAATTTTGTTTTTTTAATGTTATTTACCGTTGGTTTGGACAGCCAGTTTGGTATGTTCGAAACGATGACCAGTGCCTTTATTGATGAATTCCCAGAATTATTGGCCAAAAGAAAGGTTCTCTTTACTGCTGCCATTTGCTTCGTGGAGTTTCTACTTGGCATTCCCTTTGTTACACAAGGTGGAATTTACCTGCTCCAAGTTGTTGATTGGTACTGCTCCACATTTTCCCTAATGATTCTTTCCTTGACTGAATGTGTGGTCATTTCTTGGGTTTATGGTGCTGACCGCTTCTACAATGACATCACTTTGATGATTGGTTATAAACCCTGTGTGGTGGAAAATCTCATGGCGTTTCATTACTCCTACTATTACTCTGTTAGTGTGGCTGTTCAGTATTAGCAAACTGAGCCCAGTAACATATGGAGATTATACATACCCAACATGGTCAATTGTCTTTGGTTGGATTCTCGGAATGGTATCACTGGTTCCTCTACCTGCTGTGATGATTTCCCAAATATTCAAGACGAAGGGAACCTTTATTGAGCGCGT | |
| TGGCTGCGT CTGTGAGAA | CATGTGCTC TGTCTCGTT | CCGGTCTTT GGAGAATCG | AGCTGCTCC CAAGGTCAC | GCATGTCCT GGGGTGGT | |
| ACACTTGGG GCACTGTCT | TGCCTCCAC TGGGAGAGT | TTGCGTCGG TGATAGTCCT | ACGCAAGTT CCGTGCACT | ACGCGCTC AATAAAGGTT | |

**Tables S2a and S2b.** Cell Counts for each size classification and location in scattered neuron slices. Cell sizes: S- "Small" (under 12 µM soma diameter); M- "Medium" (12-15 µM soma diameter); L- "Large" (15-20 µM soma diameter), XL- "Extra large" (greater than 20 µM soma diameter). Regions: AC- Aboral Contralateral to sucker; MC- Medial Contralateral to sucker; OC- Oral Contralateral to sucker; AI- Aboral Ipsilateral to sucker; MI- Medial Ipsilateral to sucker; OI- Oral Ipsilateral to sucker. Table S2a is raw counts in triplicate, and Table S2b is mean values.

#### Table S2a

| | | TpH | | | | | | TbH | | | | | | TyH | | | | | | Bradykinin-like | | | | | | FLRIamide | | | | | |
|---|---|---|---|---|---|---|---|---|---|---|---|---|---|---|---|---|---|---|---|---|---|---|---|---|---|---|---|---|---|---|---|
| Region | Cell Size | Base 3 | Base 6 | Base 9 | Tip 1 | Tip 4 | Tip 7 | Base 3 | Base 6 | Base 9 | Tip 1 | Tip 4 | Tip 7 | Base 3 | Base 6 | Base 9 | Tip 1 | Tip 4 | Tip 7 | Base 3 | Base 6 | Base 9 | Tip 1 | Tip 4 | Tip 7 | Base 3 | Base 6 | Base 9 | Tip 1 | Tip 4 | Tip 7 |
| AC | S | 2 | 0 | 0 | 1 | 0 | 0 | 12 | 3 | 4 | 3 | 3 | 10 | 0 | 0 | 0 | 2 | 1 | 3 | 2 | 3 | 0 | 2 | 3 | 2 | 3 | 0 | 2 | 9 | 9 | 11 |
| AC | M | 1 | 1 | 0 | 0 | 0 | 1 | 1 | 2 | 0 | 0 | 2 | 0 | 0 | 0 | 0 | 1 | 1 | 0 | 0 | 2 | 0 | 0 | 2 | 1 | 3 | 1 | 0 | 3 | 5 | 3 |
| AC | L | 0 | 1 | 1 | 0 | 0 | 0 | 0 | 1 | 0 | 1 | 1 | 2 | 0 | 0 | 0 | 0 | 4 | 0 | 1 | 2 | 1 | 1 | 1 | 2 | 1 | 3 | 3 | 0 | 0 | 2 |
| AC | XL | 0 | 0 | 0 | 0 | 0 | 0 | 1 | 0 | 0 | 0 | 0 | 1 | 0 | 0 | 0 | 0 | 0 | 0 | 0 | 0 | 0 | 0 | 0 | 0 | 0 | 2 | 0 | 1 | 0 | 0 |
| MC | S | 4 | 1 | 0 | 0 | 0 | 0 | 1 | 2 | 0 | 0 | 1 | 4 | 2 | 7 | 8 | 0 | 1 | 7 | 2 | 2 | 3 | 1 | 7 | 8 | 23 | 23 | 18 | 8 | 19 | 25 |
| MC | M | 1 | 2 | 0 | 0 | 0 | 0 | 0 | 0 | 2 | 0 | 1 | 1 | 3 | 0 | 2 | 2 | 2 | 1 | 1 | 2 | 0 | 1 | 1 | 1 | 2 | 5 | 2 | 1 | 4 | 0 |
| MC | L | 0 | 0 | 0 | 0 | 0 | 1 | 0 | 0 | 0 | 0 | 1 | 0 | 0 | 0 | 0 | 0 | 0 | 0 | 0 | 0 | 0 | 0 | 1 | 1 | 1 | 2 | 1 | 0 | 1 | 0 |
| MC | XL | 0 | 0 | 0 | 0 | 0 | 0 | 0 | 0 | 0 | 0 | 0 | 0 | 0 | 0 | 0 | 0 | 0 | 0 | 0 | 0 | 1 | 0 | 0 | 0 | 0 | 0 | 0 | 0 | 0 | 0 |
| OC | S | 0 | 0 | 0 | 0 | 0 | 0 | 0 | 0 | 1 | 0 | 0 | 0 | 4 | 2 | 7 | 0 | 0 | 0 | 2 | 11 | 8 | 8 | 2 | 1 | 12 | 13 | 22 | 8 | 3 | 7 |
| OC | M | 1 | 0 | 1 | 0 | 0 | 1 | 0 | 0 | 1 | 0 | 1 | 0 | 1 | 2 | 1 | 0 | 0 | 0 | 1 | 3 | 3 | 0 | 0 | 2 | 3 | 6 | 7 | 3 | 5 | 3 |
| OC | L | 0 | 1 | 0 | 0 | 0 | 0 | 0 | 0 | 0 | 0 | 0 | 0 | 0 | 1 | 1 | 0 | 0 | 0 | 1 | 0 | 3 | 0 | 2 | 0 | 3 | 0 | 2 | 0 | 1 | 1 |
| OC | XL | 0 | 0 | 0 | 0 | 0 | 0 | 0 | 0 | 0 | 0 | 0 | 0 | 0 | 0 | 0 | 0 | 0 | 0 | 1 | 0 | 1 | 0 | 0 | 0 | 1 | 1 | 1 | 0 | 0 | 0 |
| AI | S | 0 | 0 | 0 | 1 | 0 | 0 | 11 | 6 | 13 | 2 | 2 | 1 | 0 | 0 | 2 | 0 | 0 | 0 | 4 | 1 | 0 | 1 | 0 | 3 | 6 | 1 | 3 | 6 | 1 | 3 |
| AI | M | 0 | 0 | 2 | 0 | 0 | 0 | 0 | 0 | 1 | 1 | 3 | 0 | 0 | 0 | 0 | 2 | 0 | 0 | 2 | 1 | 0 | 2 | 1 | 1 | 5 | 0 | 1 | 2 | 0 | 1 |
| AI | L | 0 | 3 | 2 | 0 | 0 | 0 | 1 | 0 | 0 | 0 | 2 | 0 | 1 | 1 | 0 | 0 | 0 | 0 | 0 | 1 | 0 | 0 | 2 | 0 | 0 | 1 | 2 | 1 | 2 | 1 |
| AI | XL | 1 | 1 | 0 | 0 | 0 | 0 | 0 | 0 | 0 | 0 | 0 | 0 | 0 | 0 | 0 | 0 | 0 | 0 | 0 | 0 | 0 | 0 | 0 | 0 | 1 | 1 | 0 | 0 | 0 | 0 |
| MI | S | 1 | 2 | 1 | 0 | 0 | 0 | 7 | 1 | 6 | 1 | 0 | 7 | 2 | 3 | 9 | 1 | 0 | 6 | 3 | 3 | 8 | 5 | 2 | 3 | 17 | 16 | 22 | 12 | 12 | 22 |
| MI | M | 1 | 1 | 2 | 0 | 0 | 0 | 2 | 0 | 2 | 0 | 0 | 4 | 3 | 1 | 0 | 2 | 1 | 2 | 0 | 2 | 1 | 0 | 1 | 1 | 3 | 3 | 7 | 1 | 0 | 1 |
| MI | L | 3 | 2 | 1 | 0 | 0 | 1 | 0 | 2 | 0 | 0 | 0 | 0 | 1 | 2 | 4 | 1 | 0 | 0 | 0 | 0 | 0 | 0 | 0 | 1 | 3 | 0 | 3 | 0 | 0 | 2 |
| MI | XL | 0 | 1 | 0 | 0 | 0 | 0 | 0 | 0 | 0 | 0 | 0 | 0 | 0 | 0 | 0 | 0 | 0 | 0 | 0 | 0 | 0 | 0 | 0 | 0 | 0 | 0 | 0 | 0 | 0 | 0 |
| OI | S | 0 | 0 | 0 | 0 | 2 | 0 | 4 | 0 | 1 | 0 | 1 | 0 | 4 | 2 | 6 | 0 | 1 | 0 | 6 | 6 | 6 | 3 | 9 | 1 | 17 | 18 | 20 | 2 | 8 | 8 |
| OI | M | 2 | 0 | 1 | 0 | 0 | 0 | 0 | 0 | 0 | 0 | 1 | 0 | 2 | 2 | 1 | 0 | 1 | 0 | 0 | 1 | 2 | 1 | 1 | 0 | 1 | 3 | 7 | 1 | 3 | 1 |
| OI | L | 0 | 1 | 0 | 0 | 0 | 0 | 1 | 0 | 0 | 0 | 1 | 0 | 1 | 0 | 0 | 0 | 0 | 0 | 2 | 1 | 1 | 0 | 2 | 0 | 1 | 0 | 2 | 0 | 0 | 1 |
| OI | XL | 0 | 0 | 0 | 0 | 0 | 0 | 0 | 0 | 0 | 0 | 0 | 0 | 0 | 0 | 0 | 0 | 0 | 0 | 0 | 0 | 0 | 1 | 0 | 0 | 0 | 0 | 0 | 0 | 0 | 0 |

**Table S2b.** Mean cell counts

| Location | Cell Size | Brady Tips | Brady Base | FLRI Tips | FLRI Base | TyH Tips | TyH Base | TpH Tips | TpH Base | TbH Tips | TbH Base |
|---|---|---|---|---|---|---|---|---|---|---|---|
| AC | S | 2.3 | 1.7 | 9.7 | 1.7 | 2.0 | 0.0 | 0.3 | 0.7 | 5.3 | 6.3 |
| AC | M | 1.0 | 0.7 | 3.7 | 1.3 | 0.7 | 0.0 | 0.3 | 0.7 | 0.7 | 1.0 |
| AC | L | 1.3 | 1.3 | 0.7 | 2.3 | 1.3 | 0.0 | 0.0 | 0.7 | 1.3 | 0.3 |
| AC | XL | 0.0 | 0.0 | 0.3 | 0.7 | 0.0 | 0.0 | 0.0 | 0.0 | 0.3 | 0.3 |
| MC | S | 5.3 | 2.3 | 17.3 | 21.3 | 2.7 | 5.7 | 0.0 | 1.7 | 1.7 | 1.0 |
| MC | M | 1.0 | 1.0 | 1.7 | 3.0 | 1.7 | 1.7 | 0.0 | 1.0 | 0.7 | 0.7 |
| MC | L | 0.7 | 0.0 | 0.3 | 1.3 | 0.0 | 0.0 | 0.3 | 0.0 | 0.3 | 0.0 |
| MC | XL | 0.0 | 0.3 | 0.0 | 0.0 | 0.0 | 0.0 | 0.0 | 0.0 | 0.0 | 0.0 |
| OC | S | 3.7 | 7.0 | 6.0 | 15.7 | 0.0 | 4.3 | 0.0 | 0.0 | 0.0 | 0.3 |
| OC | M | 0.7 | 2.3 | 3.7 | 5.3 | 0.0 | 1.3 | 0.3 | 0.7 | 0.3 | 0.3 |
| OC | L | 0.7 | 1.3 | 0.7 | 1.7 | 0.0 | 0.7 | 0.0 | 0.3 | 0.0 | 0.0 |
| OC | XL | 0.0 | 0.7 | 0.0 | 1.0 | 0.0 | 0.0 | 0.0 | 0.0 | 0.0 | 0.0 |
| AI | S | 1.3 | 1.7 | 3.3 | 3.3 | 0.0 | 0.7 | 0.3 | 0.0 | 1.7 | 10.0 |
| AI | M | 1.3 | 1.0 | 1.0 | 2.0 | 0.7 | 0.0 | 0.0 | 0.7 | 1.3 | 0.3 |
| AI | L | 0.7 | 0.3 | 1.3 | 1.0 | 0.0 | 0.7 | 0.0 | 1.7 | 0.7 | 0.3 |
| AI | XL | 0.0 | 0.0 | 0.0 | 0.7 | 0.0 | 0.0 | 0.0 | 0.7 | 0.0 | 0.0 |
| MI | S | 3.3 | 4.7 | 15.3 | 18.3 | 2.3 | 4.7 | 0.0 | 1.3 | 2.7 | 4.7 |
| MI | M | 0.7 | 1.0 | 0.7 | 4.3 | 1.7 | 1.3 | 0.0 | 1.3 | 1.3 | 1.3 |
| MI | L | 0.3 | 0.0 | 0.7 | 2.0 | 0.3 | 2.3 | 0.3 | 2.0 | 0.0 | 0.7 |
| MI | XL | 0.0 | 0.0 | 0.0 | 0.0 | 0.0 | 0.0 | 0.0 | 0.3 | 0.0 | 0.0 |
| OI | S | 4.3 | 6.0 | 6.0 | 18.3 | 0.3 | 4.0 | 0.7 | 0.0 | 0.3 | 1.7 |
| OI | M | 0.7 | 1.0 | 1.7 | 3.7 | 0.3 | 1.7 | 0.0 | 1.0 | 0.3 | 0.0 |
| OI | L | 0.7 | 1.3 | 0.3 | 1.0 | 0.0 | 0.3 | 0.0 | 0.3 | 0.3 | 0.3 |
| OI | XL | 0.3 | 0.0 | 0.0 | 0.0 | 0.0 | 0.0 | 0.0 | 0.0 | 0.0 | 0.0 |

Table S3. Generalized linear mixed models were fitted to the cell count data to estimate the fixed effects of cell location. Here we compared the number of cells in arm tip vs base for each SCN transcript. Significant adjusted p-values are in **bold**.

| **Transcript** | **estimate: log coefficient comparing base to tip** | **p-value** |
|---|---|---|
| **Tyrosine Hydroxylase** | -0.7365316 | **0.00161168** |
| **Tryptophan Hydroxylase** | -1.7272209 | **6.7463E-06** |
| **Tyramine beta Hydroxylase** | -0.457131 | 0.19596546 |
| **FLRIamide** | -0.3939288 | **0.00070369** |
| **Bradykinin-like neuropeptide** | -0.1619693 | 0.25602834 |

**Table S4.** Generalized linear mixed models (GLMMs) were fitted to the cell count data to estimate the fixed effects of cell location and size. Here we compared the number of cells of each size (S, M, L, XL) in tip vs base for each SCN transcript. Post-hoc correction was applied to account for multiple comparisons using hard Bonferroni adjustments. Specifically, adjusted p-values were calculated by multiplying original p-values by the 4. Significant adjusted p-values are in **bold**.

| Transcript | Size | estimate: log coefficient comparing base to tip (positive value indicates more cells in tip, negative value indicates more cells in base) | adjusted p-value |
|---|---|---|---|
| Tyrosine Hydroxylase | Small | -0.9662623 | **0.00306375** |
| | Medium | -0.1791971 | >0.99 |
| | Large | -0.8723173 | 0.44956048 |
| | X-Large | 0.06325243 | >0.99 |
| Tryptophan Hydroxylase | Small | -1.0116009 | 0.33269101 |
| | Medium | -2.0794416 | **0.02224494** |
| | Large | -2.0149032 | **0.0297456** |
| | X-Large | -20.165578 | >0.99 |
| Tyramine beta Hydroxylase | Small | -0.7503824 | 0.17594755 |
| | Medium | 0.21233604 | >0.99 |
| | Large | 0.44112632 | >0.99 |
| | X-Large | -0.0284681 | >0.99 |
| FLRIamide | Small | -0.3125544 | 0.054118 |
| | Medium | -0.4686267 | 0.14396791 |
| | Large | -0.8494148 | 0.06503124 |
| | X-Large | -1.9479534 | 0.27605601 |
| Bradykinin-like neuropeptide | Small | -0.1376214 | >0.99 |
| | Medium | -0.2719337 | >0.99 |
| | Large | -3.616E-10 | >0.99 |
| | X-Large | -1.0986123 | >0.99 |

Table S5.
Generalized linear mixed models (GLMMs) were fitted to the cell count data to estimate the fixed effects of cell location. Here we compared the number of cells in each vertical region (oral, medial, and aboral) in arm tip and in the arm base for each SCN transcript. Post-hoc correction was applied to account for multiple comparisons using hard Bonferroni adjustments. Specifically, adjusted p-values were calculated by multiplying original p-values by the 3 comparisons conducted within each data set. Significant adjusted p-values are in **bold**.

| Transcript | Location | Comparison (variable 1 vs variable 2) | estimate: log coefficient (positive value indicates more cells in variable 1, negative value indicates more cells in variable 2) | adjusted p-value |
|---|---|---|---|---|
| **Tyrosine Hydroxylase** | **Tip** | Medial vs Dorsal | 0.61905218 | 0.185059578 |
| | | Ventral vs Dorsal | -1.9459101 | **0.030141893** |
| | | Ventral vs Medial | -2.5649711 | **0.001274823** |
| | **Base** | Medial vs Dorsal | 2.463882456 | **6.24E-06** |
| | | Ventral vs Dorsal | 2.224623552 | **7.12E-05** |
| | | Ventral vs Medial | -0.239219992 | 0.825716057 |
| **Tryptophan Hydroxylase** | **Tip** | Medial vs Dorsal | -0.4054651 | >0.99 |
| | | Ventral vs Dorsal | 4.86567E-18 | >0.99 |
| | | Ventral vs Medial | 0.405465108 | >0.99 |
| | **Base** | Medial vs Dorsal | 0.427444015 | 0.593296343 |
| | | Ventral vs Dorsal | -0.762140052 | 0.287729405 |
| | | Ventral vs Medial | -1.189584067 | **0.017564719** |
| **Tyramine beta Hydroxylase** | **Tip** | Medial vs Dorsal | -0.530629436 | 0.172768993 |
| | | Ventral vs Dorsal | -2.140060508 | **0.000150072** |
| | | Ventral vs Medial | -1.609433952 | **0.008134595** |
| | **Base** | Medial vs Dorsal | -0.806479743 | **0.002313111** |
| | | Ventral vs Dorsal | -1.945910012 | **6.86E-07** |
| | | Ventral vs Medial | -1.139434162 | **0.014493499** |
| **FLRIamide** | **Tip** | Medial vs Dorsal | 0.587789381 | **0.000770237** |
| | | Ventral vs Dorsal | -0.087011377 | >0.99 |
| | | Ventral vs Medial | -0.674790144 | **0.000132801** |
| | **Base** | Medial vs Dorsal | 1.353718191 | **1.45E-13** |
| | | Ventral vs Dorsal | 1.278081682 | **4.95E-12** |
| | | Ventral vs Medial | -0.075634911 | >0.99 |
| **Bradykinin-like neuropeptide** | **Tip** | Medial vs Dorsal | 0.348304327 | 0.57401575 |
| | | Ventral vs Dorsal | 0.318459246 | 0.704770688 |
| | | Ventral vs Medial | -0.029852963 | >0.99 |
| | **Base** | Medial vs Dorsal | 0.336472237 | 0.751332181 |
| | | Ventral vs Dorsal | 1.08180517 | **8.71E-05** |
| | | Ventral vs Medial | 0.745340968 | **0.003402951** |

Table S6. Generalized linear mixed models (GLMMs) were fitted to the cell count data to estimate the fixed effects of cell location. Here we compared the number of cells in the ipsilateral vs contralateral (to the sucker) sides for SCN transcript in the base and in the tip. Significant adjusted p-values are in **bold**.

| Transcript | Location | estimate: log coefficient comparing ipsilateral cell number to contralateral cell number | p-value |
|---|---|---|---|
| **Tyrosine Hydroxylase** | **Tip** | -0.3856625 | 0.21989413 |
| | **Base** | 0.13657625 | 0.52167904 |
| **Tryptophan Hydroxylase** | **Tip** | 1.11E-16 | >0.99 |
| | **Base** | 0.13657625 | 0.52167904 |
| **Tyramine beta Hydroxylase** | **Tip** | -0.2076437 | 0.4279367 |
| | **Base** | 0.6264583 | **0.00468615** |
| **FLRIamide** | **Tip** | -0.3719462 | **0.00627023** |
| | **Base** | -0.0121168 | 0.91236503 |
| **Bradykinin-like neuropeptide** | **Tip** | -0.1984509 | 0.34623874 |
| | **Base** | -0.0935261 | 0.62895933 |

Supplementary Chromogenic Images
(Slices are 20 μM thick)

**Tryptophan hydroxylase (Probe 51):** Cross sections
20x. Dorsal ANC (axon tracts) oriented toward the top of the image from viewer's perspective

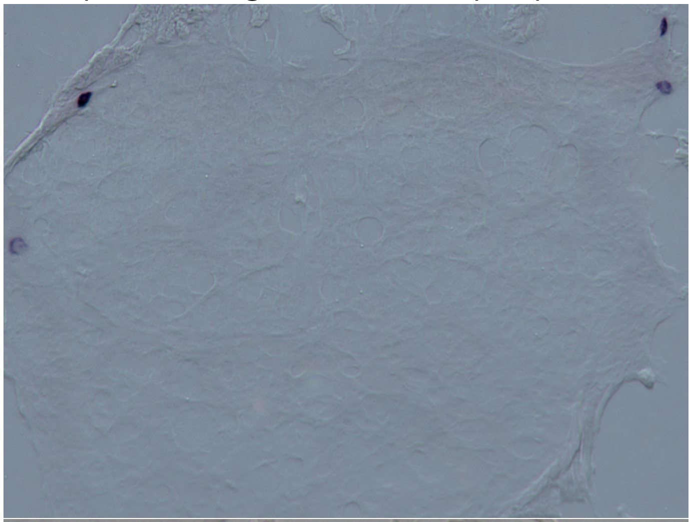

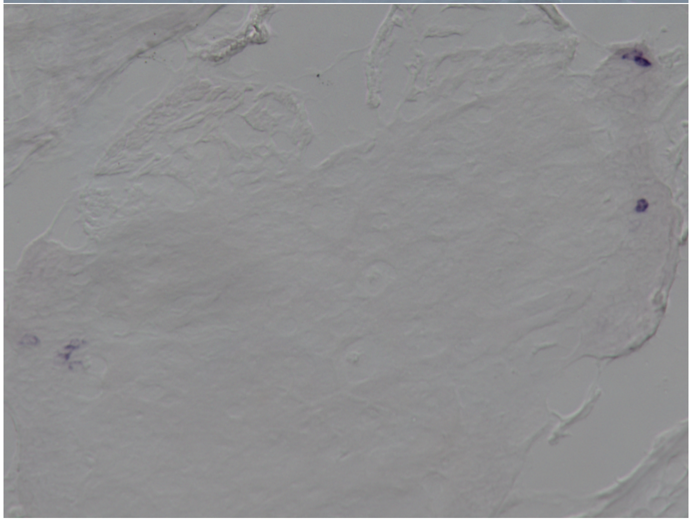

20x. Dorsal ANC (tracts) oriented toward the top left of the image from viewer's perspective

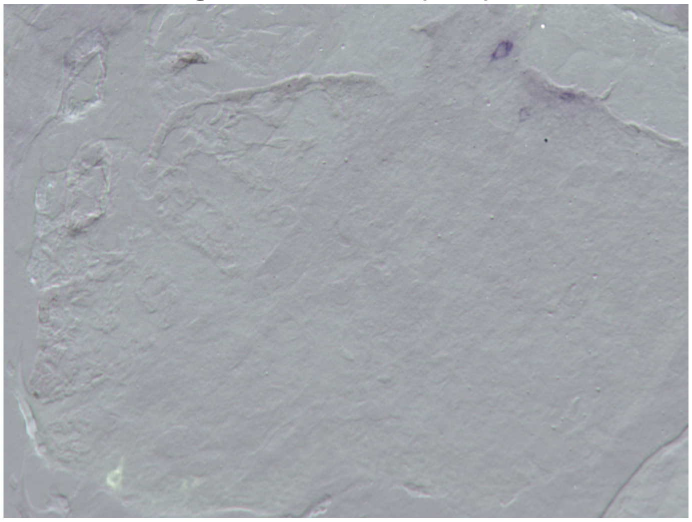

**Tyrosine Hydroxylase** (Probe 42)

Longitudinal slices (10x and 20x). Ventral ANC is toward the left side of image from viewer's perspective

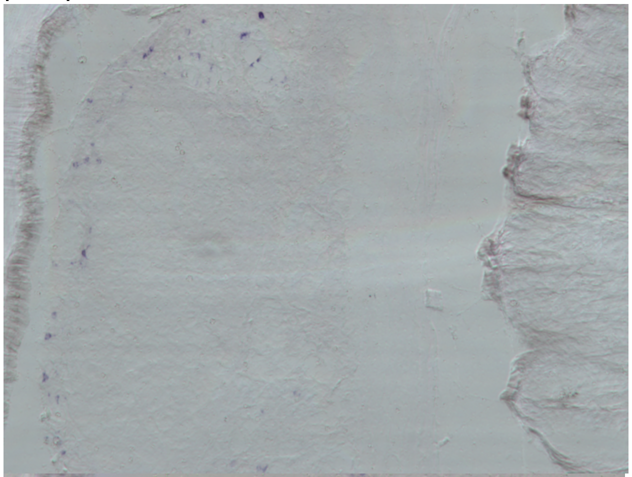

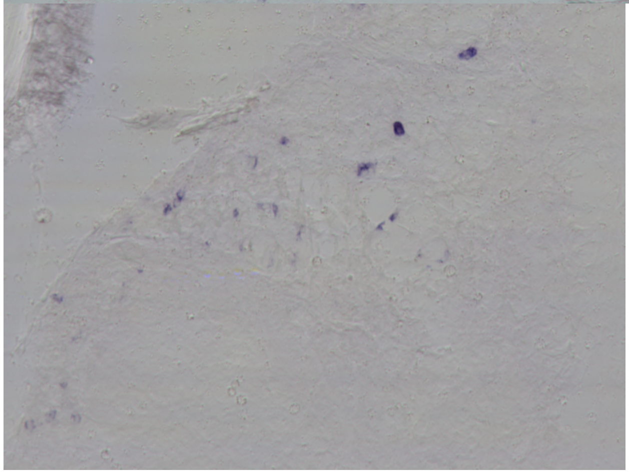

Cross section (20x): Ventral ANC oriented toward the top left of the image

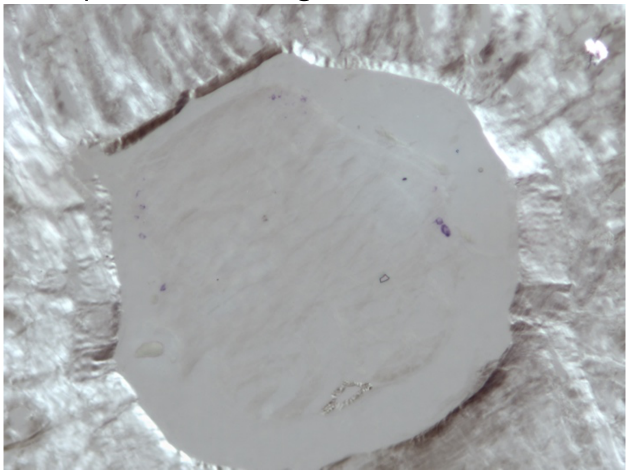

Cross section (40x): Ventral ANC oriented toward the top left of the image

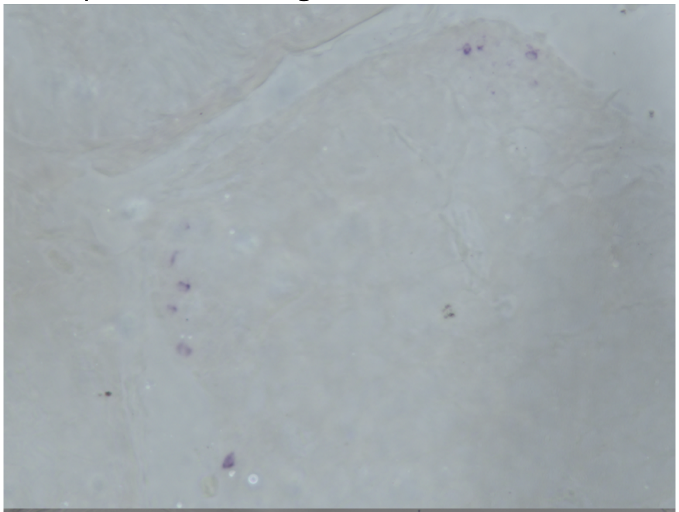

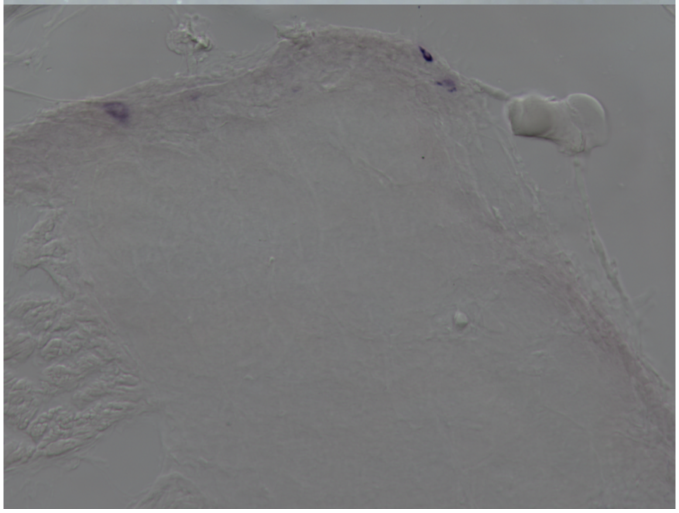

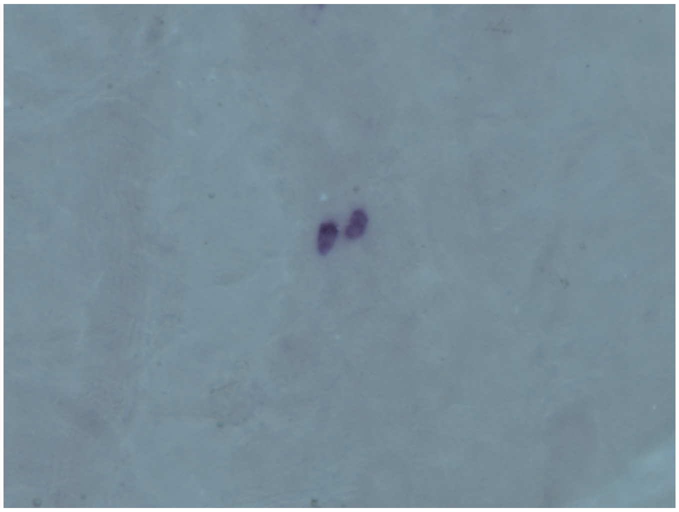

**Tyramine β Hydroxylase** (Probe 25) 10x and 40x. Cross sections: Ventral ANC is oriented toward the bottom left of the image from the viewer's perspective. Axon tracts are toward the top right.

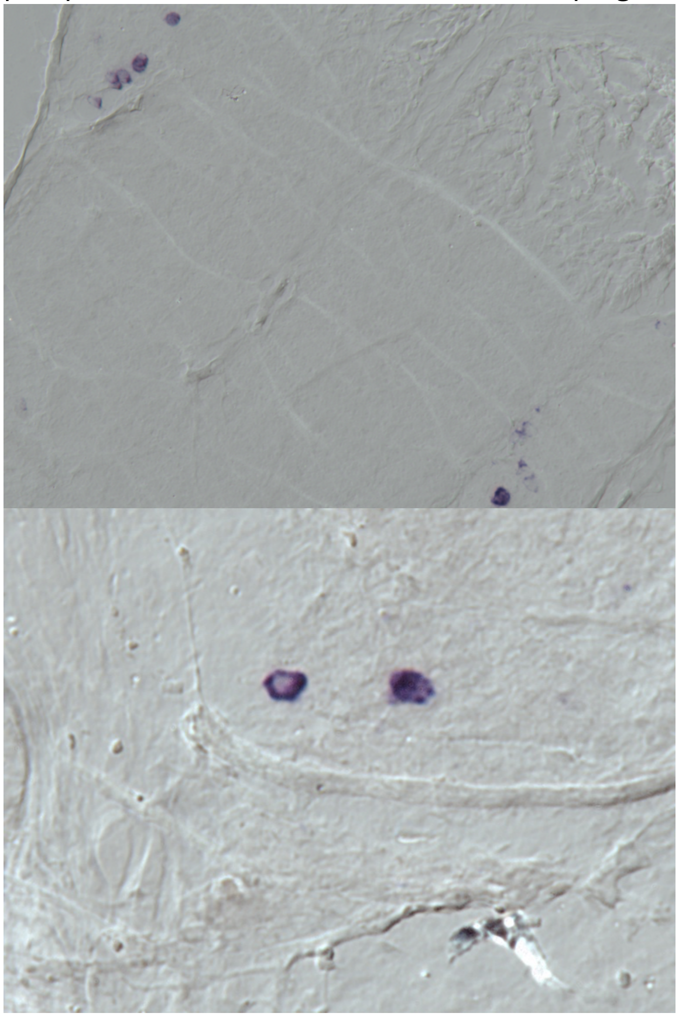

**Choline Acetyltransferase** (Probe 32)
Longitudinal slices 10x, 20x, and 40x. Ventral is left side of image from viewer's perspective

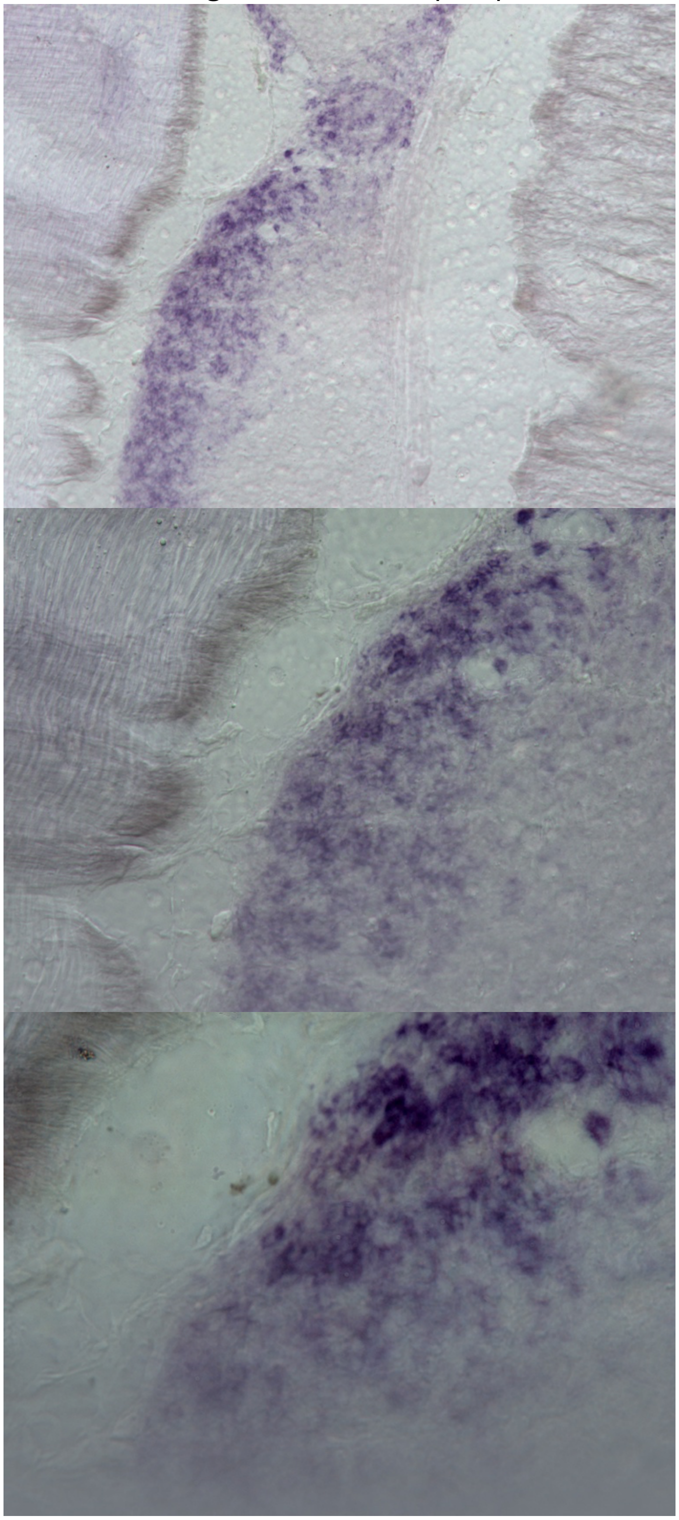

Cross sections at 10x, 20x, and 40x. Ventral ANC is oriented toward the top left of the image from the viewer's perspective

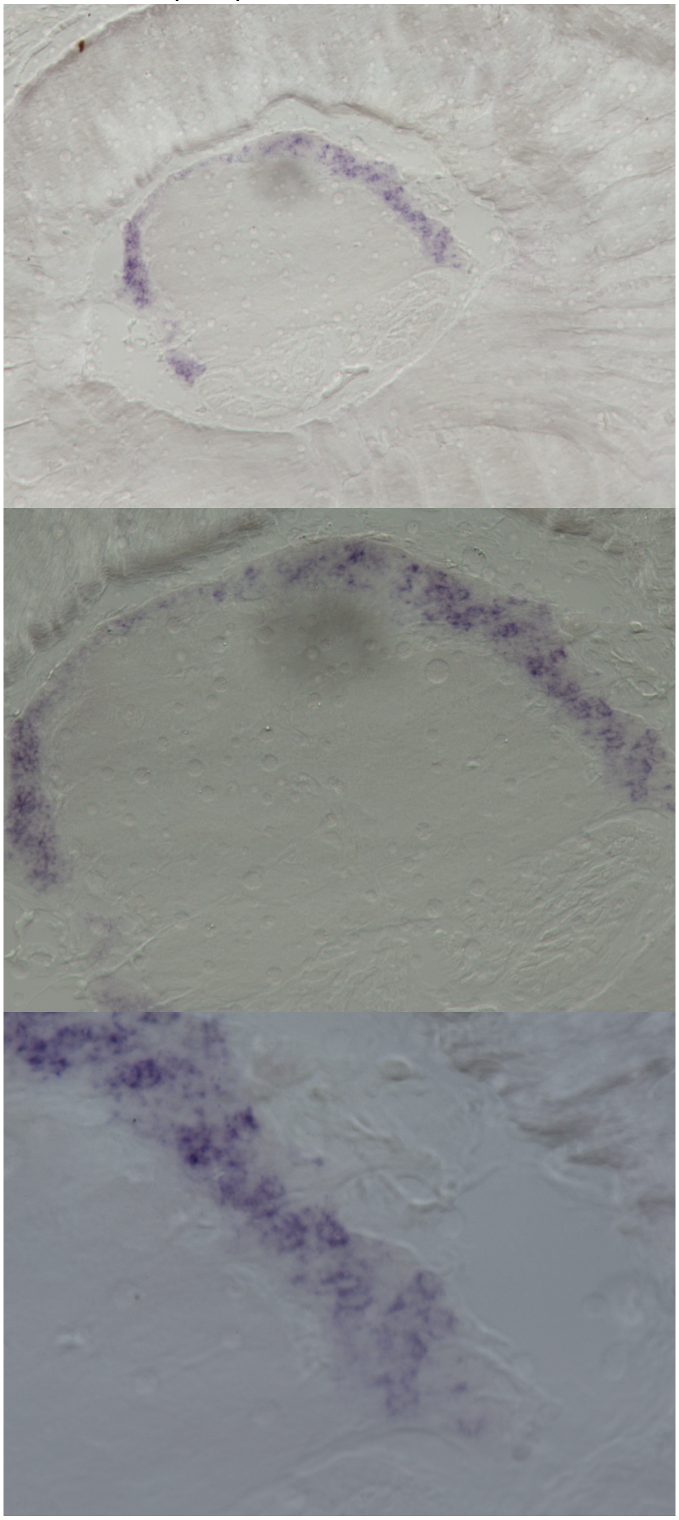

**Vesicular Glutamate Transporter** (Probe 9)
20x Cross section. Ventral ANC is oriented toward the bottom of the image.

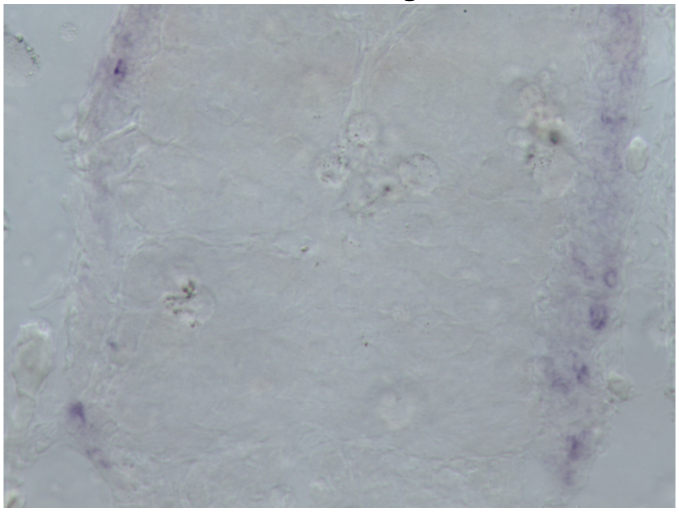

**Excitatory amino acid transporter1** (Probe 39)
20x Longitudinal section showing neuropil

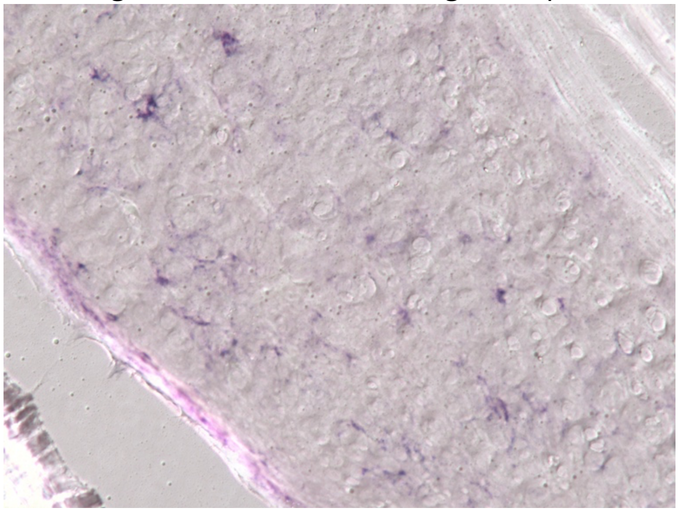

**Vesicular GABA Transport**er (Probe 6)

10x Cross section. Ventral ANC is oriented toward the bottom of the image. (Purple "smudge" toward ventral region is background due to tissue abnormalities from human error)

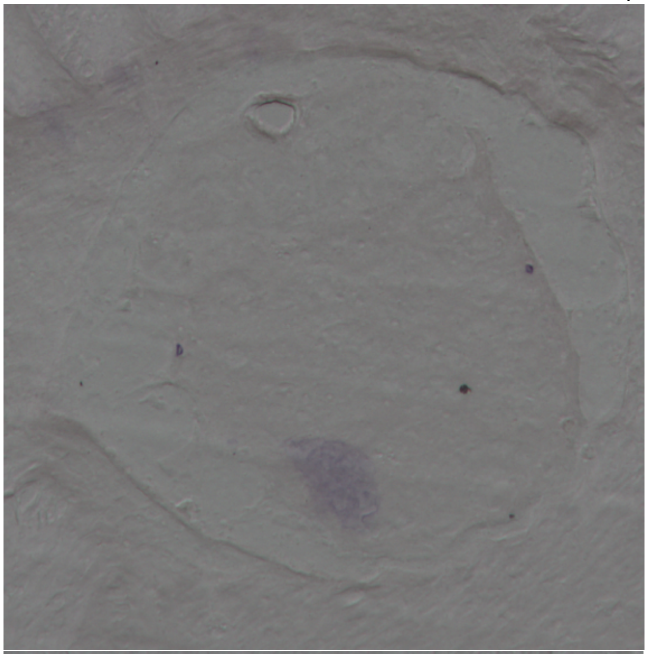

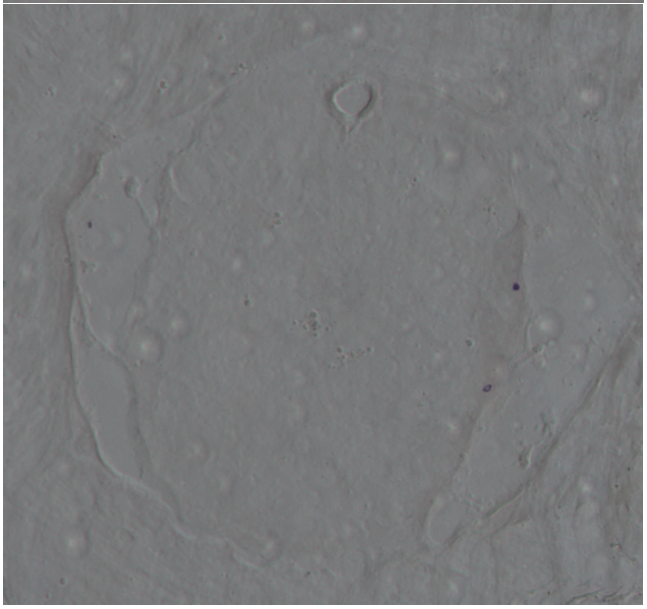

20x Cross section. Ventral ANC is oriented toward the top of the image.

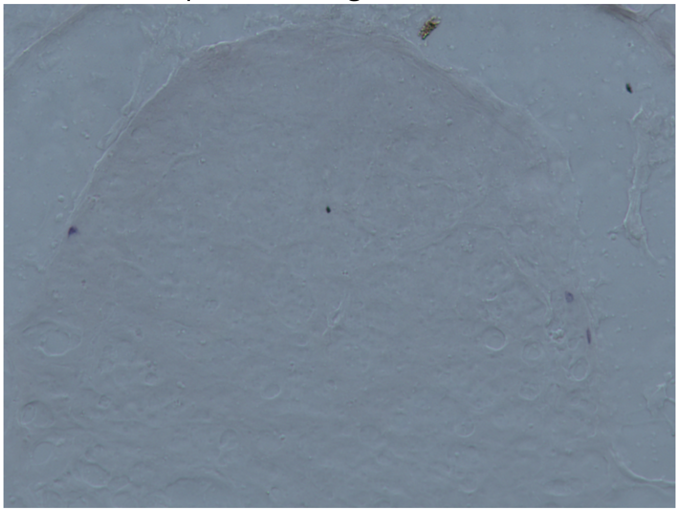

20x Cross section. Ventral ANC is oriented toward the right side of the image from the viewer's perspective.

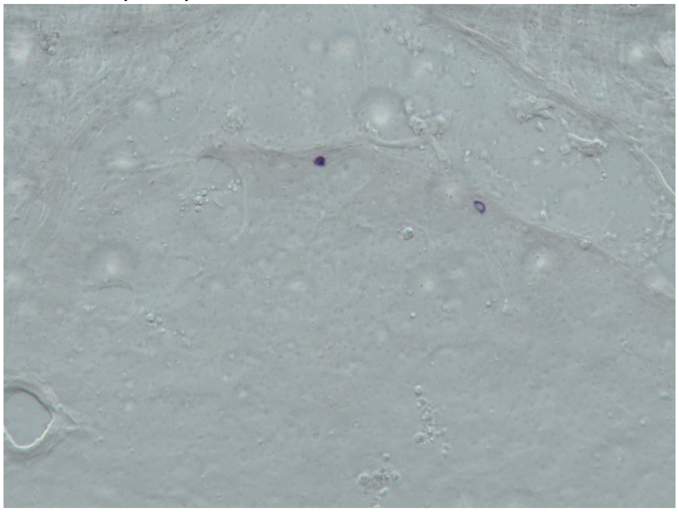

FLRIamide (Probe 16)

Ventral ANC oriented toward the top of the image
(10x and 20x cross sections)

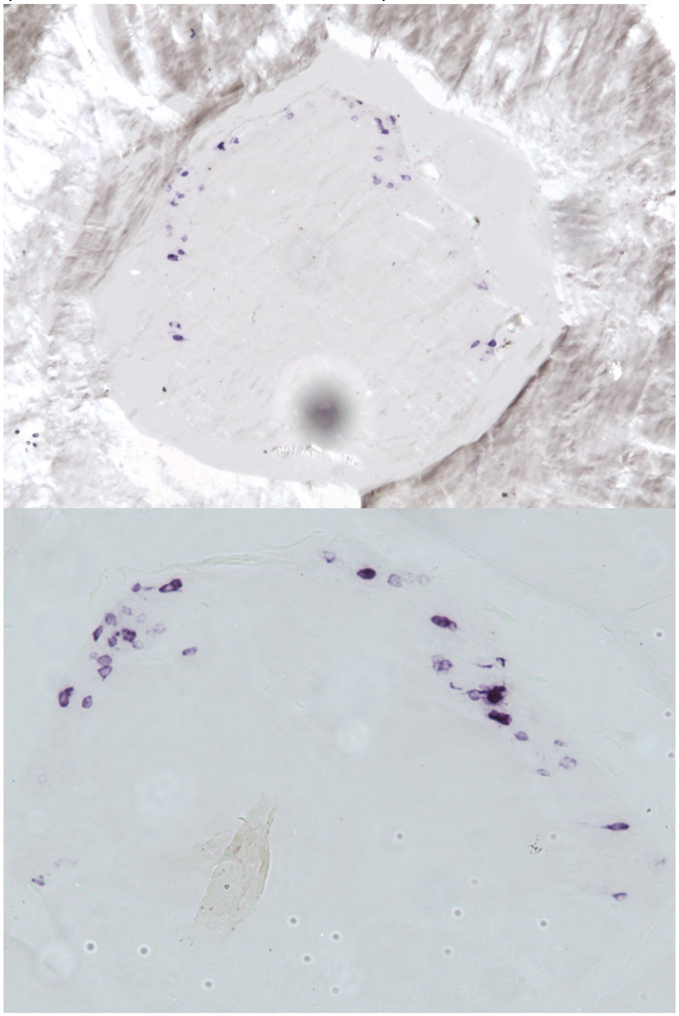

Lateral regions of ANC cortical layer at 40x
(40x cross sections)

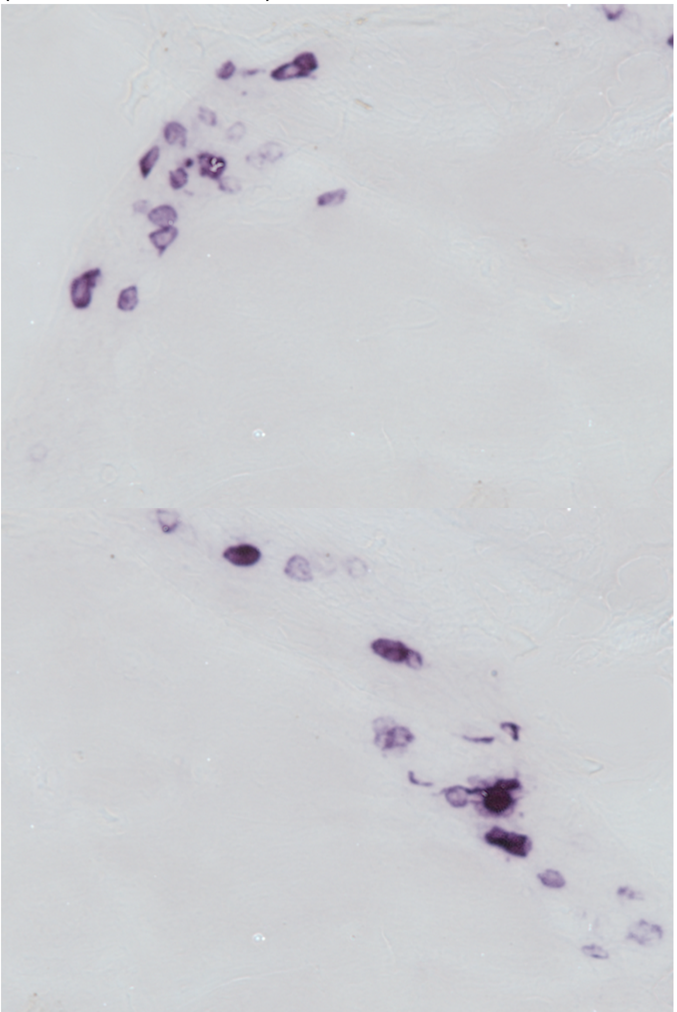

**Bradykinin-like Neuropeptide (Probe 2):** Cross sections 10x, 20x and 40x
Dorsal ANC (axon tracts) oriented toward the right of the image from viewer's perspective

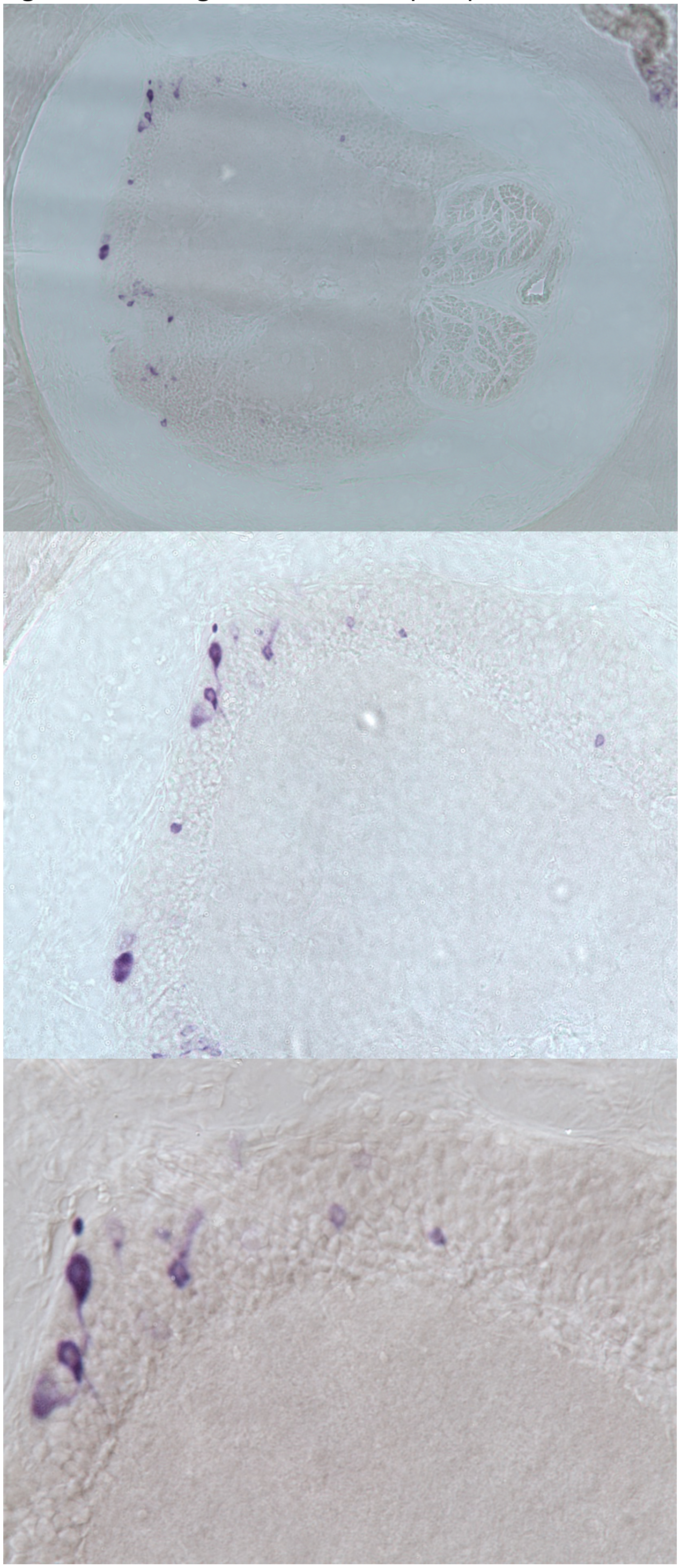

**Glycine Transporter (Probe 45):**
Cross section20x.
Dorsal ANC (axon tracts) oriented toward the top/right of the image from viewer's perspective

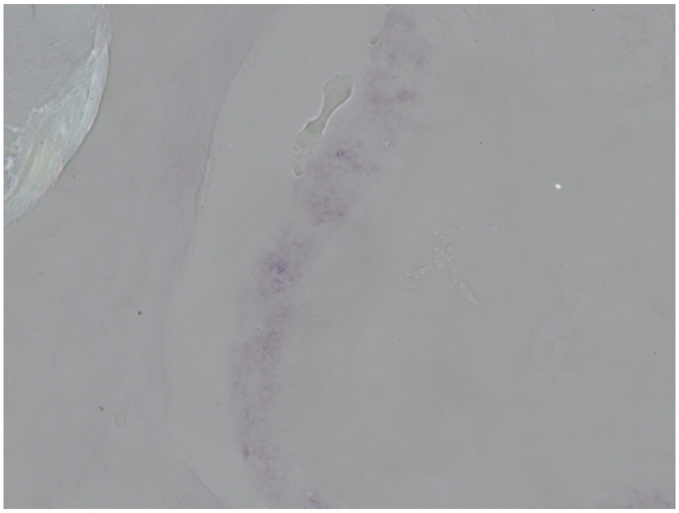

### Supplemental gene trees and aligned fasta files

**Alignment files**

Bradykinin-like neuropeptide
>Octopus bocki
-----------------------------MGINIIHVLCLVA---------------------------------------------------------VFASSVYSLPKR------------NDHVITLR-----------------------------
---------SLRQTGLSDSDSRA--------LLQAYILGKL------------SNVDIAGNKE------------------------A---DNSDYTTI-----KRKAFWRPMGYLPFDNHGGSGASSSNDNSPSGGTGSAVFRYG-
>Octopus bimaculoides- gi|918340888|gb|KOG01073.1| hypothetical protein OCBIM_22003823mg
-----------------------------MGINIIHVLCLVA---------------------------------------------------------VFASSVHSLPKR------------NDHVITLR-----------------------------
---------SLRQTGLSDSDSRA--------LLQAYILGKL------------SNVDIAGNKE------------------------A---DNSDYTTI-----KRKAFWRPMGYLPFDNHGGSGASSSNDNSPSGGTGSAVFRYG-
>Doryteuthis pealeii
-----------------------------MTVNAVHVLCIFA---------------------------------------------------------LLFACVHSLPKR------------TDHASTLR-----------------------------
---------FLQQSGLSDSDSRA--------LLQAYILGKL------------SVGDGSIGKE------------------------L---ETSEYPTI-----KRKAFWRPMGYLPFENHAGSGASSSNDNAAGGGSASAVFRYG-
>Octopus vulgaris- CAI9718386.1 Hypothetical predicted protein
-----------------------------MGINIIHVLCLVA---------------------------------------------------------VFASSVHSLPKR------------NDHVITLR-----------------------------
---------SLRQTGLSDSDSRA--------LLQAYILGKL------------SNVDIAGNKE------------------------A---DNSDYTTI-----KRKAFWRPMGYLPFDNHGGSGASSSNDNSPSGGTGSAVFRYG-
>Mytilus galloprovincialis- VDI05882.1 Hypothetical predicted protein
-----------------------------MSTSVWYICGVLC---------------------------------------------------------ALVISLTLSFEI------------DDTVLSDDEYYPQ------------------------
---------YKAVDTVSDDSRKE--------NVLAKILKYL----------DTKNAQRSALQD-------------SYVQKIFENDMD---SGARLPII----SKRKVFWQPLGYIPASVRISGNSGKSQSDDNTGGQL---FRYG-
>Pinctada margaritifera- gi|751372408|gb|AJF48838.1| hypothetical protein c45042
---------------------------MKTTILLSFSLCVTM---------------------------------------------------------TFAKSLSESFRG---------EWKQDRP---------------------------------
---------SRILSMISKSQQKE--------LELIQTLMRMTADRNKSLFFDEEDYDPSMESE------------------EDKMKNI---LSEDVKSV----SKRKVFWQPLGYVPASLRM-NGGSREQSGESSSRAGGNILRYG-
>Bulinus truncatus- KAH9498949.1 Bradykinin-like neuropeptide
MSTTSHYIAPRVTTLHHESLHCTTSHYIAPRVTTLHHESLHCTTSHHIAPRVTTLHHESPHCTTSHHIAPRVTTLHHESPHCTTSHYTAPRVTTLHHESLHCTTSHYIAPRVTTLHHESLHCTTSHYIALRVTTLHHESLLCTTTTMTSSSSVYYYVIFFT
VCLSCFVATFELEEDLNDYPDQEKRTYTASDASLEAILNIL------------KSHAQSLRQL--------ESTIYEQKRSGFRTRLG---DELNFGGV-----KRRMVWQPLGYLPASARVQHG-SQGAPRQEIQDSGSSVFRYG-
>Aplysia californica- gi|325296891|ref|NP_001191477.1| bradykininlike neuropeptide precursor
---------------------------MTSSIYGFITLSVVA---------------------------------------------------------LISQTTC-----------------RSLDLLLD-----------------------------
---------GDFNNGLASFDGSSKWSRLPLEFLAAFDLDPH------------QAQGQHLAEAPEAPLMEAMKRSRGPSPRRLRSYLR---RAAGLRGM-----KRKMFWQPLGYMPASARA-HNNVPEVVNENSQDSGTNVFRYG-
>Elysia marginata- GFR72530.1 bradykinin-like neuropeptide precursor
-----------------------------MYMSVLPMLALAVL--------------------------------------------------------MFFPPTFGAPRQ------------RYSIADLR-----------------------------
-------------DAFARPSSHK--FYLPIEIAPAALLGAFKAHARNSARADGSDADIDAIPELPLDLVQQKRTTSSMEMVDPLSAAALVQESPVFSGM-----KRKMFWQPLGYMPASARAQHSKNSGGGRNSPQENGSNVFRYG-
>Mizuhopecten yessoensis- AXN93507.1 NKY
-----------------------------MTQGTSLLLVVFA---------------------------------------------------------NFFVVQCYGSYL------------PGALGLSK-----------------------------
---------SNEASNYIDSVITE-----EEKAFYAKVIQKL------------MDRAAALEEE---------VDGHNGDNFDSDGDLS---SSSDLSGLKRSMDKRKVFWQPLGYVPASMRMSPNNKHKASQKDVGRKG---FRYGK
>Candidula unifasciata- CAG5125567.1 unnamed protein product
---------------------------MAWTINTLVKLVLLA---------------------------------------------------------LIPLC-YSRPSL------------LDGE---------------------------------
-----------DDPGYIQDQESPRRFYVPVEVPLDAIFSSL------------RAHTQSLHQS--SPLLDKRAASGSEGIGSILPKMD---EDADLRGM-----KRKMFWQPLGYLPASVRA--HNSPTGSSASENQASSSVFRYG-

Choline acetyltransferase
>Octopus_bocki
---MSHPHVRAYVKRVASIDQEAFPDWNLSKPLPKMPVPDISSTLQKYLSLVRPIVPSEQYEKTKAIVENF-
QKDGTGEKLQKMLLEYAESKDNWCYDWWLDEMYMKVQLPLPVNSNPGMVFPKRHFADQNAQLKYAAQLISAILDYKTIVDSRSLPIDRARSNRKGQPLCMEQYYRLFTSYRVPGLNKDTLVTNSTKLMPEPEHIIVICKNQLFVLDVMLNFSRLTDDDLY
TQLWRIAKVAEEGEKNAEPVGILTSLRRNKWASARQKLMEDSANRDSLDAIERSIFILSLDKKPPVSFNHQNSVDETREQQRDDVSMAIQMLHGMGSQVNSANRWFDKTMQFIVSEDGVCGLNYEHSPSEGIAVVQLVEHTLKYME-
ELRAMRLHRMQSICDLPYPRKLKWKLTDVLNKDIADASKAIDKLISDLDLYVLRFTNFGKEFPKSQNMSPDSFIQLALQLTYYKIHGHLVSTYESASTRRFRYGRVDNIRAASPAALEWVKSMTGE-
TNIPDEDKMRLLRAAMKTQTEIMTGTILGQGVDCHLMGLREISNEKELPIPEIFADESYKLANHFTLSTSQVPTTMDAFMCYGPVVPDGYGVCYNPHSDYIVAVVTSFRSHSETRSDY------------------------------------------
----------------------------------------------------
>XP 014768157.2 choline O-acetyltransferase partial Octopus bimaculoides
------------------------------------------------------------------------------------LISFFLQ----
CYDWWLDEMYMKVQLPLPVNSNPGMVFPKRHFADQNAQLKYAAQLISAILDYKTIVDSRSLPIDRARSNRKGQPLCMEQYYRLFTSYRVPGLNKDNLVTNSTKLMPEPEHIIVICKNQLFVLDVMLNFSRLTDDDLYTQLWRIAKVAEEGEKNAEPVGIL
TSLRRNKWASARQKLMEDSANRDSLDAIERSIFILSLDKKPPVSFNHQNSVDETREQQRDDVSMAIQMLHGMGSQVNSANRWFDKTMQFVVSEDGVCGLNYEHSPSEGIAVVQLVEHTLKYME-
ELRAMRLHRMQSICDLPYPRKLKWKLTEVLNKDIADASKAIDKLISDLDLYVLRFTNFGKEFPKSQNMSPDSFIQLALQLTYYKIHGHLVSTYESASTRRFRYGRVDNIRAASPAALEWVKSMTGE-
TNIPDEDKMRLLRTAMKTQTEIMTQTILGQGVDCHLMGLREISNEKELPIPEIFADESYKLANHFTLSTSQVPTTMDAFMCYGPVVPDGYGVCYNPHSDYIVAVVTSFRSHSETRSDYFAFTLESSFLQMYELCYKTKDMDKNTEQKIPEETHATK----
QQQEQQQQPQE---PSQKSELKKTDSQTEKTNSERSGATTEESSGLNKNDAK
>CAI9722745.1 choline O-acetyltransferase-like Octopus vulgaris
-------------------------------------------------------------------------------
MHYAPLVSLNIQNCSECYDWWLDEMYMKVQLPLPVNSNPGMVFPKRHFADQNAQLKYAAQLISAILDYKTIVDSRSLPIDRARSNRKGQPLCMEQYYRLFTSYRVPGLNKDTLVTNSTKLMPEPEHIIVICKNQLFVLDVMLNFSRLTDDDLYTQLWRIA
KVAEEGEKNAEPVGILTSLRRNKWASARQKLMEDSANRDSLDAIERSIFILSLDKKPPVSFNHQNSVDETREQQRDDVSMAIQMLHGMGTQVNSANRWFDKTMQFVVSEDGVCGLNYEHSPSEGIAVVQLVEHTLKYME-
ELRAMRLHRMQSICDLPYPRKLKWKLTEVLNKDIADASKAIDKLISDLDLYVLRFTNFGKEFPKSQNMSPDSFIQLALQLTYYKIHGHLVSTYESASTRRFRYGRVDNIRAASPAALEWVKSMTGE-
TNIPDEDKMRLLRAAMKTQTEIMTQTILGQGVDGHLMGLREIANEKEQPLPEIFADESYKLANHFTLSTSQVPTTMDAFMCYGPVVPDGYGVCYNPHSDYIVAVVTSFRSHSETRSDYFAFTLESSFLQMYELCFKTKDMDKNTEQKIPEETHAKTEQQE
QQQQQQQQPQEQQQPSQKSELKKTDSQTEKTNSEGSGAKTEEGSGLNKNDAK
>XP 034312330.1 choline O-acetyltransferase Crassostrea gigas

---
MSTRHVKEYVKRVASIDQEQYPPWDLNQPLPKLPVPDLHKTLSKYEDILQPILNKSQFKEAKEIISEFGKKGGHGEHLQELLVLDAETKENWAYNWWLNDMYMKIRLPLPINSNPGMVFPKQCFSDRREQLRYAARLISGILDYKTIIDARGLPVDRARH
NKKGQPMCMEQYYRLFTSYRVPGIKKDSLISNNSKLLPDPEHIIVISKNQYFVLDVIINFTRLSEEDIYTQLSRICKMAEENKDDVDPVGILTAAYRDKWALARGKLMEESTNRDSLDAIERSIFVLCLDTAAPVIPSHQNSIKTIS-
SVRDDVSLASQMLHGLGTHINSANRWYDKTMQFIISEDGACGLNYEHSPSEGIAVVQLIEHLLKYME-EVR-
KKLARMQSLCELPQPRRLQWKINKDTETDIKEAIENIDRLIKDLDLFILRFKSFGREFPKSQNMSPDSFIQLALQLTYYKIHGKLVSTYESASVRRFRLGRVDNIRANTPDALEWIRAMVGE-
IETTDVEKMALLRKAMQCQTDIMTQTILGHGMDCHLLGLREIARENGLPMPKIFEHESFRISNHFSLSTSQVPTTMDAFMCYGPVVSDGYGVCYNPHPSWILVCITSFKRHNDTNSDHFAYTLESSLLQMQELCLKTSEAKP---ILYGGKTHSEN----
-RNGAIVDDKEMSAGTRKSRLVR---QRTMPFPEVIGKDS------------
>XP 030835119.1 choline O-acetyltransferase Strongylocentrotus purpuratus
MNHMPNRHHSHYVAN-GEIDDDITNPIHLR-
PLPTLPVPPLGQSLERYLKSLSPLVSKEQYERTQALVQEFLKPGGEGETLQQKIVEMRETKVNWTYDYWLDDMYLKVRSALPINSNPGMVFPKQYFRDEGQRLRFAAKLISGILDYKVVIDARALPVDRARHNKQGQPLCMEQYYRLFSSYRVPGLLKDE
LVSPGSSLMPEPEHIIVVCQNQFFILDVIINFKRLSDQDLVSQLKRILDAAKENP-DSVPLGLLTTADRPTWASSRARLIEDSTNRDSLDMIERCIFVLCLDERMDYPPPEVG-----
KPIDSDDEPLGLHMLHGGGVENNTGNRWFDKTMQFIIGTDGQSGLNYEHSPSEGVAVVQLVEHLLKYIDSETHKRKLTRAQSICELPNPRKLNWKISPILLQDIEIAKAQITYAVSDADVKLLRFTQFGKNFPKSQNISPDAFVQLALQLTYYRIYGRLT
STYESGSTRRFQEGRVDNIRAASPEALEFVRSIKGE-
REASIEEKKFLMREAIKAQTETCLQTITGQGFDCHLLCLREVSKEIGMPLPAIFADEAFEISNRFCLSTSQVLTKLDTFMCYGPVVPDGYGASYNPHEDYILFAIASFKSCPSTDSHVFRAKLYQTLIDMHELCTGTKPNTNNNSNGYTQG---------
----------------------------------------------------
>NP 001124191.1 choline O-acetyltransferase Danio rerio
---------------
MPVSKREQSKDTGDPCALPKLPIPPLKQTLDMYLTCMGHLVPEDQFRKTKAVVEKFGAPGGVGETLQKKLLERSEQKANWVYDYWLEDMYLNNRLALPVNSSPVMVFHKQNFKGQSDVLRFAANLISGVLEYKALIDGRALPVEHARGQLAGTPLCMDQY
NKVFTSYRLPGTKTDTLVAQKSTVMPEPEHIIVACKNQFFVLDVMVNFRRLNEKDLYTQLERIRKMADIEEERQPPIGLLTSDGRTQWAEARNILIKDSTNRDSLDMIERCLCLVCLDEE--------------
TATELNDSNRALLMLHGGGTDKNGGNRWYDKPMQFVIGADGCCGVVCEHSPFEGIVLVQCSEYLLRYMR--
GSPSKLVRAASMSELPAPRRLRWKCSPDIQTFLSASADRLQKLVKNLDMNVHKFTGYGKEFIKRQKMSPDAYVQVALQFTFYRCHGRLVPTYESASIRRFQEGRVDNIRSSTPEALAFVKAMASG-
SKITDAEKMELLWTAIKAQTNYTILAITGMAIDNHLLGLREIAKELKLEKPELFSDTTYATSIHFTLSTSQVPTTEEMFCCYGPVVPNGYGACYNPQTDHILFCVSSFRECAETSSDLFVKTLEGCLKEMQDLCRKCN-
TEVKPADSTQRMEGNPKVMKNGSKS------------------------------------------------
>BAC33133.2 unnamed protein product partial Mus musculus
---------
MPILEKVPPKMPVQASSCEEVLDLPKLPVPPLQQTLATYLQCMQHLVPEEQFRKSQAIVKRFGAPGGLGETLQEKLLERQEKTANWVSEYWLNDMYLNNRLALPVNSSPAVIFARQHFQDTNDQLRFAASLISGVLSYKALLDSQSIPTDWAKGQLSGQP
LCMKQYYRLFSSYRLPGHTQDTLVAQKSSIMPEPEHVIVACCNQFFVLDVVINFRRLSEGDLFTQLRKIVKMASNEDERLPPIGLLTSDGRSEWAKARTVLLKDSTNRDSLDMIERCICLVCLDGP--------------
GTGDFSDTHRALQLLHGGGCSLNGANRWYDKSLQFVVGRDGTCGVVCEHSPFDGIVLVQCTEHLLKHMM--
TGNKKLVRADSVSELPAPRRLRWKCSPETQGHLASSAEKLQRIVKNLDFIVYKFDNYGKTFIKKQKCSPDGFIQVALQLAYYRLYQRLVPTYESASIRRFQEGRVDNIRSATPEALAFVQAMTDHKAAVLASEKLQLLQRAIQAQTEYTVMAITGMAIDN
HLLALRELARDLCKEPPEMFMDETYLMSNRFILSTSQVPTTMEMFCCYGPVVPNGYGACYNPQPEAITFCISSFHGCKETSSVEFAEAVGASLVDMRDLCSSRQPAEGKPP------------------------------------------------
------------------
>AAI30618.1 CHAT protein Homo sapiens
-------------------MTAKTPSSEES-
GLPKLPVPPLQQTLATYLQCMRHLVSEEQFRKSQAIVQQFGAPGGLGETLQQKLLERQEKTANWVSEYWLNDMYLNNRLALPVNSSPAVIFARQHFPGTDDQLRFAASLISGVLSYKALLDSHSIPTDCAKGQLSGQPLCMKQYYGLFSSYRLPGHTQDT
LVAQNSSIMPEPEHVIVACCNQFFVLDVVINFRRLSEGDLFTQLRKIVKMASNEDERLPPIGLLTSDGRSEWAEARTVLVKDSTNRDSLDMIERCICLVCLDAP--------------
GGVELSDTHRALQLLHGGGYSKNGANRWYDKSLQFVVGRDGTCGVVCEHSPFDGIVLVQCTEHLLKHMT--
QSSRKLIRADSVSELPAPRRLRWKCSPEIQGHLASSAEKLQRIVKNLDFIVYKFDNYGKTFIKKQKCSPDAFIQVALQLAFYRLHRRLVPTYESASIRRFQEGRVDNIRSATPEALAFVRAVTDHKAAVPASEKLLLLKDAIRAQTAYTVMAITGMAIDN
HLLALRELARAMCKELPEMFMDETYLMSNRFVLSTSQVPTTTEMFCCYGPVVPNGYGACYNPQPETILFCISSFHSCKETSSSKFAKAVEESLIDMRDLCSLLPPTESKPLATKEKATRPSQGHQP----------------------------------
------------------
>ABG81467.1 dopamine beta-hydroxylase precursor partial Bos taurus
-------------------------------PSPSVREAASMYGTA------------------------------------------------VAVFLVILVAALQG-SAPAESPFPFHIP---LD----PEGTLELSWNISYAQETIYFQLLVRE---
LKAGVLFGMSDRGELENADLVVLWTDRD-GAYFGDAWSDQKGQ-VHLDSQQDYQLLRAQRTPEGLYLLFKRPFGTCDPNDYLIEDGTVHLVYGFLEEPLRSLE-SINT-SGLHT---GLQRVQLLKPS-
IPKPALPADTRTMEIRAPDVFIPGQQTTYWCYVTELPDGFPR--HHIVMYEPIVTEGNEALVHHMEVFQC-
AAEFETIPHFSGPCDSKMKPQRLNFCRHVLAAWALGAKAFYYPEEAGLAFGGPGSSRFLRLEVHYHNPLVITGRRDSSGIRLYYTAALRRFDAGIMELGLAYTPVMAIPPQETAFVLTGYCTDKCTQLALPASGIHIFASQLHTHLTGRKVVTVLARDGR
ETEIVNRDNHYSPHFQEIRMLKKVVSVQPGDVLITSCTYNTEDRRLATVGGFGILEEMCVNYVHYYPQTQLELCKSAVDPGFLHKYFRLVNRFNSEE-VCTCPQASVPEQFASVP-WNSFNREVLKALYGFAPISMHCNRSSAVRFQG--
EWNRQPLPEIVSRLEEPTPH-CPASQAQSPAGPTVLNISGGKG-----------------------------------------------------------------------------------------------------------------
-----------------------------------------------------------------------------------------------------------------------------------------------------------------
-------------------------------------------------------------------------------------------------------------------------------------------------------
>AZK16218.1 tyramine beta-hydroxylase Lymnaea stagnalis
------------------------------------MAGHAWSISA-------------------------------------------------VVWLVVLSADHSV------QAYQYQVT---LD----PDSRYLFQWSVDYKESLINVQLTCKVT--
PESWLAFGFSDYGDVTSADLILFWTDGDGKHHFFDGHTTPDGI-FLPDRQQDYHLTSVADDRGSVVLDFYRHFNTCDPEDYALDNGTTHLVYVESAQPEGPPL-ARDV-TRLRH---GVQRLQLLKPE-
ISAPVFPEDTWSFEVRAPEVLVPAEETTYWWHTTILPDMPSP--
HHIIQYEGIVAEGSGDLVHHMEVFHCQVQKGHGVPYYNGPGIAEGKPEGLEVCRKVIGAWAMGAEAMIYPEEAGVPVGGQGFSRFALLEVHYNNPQKKSGRMDSSGIRFHVTSQLRKYDAGIMELGLEYVNKMAVPPGQRDFKLSGYCVHKCTQMSLPPA
GIHVFASQLHTHLTGRRVYTKHARDGAELPEVNRDNHYSPHFQEIRRLPQPHHVLPGDVLITTCEYDTTKRSKATVGGFSITDEMCLNYVHYYPRSDLEVCKSSVRTDSLHTFFFLLNRFENSK-V--SPEAGDRANYESIE-
WSPLNVRLLEDLYSTSPLSMQCNRSDGTRFPG--EWEHVRVPDIVRPLVVDTSEVCSGHVTAE-------------------------------------------------------------------------------------------------
-----------------------------------------------------------------------------------------------------------------------------------------------------------------
-----------------------------------------------------------------------------------------------------------------------------------------------------------------
-------------------------
>CAA94391.2 tyramine-beta-hydroxylase Drosophila melanogaster
---------------------------MLKMPLQLSSQDGIWPARSARRLHHHHQLAYHHHK--------QQQQQQAKQKQKQNGVQQGRSPTFMPVMLLLLMATLLTRPL---SAFSNRLSDTKLHEIYLDDKEIKLSWMVDWYKQEVLFHLQNAFNE-
QHRWFYLGFSKRGGLADADICFFENQNGFFNAVTDTYTSPDGQWVRRDYQQDCEVFKM----DEFTLAFRRKFDTCDPLDLRLHEGTMYVVWARGETELALEDHQFALPNVTAPHEAGVKMLQLLRADKILIPESELD--
HMEITLQEAPIPSQETTYWCHVQRLEGNLRRR-HHIVQFEPLIR--TPGIVHHMEVFHCEAGEHEEIPLYNGDC--
EQLPPRAKICSKVMVLWAMGAGTFTYPPEAGLPIGGPGFNPYVRLEVHFNNPEKQSGLVDNSGFRIKMSKTLRQYDAAVMELGLEYTDKMAIPPGQTAFPLSGYCVADCTRAALPATGIIIFGSQLHTHLRGVRVLTRHFRGEQELREVNRDDYYSNHFQ

EMRTLHYKPRVLPGDALVTTCYYNTKDDKTAALGGFSISDEMCVNYIHYYPATKLEVCKSSVSEETLENYFIYMKRTEHQHGV--HLNGARSSNYRSIE-WTQPRIDQLYTMYMQEPLSMQCNRSDGTRFEGRSSWEGVAATPVQIRIPIHRKL-
CPNYNPLWLKPLEKGDCDLLGECIY---------------------------------------------------------------------------------------------------------------------------------------
----------------------------------------------------------------------------------------------------------------------------------------------------------------
----------------------------------------------------------------------------------------------------------------------------------

>CAI9716180.1 dopamine beta-hydroxylase Octopus vulgaris
MLRFRLLSMDVVVVVVVVTAVNEDIFLFTKLETEMLRRPKSWLHYC-------------------------------------------------QVLLILLIVLKAAPKVTGRQHYKYNAY---LD----SQRRFHLWWDVDYKRNVVNFRLLAVVT--
LDEWFGVGYSAYGNCSNADFFVHWVDENYVQHCKDAWTDSKGI-LHIDNQQDYKILSGQLNKHALVIEYQRAFDTCDNYDFPFDNGTLHVVFFIGDNTFHYPE-GVNV-TSLSP---GLQRVLLLEAD-
VPIPRHPKDTWTFSVLAPGVTIPARETTYWWYTTKLPALPER--QHIIQFESAITKGNENLVHHIEVFLCEWKHGMKIPYFNGPSLVGEKPKELSVCRHVIGAWAMGASALHLPAEAGCAIGG-
NISRNILLEVHYNNPEMKHGVKDTSGIRFYVTPTRRKYDSGVMEIGLEYTNKMAIPPHQKSFILAGDCISECTELALPDGGIYIFASQLHTHLTGKKVTIRHIRNGTELPPVNYDDHYSPHFQIIKLLPRPVHILPGDSMVIECEYNTEKRPNATIGGFA
ITDEMCVSYLHYYPNTNLEVCKSSIETKTLNKWFEFIRRWDVDK-I--DLSRGYRNSYNSVR-WTPLTSHLLNKLYEISPLSMQCNQSDGRRFPG--NWEGMPQTEITLPLKPPERD-
CDCTLNNPEKDNRIRFGILNIQASKKKQLNWNEIVAESHMQCVVDDVQNLSHVSHRQYQAQFASGFFECPKMGLLQQTFILVGLLLCINYVPARRLVKRQSNELCGDYAEDFKKRHNEMCQMYYECNLGYLNDVQECAPGTVFNPSIQVCDYLNNTDCGN
LQVPKSAIEFGRGGGGNGGGNGGWNGGNGGGQGGGRYGQGGGNGGGGQGGGNGGGGQGGWNGGNGGNGGNRGGGGQGGWNGGNGGNRGGGGKGGWNGGNGGNRGGGQGGGGNYGQGGSGGNRGGQGGGRGGNQGTQWGNGGKRGGQWGGGGNRGGNRGGF
VNEDLRSLCSPYDYRLSNCSLAQHKGGKFGRLGYIGVFEASRKMTFRLNRNQRIAVVRYYYHCGSNAADASRRLSEKFYIHAVQGRNIKSLINKFEIKGYDVNNTPRSATSNEKVEQWKLVQSMVRINHS
>Cg XP 011429599.2 PREDICTED: dopamine beta-hydroxylase Crassostrea gigas
------------------------MNCIILNSNIGLTVGSITNFKC--------------------------VLSLGIKPGPPPSLSFSSMATFSSVSITLLVFFAQT--GTSLEIFKYKVE---LD----VNTNLVLRWGIDYNMARIYFELSATVL--
PGQGLMFGFSDYGDVTGADLVVLGTHFG-KVNLKDCWTDKNRI-LRMDFRQNYALTSGEWRNNKLLARFHRRFDTCDPMDYAIDTGTTHVIFAVTQGPPQP---GQQLPSSVKV---GLQRVQLLKPD-
LPQPDIPPDTWSFDILNPNITIPARDTTYWWFVTQLPQLPTK--
NHVIQYEGVITKGNEEFVHHIEVFHCQVDPLVKVTSYNGPGMAEGKSPELEACREVIGAWAMGASAIYLPNEAGTAIGGPHLSRYVLLEVHLNNPKLKTGVKDSSGIRFRVTRHLRPYDSGIMELGLEYTNKMAIPPGQSAFTLPGYCIPECTSIALPPN
GIRIYASQLHTHLTGKRVFTKHFRAGIELPELNRDNHYSPHFQEIRTLPRHVHVLPGDALVTSCVDSTANRSSITLGGFSIREEMCVNYVHYYPRCQLEVCKSSVADHSLHRFFQLANATQNSD--------DVAERYKSVD-
WTPSKVASLQKLYQHAPISMQCNTSSGVRFPG--YWSTKSVPRILLPRKSASTG-CVKDIPWVIP-----------------------------------------------------------------------------------------------
----------------------------------------------------------------------------------------------------------------------------------------------------------------
----------------------------------------------------------------------------------------------------------------------------------------------------------------
-------------------------

>Dm NP 788884.1 tyramine beta hydroxylase isoform B Drosophila melanogaster
---------------------------MLKIPLQLSSQDGIWPARFARRLHHHHQLAYHHHKQEQQQQQQQQQQQQAKQKQKQNGVQQGRSPTFMPVMLLLLMATLLT---RPLSAFSNRLSDTKLHEIYLDDKEIKLSWMVDWYKQEVLFHLQNAFNE-
QHRWFYLGFSKRGGLADADICFFENQNGFFNAVTDTYTSPDGQWVRRDYQQDCEVFKM----DEFTLAFRRKFDTCDPLDLRLHEGTMYVVWARGETELALEDHQFALPNVTAPHEAGVKMLQLLRADKILIPETELD--
HMEITLQEAPIPSQETTYWCHVQRLEGNLRRR-HHIVQFEPLIR--TPGIVHHMEVFHCEAGEHEEIPLYNGDC--
EQLPPRAKICSKVMVLWAMGAGTFTYPPEAGLPIGGPGFNPYVRLEVHFNNPEKQSGLVDNSGFRIKMSKTLRQYDAAVMELGLEYTDKMAIPPGQTAFPLSGYCVADCTRAALPATGIIIFGSQLHTHLRGVRVLTRHFRGEQELREVNRDDYYSNHFQ
EMRTLHYKPRVLPGDALVTTCYYNTKDDKTAALGGFSISDEMCVNYIHYYPATKLEVCKSSVSEETLENYFIYMKRTEHQHGV--HLNGARSSNYRSIE-WTQPRIDQLYTMYMQEPLSMQCNRSDGTRFEGRSSWEGVAATPVQIRIPIHRKL-
CPNYNPLWLKPLEKGDCDLLGECIY---------------------------------------------------------------------------------------------------------------------------------------
----------------------------------------------------------------------------------------------------------------------------------------------------------------
----------------------------------------------------------------------------------------------------------------------------------

>Dr NP 001103164.1 dopamine beta-hydroxylase Danio rerio
--------------------------MRLINKDLRLQDVTLMYLTV------------------------------------------------LATVVVLLVASYQAPAGQSRPTLAYHIP---LD----PSGQLELSWNISYPKQEVLLEVKVKE---
LHHGIILGMSDRGEPTNADLVILWDDGH-KSYFGDAWSDSEGR-VTLDTQQDYQLLETHQSTEGFFLLFKRSFSTCDPHDYIIEEGTVHIIYTMLEHPILSLH-ELNI-SRLQP---GVQRVLLLRPD-
TPSPHLPKDVRTLEVLAPDVIIPTQETTYWCHIYQLPPNLPK--NHIVMYESVITPGNEAIVHHIEVFEC-
SPQMDIVPQYSGSCDSKMKPRNLNYCRHVLAAWAMGAEPFYYPADAGLPMGGEGSSRFLRLEVHYHNPLLLSGRRDSSGIRLWYSPSLRRFDAGIMELGLVYTPVMAIPPRQRSFQLTGYCTAKCTQTALPVGGIHIFASQLHTHLAGLGVRTVLVRGGQ
EVEVVQEDKHFSTHYQIIRVLQKMVTVLPGDALLTTCRFNTEDRSKVTVGGFGIMEEMCVNYVHYYPRTQLELCKSHVDTDYLQKYFSLINRFQGRE-SCSCPQTSVEEQFSSVS-WDSFSGEVLNSLYVTAPFSMHCNQSTAQLFPG--
DWEKQEIPVVTAQLQRAPYP-CEINRRVASNNDSPTEVRADSSG--------------------------------------------------------------------------------------------------------------------
----------------------------------------------------------------------------------------------------------------------------------------------------------------
-------------------------------------------------------------------------------------------------------------------------------------------------------

>Hs NP 000778.3 dopamine beta-hydroxylase precursor Homo sapiens
---------------------MPALSRWASLPGPSMREAAFMYSTA------------------------------------------------VAIFLVILVAALQG-SAPRESPLPYHIP---LD----PEGSLELSWNVSYTQEAIHFQLLVRR---
LKAGVLFGMSDRGELENADLVVLWTDGD-TAYFADAWSDQKGQ-IHLDPQQDYQLLQVQRTPEGLTLLFKRPFGTCDPKDYLIEDGTVHLVYGILEEPFRSLE-AING-SGLQM---GLQRVQLLKPN-
IPEPELPSDACTMEVQAPNIQIPSQETTYWCYIKELPKGFSR--HHIIKYEPIVTKGNEALVHHMEVFQC-
APEMDSVPHFSGPCDSKMKPDRLNYCRHVLAAWALGAKAFYYPEEAGLAFGGPGSSRYLRLEVHYHNPLVIEGRNDSSGIRLYYTAKLRRFNAGIMELGLVYTPVMAIPPRETAFILTGYCTDKCTQLALPPSGIHIFASQLHTHLTGRKVVTVLVRDGR
EWEIVNQDNHYSPHFQEIRMLKKVVSVHPGDVLITSCTYNTEDRELATVGGFGILEEMCVNYVHYYPQTQLELCKSAVDAGFLQKYFHLINRFNNED-VCTCPQASVSQQFTSVP-WNSFNRDVLKALYSFAPISMHCNKSSAVRFQG--
EWNLQPLPKVISTLEEPTPQ-CPTSQGRSPAGPTVVSIGGGKG---------------------------------------------------------------------------------------------------------------------
----------------------------------------------------------------------------------------------------------------------------------------------------------------
-------------------------------------------------------------------------------------------------------------------------------------------------------

>KAI8741828.1 dopamine beta-hydroxylase-like Biomphalaria glabrata
-----------------------------------MRRCHHALSSI------------------------------------------------IGCLLIFKMVKWAS-------AFQYKLH---LD----SNGRYLFEWSVDYQKNNVNVQITAKVT--
QDIWFAVGFSDYGSVTSADLVVFWTDGHGNHHFVDGYTNDHGI-ILPDDQQDYFLTSVAADHGAVVLDFYRRFNTCDEQDYTLDNGTTHVIFFESVYANELPF-GHDV-SSLGH---AVKRVQLLKPE-
LPIPVLPNDTWTFEVRAPEILIPSVETTYWWHTTILPHIPTK--
HHIVKYSGVISEGNEDLVHHMEVFHCQVPKGQKIPYYSGPAENEDTPKGLEPCRRVIAAWAMGAQDMVYPEEAGVSIGGQDTSRFALLEVHYNNPERKSGRMDNSGIRFYVTSQLRKNEAGIMELGLEYTNKMAIPPGQKKFELSGYCVQDCTRVGLPPG
GIYVFASQLHTHLTGRRLYTKHVRQGSELPELNRDNHFSPHFQEIRRLPQPHNVLPGDELVTTCEYDTRSRDKATIGGFSITDEMCVNYIHYYPRTELEVCKSSIQTNALQTFFFLLNRLEDAK-V--YPDAGDRANYVNIN-
WTPVNAQLLDELYATSPLSMQCNRSDGVRFPG--DWEHQKVSEIAQPLVVRDIDTCPSS-----------------------------------------------------------------------------------------------------
----------------------------------------------------------------------------------------------------------------------------------------------------------------
----------------------------------------------------------------------------------------------------------------------------------------------------------------
-------------------------

>Mm NP 620392.2 dopamine beta-hydroxylase Mus musculus
-----------------MQAHLSHQPCWSSLPSPSVREAASMYGTA------------------------------------------------VAIFLVILVAALRG-SEPPESPFPYHIP---LD----PEGILELSWNVSYVQEIIHFQLQVQG---
LRAGVLFGMSDRGEMENADLIMLWTDGD-RAYFADAWSDRKGQ-IHLDSQQDYQLLQAQRTRDGLSLLFKRPFVTCDPKDYVIEDDTVHLVYGILEEPFQSLE-AINT-SGLHT---GLQRVQLLKSE-

```
VPTPSMPEDVQTMDIRAPDILIPDNETTYWCYITELPPRFPR--HHIIMYEAIVTEGNEALVHHMEVFQC-
AAESEDFPQFNGPCDSKMKPDRLNYCRHVLAAWALGAKAFYYPKEAGVPFGGPGSSPFLRLEVHYHNPRKIQGRQDSSGIRLHYTATLRRYDAGIMELGLVYTPLMAIPPQETAFVLTGYCTDKCTQMALQDSGIHIFASQLHTHLTGRKVVTVLARDGQ
ERKVVNRDNHYSPHFQEIRMLKKVVTVYPGDVLITSCTYNTENKTLATVGGFGILEEMCVNYVHYYPQTELELCKSAVDDGFLQKYFHMVNRFSSEE-VCTCPQASVPQQFSSVP-WNSFNRDMLKALYDYAPISMHCNKTSAVRFPG--
EWNLQPLPKITSTLEEPTPR-CPIRQTQSPANPTVPITTEADAE-------------------------------------------------------------------------------------------------------------------------
---------------------------------------------------------------------------------------------------------------------------------------------------------------------
-----------------------------------------------------------------------------------------------------------------------------------------------------------
>NP 001243923.1 tyramine beta hydroxylase precursor Bombyx mori
---------------------------------------MSNWFFVF-------------------------------------------------FLSVYTFIYTFSVNV-----SPGHRFAESESVLD----
PSGEFLLKWRVDYAVRKIKFTLTVSEKAPAFNWFALGFSDRGQLNNSDVCLFWTDYKARDHFEDMHTNGQGN-LIRDQKQNCEGFYLNT--NSQSIIFDRLFDTCDDDDYVIEDGTVHVVWAHGIDKLFSSK-GLCL-TCTVPQRHGFVRVRLL----
TPPGLQKANGYQLRITNKDLKVPGDDTTYWCKVVRLPEFVTSKVHHIVQFESTITPGNEGLVHHMEVFYCDEDPHKEMPAYEGNCFAAERPAITKSCSKVKAAWAMGAPPFTYPKEAGLPLGGPKANKYVMLEVHYNNPELRKDWVDSSGIVLYITGRKR
TYDAAIMELGLEYTDKMAIPGEQKAFPLTGYCIPQCTGVGLPDEGITVFGSQLHTHLTGAAVWTRHSRQGVELPVLNKDMHYSTHFQEIRILHRPVKVLPGDFLETTCIYNTEDKLNATIGGHAITDEMCVNYIHYYPATELEVCKSAVSNEALEKYFDF
EKRWDDIP-I--SSKATPRDNYLAIQPWTRLRADTLHTLYVESPISMQCNKSDGSRFQG--DWEGIPIPKIKLPLSEEIRI-CPKINYLVET----------------------------------------------------------------------
---------------------------------------------------------------------------------------------------------------------------------------------------------------------
---------------------------------------------------------------------------------------------------------------------------------------------------------------------
------------------------------------------------------
>XP 052825887.1 dopamine beta-hydroxylase Octopus bimaculoides
-----------------------------------MLRRLKTWLHYC---------------------------------------------------QVLLVLLIVLKAAPKVTGRQHYKYNAY---LD----SQRRFHLWWDVDYKRNVVNFRLLAVVT--
LDEWFGVGYSAYGNSSNADFFVHWVDENYVQHCKDAWTDSKGI-LHIDNQQDYKILSGQLNKHALVIEYQRPFDTCDNYDFPFD-----------------------------------------------------------------
VTIPAKETTYWWYTTKLPALPER--QHIIQFESAITKGNENFVHHIEVFLCEWKHGMKIPYFNGPSLVGEKPKELSVCRHVIGAWAMGASALHLPAEAGCAIGG-
NISRNILLEVHYNNPEIKHGVKDTSGIRFYVTPTRRKYDSGVMEIGLEYTNKMAIPPHQNSFILAGDCISECTELALPDGGIYIFASQLHTHLTGKKVTIRHIRNGTELPPVNYDDHYSPHFQIIKLLPRPVHILPGDSMVIECEYNTEKRSNATIGGFA
ITDEMCVSYLHYYPNTNLEVCKSSIETKTLNKWFEFIRRWDVDK-I--DLSKGYRNSYNSVR-WTPLTSHLLNKLYEISPLSMQCNQSDGRRFPG--NWEGMPQTEITLPLKPPERD-CDCTLNNPEKDNVIA----------------------------
---------------------------------------------------------------------------------------------------------------------------------------------------------------------
---------------------------------------------------------------------------------------------------------------------------------------------------------------------
-------------------------------------------------------------------------------------------

Tyramine beta hydroxylase
>O.bocki_TBH
-------------------------------------------------------------------------------------------------------------AAPKVTGRQHYKYNVY---LD----SQRRFHLWWDVDYKRNVVNFRLLAVVT--
LDEWFGVGYSAYGNSSNADFFVHWVDENYVQHCKDAWTDSKGI-LHIDNQQDYKILSGKLNKHALVIEYERPFDTCDNYDFPFDNGTLHVVFFIGDNTFHYPE-GVNV-TSLSP---GLQRVLLLEAD-
VPVPRHPKDTWTFSVLAPGVTIPAKETTNWWYTTKLPALNER--QHIIQFESAITKGNKNFVHHIEVFLCEWKRGMKIPYFNGPSMVGEKPKELSVCRHVIGAWAMGASALHLPAXAGCAIGG-NISRDILLEVHYNNPEMKHGVXDTSGI---------
------------------------------------SXMLXLHEG--------------------------------------------------------------------------------------------------------------STILESW---
------------------------KLX-W----------------------------------------------------------------------------------------------------------------------------------------
---------------------------------------------------------------------------------------------------------------------------------------------------------------------
---------------------------------------------------------------------------------------------------------------------------------------------------------------------
------------------------------------------------------
>XP 029645364.1 dopamine beta-hydroxylase Octopus sinensis
-----------------------------------MLRRPKSWLHYC---------------------------------------------------QVLLILLIVLKAAPKVTGRQHYKYNAY---LD----SQRRFHLWWDVDYKRNVVNFRLLAVVT--
LDEWFGVGYSAYGNCSNADFFVHWVDENYVQHCKDAWTDSKGI-LHIDNQQDYKILSGQLNKHALVIEYQRAFDTCDNYDFPFDNGTLHVVFFIGDNTFHYPE-GVNV-TSLSP---GLQRVLLLEAD-
VPIPRHPKDTWTFSVLAPGVTIPARETTYWWYTTKLPALPER--QHIIQFESAITKGNENLVHHIEVFLCEWKHGMKIPYFNGPSLVGEKPKELSVCRHVIGAWAMGASALHLPAEAGCAIGG-
NISRNILLEVHYNNPEMKHGVKDTSGIRFYVTPTRRKYDSGVMEIGLEYTNKMAIPPHQKSFILAGDCISECTELALPDGGIYIFASQLHTHLTGKKVTIRHIRNGTELPPVNYDDHYSPHFQIIKLLPRPVHILPGDSMVIECEYNTEKRPNATIGGFA
ITDEMCVSYLHYYPNTNLEVCKSSIETKTLNKWFEFIRRWDVDK-I--DLSRGYRNSYNSVR-WTPLTSHLLNKLYEISPLSMQCNQSDGRRFPG--NWEGMPQTEITLPLKPPERD-CDCTLNNPEKDNVIA----------------------------
---------------------------------------------------------------------------------------------------------------------------------------------------------------------
---------------------------------------------------------------------------------------------------------------------------------------------------------------------
-------------------------------------------------------------------------------------------
>XP 974169.1 PREDICTED: tyramine beta-hydroxylase Tribolium castaneum
--------------------------------------------MRF---------------------------------------------------LIAIIILTCLKSS-----HSGEIFHVP---LN----GDGSISLNWVLDYPTQTVTFEVHLPEN--
FG-WFAIGFSDQGAHFPADYCILWKTIKRKIQFEDTWADTTGI-IRLDRQQDCQNFKIKRAGNVTKFTFRRKFDTCDFEDYVIEDGTTHIVWARGAHPLYKVV-GLNISSPEKEQ--GMVRVQLLKNT-
NVKAILPNYVQTLDVFAHEVRVPDKETTYWCHVHKLGEEFKEK-
HHVYRYEAHIPSSSEGLVHHMEVFHCVAPPNQQIPLYVGNCFAKDRPKETQVCKRVLAAWAMGAPPFTYPEEAGLPLGGPDFNPYVMLEVHYNNPEHKTGFVDSSGIRFHVSSKLKKMDAGVIELGLEYTDKMAIPPGQEAFPLTGYCVSECTAVSLPPE
GITIFGSQLHTHLTGVKVYTRHIRDGIELRELNRDDHYSTHFQEIRRLKQPVKVLPGDALVTRCYYNTQERENITLGGFSITDEMCVNYVHYFPATQLEVCKSAISDQALSTYFNYMKEWEGQK-I--SLHHVISDNYKSIK-
WNKMRVQLLSDVYHEAPLSMQCNMSSGDRFPG--YWENTPITPILTPLPPPPRT-C-----------------------------------------------------------------------------------------------------------
---------------------------------------------------------------------------------------------------------------------------------------------------------------------
---------------------------------------------------------------------------------------------------------------------------------------------------------------------
--------------------------
>XP 011429599.3 dopamine beta-hydroxylase Crassostrea gigas
------------------------MNCIILNSNIGLTVGSITNFKC----------------------------VLSLGIKPGPPPSLSFSSMETFSSVSITLLVFFAQT--GTSLEIFKYKVE---LD----VNTNLVLRWGIDYNMARIYFELSATVL--
PGQGLMFGFSDYGDVTGADLAVLGTHFG-KVNLKDCWTDKNGI-LRMDFRQNYALTSGEWRNNKLLARFHRRFDTCDPMDYAIDTGTTHVIFAVTQGPLHP---GQQLPSSVKV---GLQRVQLLKPD-
LPQPDIPPDTWSFDILNPNITIPARDTTYWWYVTQLPQLPIK--
NHVIQYEGIITKGNEEFVHHIEVFHCQVDPWVKVTSYNGPGMAEDKSPELEACREVIGAWAMGASAIYLPNEAGTAIGGPHLSRYVLLEVHLNNPKLKTGVKDSSGIRFRVTRHLRPYDSGIMELGLEYTNKMAIPPGQSAFTLPGYCIPECTSIALPPS
GIRIYASQLHTHLTGKKVFTKHFRAGIELPELNRDNHYSPHFQEIRTLPRHVHVLPGDALVTSCVDSTANRSSITLGGFSIREEMCVNYVHYYPRCQLEVCKSSVADHSLHRFFQLANATQNSD--------DVAERYKSVD-
WTPSKVASLQKLYQHAPISMQCNTSSGVRFPG--YWSTKSVPRILLPRKSASTG-CVKDIPWVIP--------------------------------------------------------------------------------------------------
---------------------------------------------------------------------------------------------------------------------------------------------------------------------
```

--------------------------------------------------------------------------------------------------------------------------------------------------------------------------------
-------------------------
>XP 021341155.1 dopamine beta-hydroxylase-like Mizuhopecten yessoensis
----------------------------------------MNFLTRV---------------------------------------------------LSFGLLLQTALSLP-------SYRYHTT---LD----
TGGQLTLDWDVDRVQQKVKFRLTAKLDHHKNRWFGLGFSDYGMITDADFAVYWTDTR-VHHFQDCWTDDRGL-LHVDRHQDFILMTSSIDKGRRILEFQRRYDTCDAHDYIIENGTTHILFFIGESSPQSLE-GVAL-SDLGV---QITRVQLLKPE-
IPVPVYPSDTWIFDVTTPSITVPDTDTTYWWYLTRLPHLSDK--
HHIIKYESVIQRGSEHLVHHMEVFQCEIHPDDDIKPYNGPGMAEGKPPELSTCRNVIGAWAMGSGPFILPPDAGIPIGERSASSFILLEIHYNNPNLLKGIVDSSGIRFYVTRHLRRHDAGIMELGLEYTNKMAVPPGQTSFPLRGYCIAECTKVGLGHH
SIHIYGSQLHTHMTGRKVYTKHLREGVELPELNRDNHYSPHFQEIRRLPVPVEIRPGDALVTTCDYHTEDRPNVTVGGFSIKDEMCVNYVHYYPRTNLEVCKSSIQTSSLHKFFSFLNVWDKAD-T--SQTKGDRDNYSSIR-
WSTTTTKLLLSLYDMAPLSMQCNKSDGTRFPG--VWESAQKPLIYLPLPKPKSR-CRRLRLLN-------------------------------------------------------------------------------------------------------
--------------------------------------------------------------------------------------------------------------------------------------------------------------------------------
--------------------------------------------------------------------------------------------------------------------------------------------------------------------------------
-------------------------
>Xt XP 002942324.2 PREDICTED: dopamine beta-hydroxylase Xenopus tropicalis
----------------------MAKRRFCSFSNLRIREVVPVYFTM----------------------------------------------------LAVFMILLVASLQGPSRQQKSSFPYQVP---LD----PQGSLQLYWNVSYSEKVVYFRILIKD---
LKFGLIFGMSDRGDFEDADMAVLWTNEL-GSFFGDAWSDQKGQ-LHMDSQQDYELLNAQQSQEGLYLLFRRPFATCDPKDYLIEDGTVHLIYALLEKPFSTLS-SIDV-SAIKNR--GLQRVQLLKPD-
LPIPRLPADVLNMEVRAPQVLIPAKETTYWCHITELPKDFTK--HHIVMYEPVITKGHEAIVHHIEVFQC-
ASNYYTIPRYDGPCDSKMKPQSLNSCRHVLAAWAMGAKAFYYPEESGLAFGGPDSSRYLRLEVHYHNPLELKGLRDSSGIRLYYTSTLRRYDAGIMEVGLVYSPVMAIPPGQKDFLLTGYCTDKCTDRALPSNGIKIFASQLHTHLGGRGVPPILVREGK
EAEVVNADGHYSPHFQEIRMLKKAVHVLPGDVMVTSCSYNTEDRKNITVGGFSITDEMCVNYVHYYPRTDLELCKSMVDPGYLQKYFHMVNRFNSDD-VSTSPNTTVTKQFQEVP-WNSFSAGVLKSLYSFAPISMHCNRSSAVRFPG--
EWEKQPLPQINEKLPSPPAT-CPAMDPAKSKGPTLVRLLQ--------------------------------------------------------------------------------------------------------------------
--------------------------------------------------------------------------------------------------------------------------------------------------------------------------------
--------------------------------------------------------------------------------------------------------------------------------------------------------------------

Excitatory Amino Acid Transporter 1
>Obocki
---------------MENESSNSGKVNTHVFVSSPDEDTSRAGHLKKFGKQFGKRCLSAGKE---------------
NLLLILLLGSVIFGCAVGFLVRAVRPSMMTNKREVMYLMFPGEMLMRMLRMLILPLIMSSLISGIAGLDAKTCGKMGLRTIGYFASTTIISVVTGIIMCVTIQPGSGKSSEKLKRYGSTKRVNSVDTFLDLIRNMFPDNIITSCFDSFRTEEVL-----
-----EEPDVVENATDILTSTIST--LALSTQNLTFMNDTMKQV-----
YVPKAGSTGQTNVLGIVVFSVLMGITLGRMAIRGKPLLDVCNCINEATMKLFKIFIWYSPLGITFLIAAKIVEMRDFGVLLGKVGLYFITVLCGLAVHGACILPALYFLCVRKNPYKFIAGIFQAMATAFGTSSSSATLPVTLHCLEHNCGVDPRVSSFV
IPVGATINMDGTALYEAVAALFIAQVNNASMDFG----------------------------------------------------------------------------------------------------------------
>XP_029641976.1 excitatory amino acid transporter 1 Octopus sinensis
---------------MENESSNSGKVNTHVFVSSPDEDTSRAGHLKKFGKQFGKRCLSAGKE---------------
NLLLILLLGSVIFGCAVGFLVRAVRPSMMTNKREVMYLMFPGEMLMRMLRMLILPLIMSSLISGIAGLDAKTCGKMGLRTIGYFASTTIISVVTGIIMCVTIQPGSGKSSEKLKRYGSTKRVNSVDTFLDLIRNMFPDNIITSCFDSFRTEEVL-----
-----EEPPVVENATDIFTSTISS--LALSTQNLTFMNDTVKQV-----
YVPKAGTTGQTNVLGVVVFSVLMGVTLGRMAIRGKPLLDVCNCINEATMKLFKIFIWYSPLGITFLIAAKIVEMRDFGVLLGKVGLYFITVLCGLAVHGACILPALYFLCVRKNPYKFIAGIFQAMATAFGTSSSSATLPVTLHCLEHNCGVDPRVSSFV
IPVGATINMDGTALYEAVAALFIAQVNNLSMDFGQIVTISITATAASIGAAGVPQAGLVTMVIVLSAVGLPTDDITLILVVDWFLDRFRTMINVEGDSLGAGIVYHLSKQELAVAEKEGEEGALTPFTGIDDKKPNGTDS---------AAITAV
>XP_052826793.1 excitatory amino acid transporter 1 Octopus bimaculoides
---------------MENESSNSGKVNTHVFVSSPDEDTSRAGHLKKFGKQFGKRCLSAGKE---------------
NLLLILLLGSVIFGCAVGFLVRAVRPSMMTNKREVMYLMFPGEMLMRMLRMLILPLIMSSLISGIAGLDAKTCGKMGLRTIGYFASTTIISVVTGIIMCVTIQPGSGKSSEKLKRYGSTKRVNSVDTFLDLIRNMFPDNIITSCFDSFRTEEVLE----
-----EPDVVAENATDIFTSTISS--LALSTQNLTFMNDTVKQV-----
YVPKAGSTGQTNVLGVVVFSVLMGVTLGRMAIRGKPLLDVCNCINEATMKLFKIFIWYSPLGITFLIAAKIVEMRDFGVLLGKVGLYFITVLCGLAVHGACILPALYFLCVRKNPYKFIAGIFQAMATAFGTSSSSATLPVTLHCLEHNCGVDPRVSSFV
IPVGATINMDGTALYEAVAALFIAQVNNLSMDFGQIVTISITATAASIGAAGVPQAGLVTMVIVLSAVGLPTDDITLILVVDWFLDRFRTMINVEGDSLGAGIVYHLSKQELAAVEKEGEEGVLTPFTGIDDKKPNGTDS---------AAITAV
>XP_009046107.1 hypothetical protein LOTGIDRAFT_61417 partial Lottia gigantea
------------------------------------------------------------KE---------------NLLLVLLFCGVFLGVGMGFMVRSIGGDF--
SKRQIMYLMFPGEILMRMLKMLILPLIVASLIAGIAGLDAKTCGKMGLRTIAYFACTTILAVILGIIMAVSIRPGGGAEASEIKRYGAAKRVNSVDTFLDLIRNLFPDNLILSCVDAYRTVEKKEF---------
KEKIMPKTNDTLNTTASPTSLVYYLLNNATNVTEKVGEPYE---
WVPTKSSTGQTNVLGIVMFSILFGVMLGRMDSRGKPVLDFCSVIVEVTMKLFTVFIWYSPVGIAFLIAAKIVEMEDFSVLVGKVGLYFITVLSGLFVHGSVVLPGLFFLGSRQNPYKFIYGISQAMATAFGTSSSSATMPVTLHCLEHNNHVDPRVSSFV
IPVGATINMDGTALYEAVAALFIAQVNNMDMDFGQIVTISITATAASIGAAGVPQAGLVTMVIVLSAVGLPVNDITLILVVDWFLDRFRTMTNVMGDSLGAGLVYHLSKHEL-------------------------------------------
>XP_055880052.1 excitatory amino acid transporter 1-like isoform X1 Biomphalaria glabrata
---MERKRSSNADGRDGVGENNVNPVNTHVFITVPEEDLGFSGNLKKYG----KKCIQGSKQ---------------NALLIGLLLSVVMGILLGLVIRSTKDKF--
SKREVMYLGFPGELLMRMLQMLILPIILSSLISGIAGLDGKTCGKMGLRTIGYFTVTTFMAVFLGILLAMTIQPGGNADRSSMRRYGKAEPLNSADTFLDLIRNVFPDNLIVTCFNAYRTKEVRVPIM-------SIPVVNLSATTTSTITTET--
MTTESPLATTLNETEEVQYD---
ISLLNEGTGKPNVLGIVTFAILFGIMLGRMGERGKPILAFCDCLVEVTMKLFTFFLWYSPFGIAFLIAAKIVEMEDFSVLLGKVGMYFITVLIGLFIHGSIVLPLIYFVLVRKNPYTFIYGISQALATAFGTSSSSATMPVTLKCLEHNNHVDSRVASFV
IPVGATINMDGTALYEAVAALFIAQVNNQSMNFGNIITISITATAASVGAAGVPQAGLVTMVIVLAAVGLPIDDVTLILVVDWFLDRFRTMTNVMGDSLGAGIVHHYSVDEMPPLEHDLHDHEMEKLPNGQQ-----------------QETTPV
>XP_011419763.2 excitatory amino acid transporter 1 Crassostrea gigas
MDYKPPKPVVDPESKLLCSNDTSAKYDTQVLYENEG------SYQQRHGTNMLHLCVKGLKQ---------------NLILVLLLLSVVFGCAIGFGIRATGQLT---
KREIMYLQFPGEMLMRMLKMLIIPLVVASLIGGIAGLDAKTCGKMGLRTMAYFGTTTMLAVILGIIMAVTIRPGSNGDQTEIPRYGEAKKLNSVDTFLDLIRNMFPDNLIVACYTGYRTDLNEKK---------TVKNVSLINENSTSEMMNQ--
TFELKETVT--------------
LEPSKGETGQSNILGLVVFSVCFGIVIGRMGDKGKPLLNFCNCVTDCTMKLFVVFIWYSPIGITFLIAGKIVGMEDFGVLMTRVGLYFLTVLFGLTIHGGVLLPLLYFVITRRNPYVFLRGILEAMATAFGTSSSSATMPVTLRCLETKNGIDQRVSSFV
VPVGATINMDGTALYEAVAALFIGQVNDIDMSFGQIVTISITATAASIGAAGVPQAGLVTMVIVLAAVGLPIDDVTLILVVDWFLDRFRTMTNVVGDSIGAGIVYHLSKDELSGYNEDKTDNLNLVRVDIPNGK---------------DAITAV
>XP_012940305.2 excitatory amino acid transporter 1 partial Aplysia californica
-MDSQKGPVDSRDGPQSNTDNNVSPVNTHVFITVPEEDLGFRGNLKKYG----RKCLAGSKE---------------NALLIGLLLSVVGGVVIGVVARSLKDKY--
SKREIMYLGFPGEILMRMLQMLILPIILSSLISGIAGLDAKTCGKMGLRTIGYFGTTTFLAVFLGIVLAVTIQPGGNTDLSEMKRYGKAEQLNSADTFLDLVRNVFPDNLIVACYNAYRTKEVEKEEFREATTNSTASITTLSPEPTTTLSPRS--

ITTLSPSSTTIPENGTELVAVTVIELVNGGTGKPNVLGVVMFSILFGVMLGRMGERGKPIVHFCDCLVEVTMRLFTMFLWYSPVGIAFLIAAKIVEMEDFNVLLGKLGMYFITVLIGLFIHGSIILPFLYFITVRKNPYKFIYGISQAMATAFGTSSSSA
TMPVTLRCLEHNNHVDARVASFVIPVG-------------------------------------------------------------------------------------------------------------------------------------
------------------
>NP_001160167.1 excitatory amino acid transporter 1 isoform 2 Homo sapiens
------------------------------MTKSNGEEPKMGGRMERFQQGVRKRTLLAKKKVQNITKEDVKSYLFRNAFVLLTVTAVIVGTILGFTLRPYRMSY----
REVKYFSFPGELLMRMLQMLVLPLIISSLVTGMAALDSKASGKMGMRAVVYYMTTTIIAVVIGIIIVIIIHPGKGT-KENMHREGKIVRVTAADAFLDLIRNMFPPNLVEACFKQFKTNYEKR----------SFKVPIQANETLVGAVINN--
VSEAMETLTRITEE---------
LVPVPGSVNGVNALGLVVFSMCFGFVIGNMKEQGQALREFFDSLNEAIMRLVAVIMWYAPVGILFLIAGKIVEMEDMGVIGGQLAMYTVTVIVGLLIHAVIVLPLLYFLVTRKNPWVFIGGLLQALITALGTSSSSATLPITFKCLEENNGVDKRVTRFV
LPVGATINMDGTALYEALAAIFIAQVNNFELNFGQIITIR----------------------------------------------DRLRTTTNVLGDSLGAGIVEHLSRHELKNRDVEMGNSVIEENEMKKPYQLIAQDN--ETEKPI-DSETKM
>AAH37310.1 Solute carrier family 1 (glial high affinity glutamate transporter) member 3 Homo sapiens
------------------------------MTKSNGEEPKMGGRMERFQQGVSKRTLLAKKKVQNITKEDVKSYLFRNAFVLLTVTAVIVGTILGFTLRPYRMSY----
REVKYFSFPGELLMRMLQMLVLPLIISSLVTGMAALDSKASGKMGMRAVVYYMTTTIIAVVIGIIIVIIIHPGKGT-KENMHREGKIVRVTAADAFLDLIRNMFPPNLVEACFKQFKTNYEKR----------SFKVPIQANETLVGAVINN--
VSEAMETLTRITEE---------
LVPVPGSVNGVNALGLVVFSMCFGFVIGNMKEQGQALREFFDSLNEAIMRLVAVIMWYAPVGILFLIAGKIVEMEDMGVIGGQLAMYTVTVIVGLLIHAVIVLPLLYFLVTRKNPWVFIGGLLQALITALGTSSSSATLPITFKCLEENNGVDKRVTRFV
LPVGATINMDGTALYEALAAIFIAQVNNFELNFGQIITISITATAASIGAAGIPQAGLVTMVIVLTSVGLPTDDITLIIAVDWFLDRLRTTTNVLGDSLGAGIVEHLSRHELKNRDVEMGNSVIEENEMKKPYQLIAQDN--ETEKPI-DSETKM
>NP_683740.1 excitatory amino acid transporter 1 Mus musculus
------------------------------MTKSNGEEPRMGGRMERLQQGVRKRTLLAKKKVQSLTKEDVKSYLFRNAFVLLTVTAVIVGTILGFALRPYKMSY----
REVKYFSFPGELLMRMLQMLVLPLIISSLVTGMAALDSKASGKMGMRAVVYYMTTTIIAVVIGIIIVIIIHPGKGT-KENMYREGKIVQVTAADAFLDLIRNMFPPNLVEACFKQFKTSYEKR----------SFKVPIQSNETLLGAVINN--
VSEAMETLTRIREE---------
MVPVPGSVNGVNALGLVVFSMCFGFVIGNMKEQGQALREFFDSLNEAIMRLVAVIMWYAPLGILFLIAGKIVEMEDMGVIGGQLAMYTVTVIVGLLIHAVIVLPLLYFLVTRKNPWVFIGGLLQALITALGTSSSSATLPITFKCLEENNGVDKRITRFV
LPVGATINMDGTALYEALAAIFIAQVNNFDLNFGQIITISITATAASIGAAGIPQAGLVTMVIVLTSVGLPTDDITLIIAVDWFLDRLRTTTNVLGDSLGAGIVEHLSRHELKNRDVEMGNSVIEENEMKKPYQLIAQDN--EPEKPVADSETKM
>NP_001177232.1 solute carrier family 1 member 3b Danio rerio
------------------------------MTKSTGEKPRSRSRVQQFREGIQLRSLKARKKVEDISKDDVKGFLRRNAFVLFTIGAVVFGITLGFALRSYKMSY----
REVKYFSFPGELLMRMLQMLVLPLLVSSLITGMAALDSRASGKMGMRAVIYYMTTTFIAVFIGIIMVLIIHPGKGS-KDEFTKQQKIEQVSPADAFLDLIRNMFPPNLVQACTQQFKTQYGKR----------IIHVKMIVNDSIFNLTNAT--
QEIAQEE----------------
VIPLSGTTNGVNALGLVVFSMCFGLIIGNMKEQGQALRDFFDSLNEAIMRLVAIIMWYAPIGILFLIAGKIVEMDDITAMGGQLGMYTVTVIIGLMIHGIIILPTLYFVITRKNPFIFITGLLQALITALGTSSSSATLPITFKCLEENNKVDKRVTRFV
LPVGATINMDGTALYEALAAIFIAQVNNMEMNFGQIITISITATAASIGAAGIPQAGLVTMVIVLTSVGLPTDDITLIIAVDWFLDRLRTTTNVLGDSIGAGIIEFLSKDELQGGDVELGSSVLEENELKKPYQKIPQENEYENEKPP-DSETKM

**FXRI (FLRIamide)**
>ObockiFLRI
----------------------------------------L-----LKKMAGLWRIVLLATITLMSVTSQWV-------IYVQA-----------------------EEASNNDEL-----VNSANQ--------------------ADPE----
MDSENDL-KRANTFLRIGKAN-------GVLRLARS---------PSSFLRIGRRN---------Q----LRFIRIGKAPSSM-------FLRIGKRVDDSENDDLNEYDADKMSGRDIRA--SPS--------------SFLRIGKSGNEVEDE-----
----IDDTD-EIAKRVNAFLRIGRQ-----------NDPSSFLRIGKSLNNE------DLSKD-----------------------KRTNAFLRI-------------------------------------GKIPTSSFIRLGRGP--
FTEDNGINTRGFRGPTHG-----FLRIGKRAANTDGS---HTDYFSDLNVK--
>CAI9719694.1 Hypothetical predicted protein Octopus vulgaris
-MQFDLLKRKTPHTQLLMYLPSHAVSKINNFSAWNSNRRIL-----LKKMAGLWRIVLLATITLMCVTSQWV-------IYVQA-----------------------EEASSSDEL-----VNSANE--------------------AEPE----
MDSEDDL-KRANTFLRIGKAN-------GILRLARS---------PSSFLRIGRRP---------P----LHFVRIGKAPSSM-------FLRIGKRVDDSENEDLNDYDADKMSGRDTRA--SPS--------------SFLRIGKSGNEVEDE-----
----IDDTD-ETVKRVNAFLRIGRQ-----------NDPSSFLRIGKSLNNE------DLSKD-----------------------KRTNAFLRI-------------------------------------GKIPASSFIRLGRGP--
FTEDNGINTRGFRGPTRG-----FLRIGKRAAIPDGS---HADYFSDLNVKSQ
>XP 014791297.1 FMRFamide neuropeptides Octopus bimaculoides
-------------------------------------------------MAGLWRIVLLATITLMCVTSQWV-------IYVQA-----------------------EEASSSDEL-----VNSANE--------------------ADPE----
MDSEDDL-KRANTFLRIGKAN-------GILRLARS---------PSSFLRIGRRP---------P----LHFVRIGKAPSSM-------FLRIGKRVDDSENDDLNEYDADKMSGRDTRA--SPS--------------SFLRIGKSGNEVEDE-----
----IDDTD-ETVKRVNAFLRIGRQ-----------NDPSSFLRIGKSLNND------DLSKD-----------------------KRTNAFLRI-------------------------------------GKIPASSFIRLGRGP--
FTEDNGINTRGFRGPTRG-----FLRIGKRAAIPDGS---HADYFSDLNVKSQ
>XP 029634231.1 FMRFamide neuropeptides-like Octopus sinensis
-------------------------------------------------MAGLWRIVLLATITLMCVTSQWV-------IYVQA-----------------------EEASSSDEL-----VNSANE--------------------AEPE----
MDSEDDL-KRANTFLRIGKAN-------GILRLARS---------PSSFLRIGRRP---------P----LHFVRIGKAPSSM-------FLRIGKRVDDSENEDLNDYDADKMSGRDTRA--SPS--------------SFLRIGKSGNEVEDE-----
----IDDTD-ETVKRVNAFLRIGRQ-----------NDPSSFLRIGKSLNNE------DLSKD-----------------------KRTNAFLRI-------------------------------------GKIPASSFIRLGRGP--
FTEDNGINTRGFRGPTRG-----FLRIGKRAAIPDGS---HADYFSDLNVKSQ
>CAE1160964.1 unnamed protein product Sepia pharaonis
-------------------------------------------------------------------MTQWA-------SFVRA-----------------------ESPSNGEDL-----VNAAGA---------AVESA------
DEPSGRSVSDSPYDM-KRANQFLRIGRGS-------HFIRIGRG--------GASSFLRIGRNP---------L----SQFVRIGKAPSSM-------FLRIGKS-SAAGNPEVG----------DLAA--GPS--------------SL----------
GDS---------NIDED-EVLKRASSFLRIGRS-----------N-PSTFLRIGKSAGNLDE----DTAAN-------EEIVTDDIDVPSESVAKRANAFLRI-------------------------------------GKIPASSFVRIGRGP--
YGIDNRSNPRGFLSVGSR-----FVRIGKREAIPSETGPVHARLLPNLHDQAQ
>KAI8760658.1 FMRF-amide neuropeptides-like Biomphalaria glabrata
MKTIMFINIILISVTVLLHGVSGDIAEDNLEDDKRASSFVR-----IGRPSSFVRIGRGDNVEDLETDPNYVDLEKKASNFVRIGRYPTMSRFIRIGRTPMEGAGSYEDDSSEEEP-----IGDDGKRASSFVRIGKRKSSFVRIGKSLAE----
EDSDEDVDKRASSFVRIGKSPS------SFVRIGKA---------PSSFVRIGKSPSSFVRIGKSP----SSFVRIGKVPSS--------
FVRIGRSIENDLKSELDDIDEEKKASSFVRIGKSPNEELVDSEEEKKRASSFVRIGKSGLNDQDLFKRVSSFVRIGKSQGEEDKRVSSFVRIGKSGADEVEDEG--KRASSFVRIGKS----------DTPMD----------------------
KKASSFVRIGKSSTSPAETSSDSANSAISDEDPINIASRSSAFVRIGKIPSSAFVRIGKNTNLLTAPSENWKLGFRRGSRE-GQSSFVRIGK------------------------
>XP 009051543.1 hypothetical protein LOTGIDRAFT 228288 Lottia gigantea
--------------------------MECDESSVRRHQPIL-----WSKDSLLSVIDIGQHL-HMTTFDEMI------QLTRQIMEINLVFSLLLVCSISFVLSHPFTDDSNNDDQD----LKSAIQ-----------------------D----
GDTVPQAVKRPSSFVRIGRNPS------SFVRIGKA---------FGRFIRIGKND---------PNKRLSSFVRIGKSDPNK---RISSFVRIGK------SQEFNQ-EPEKRQSSFVRIGKSPE--LNENEIPNKRYSSFVRIGKSMDDGS-------

----LENPD----KRYSSFVRIGKNIENELTNAGLEKRPSSFVRIGKSYFAEPG----DMDAE----------------------KRLSNFVRIGKSGLEEPEM------------------EQKRAFVRIGKIPSSAFVRIGRMP--
LYDAILQKPLGYYNVARRMGKSSFVRIGKRNNEA------------------
>XP 011448017.3 FMRF-amide neuropeptides Crassostrea gigas
MLRPYHVIIVGLFYCYTTNAEINENKLLHPIKTEEGANEILGDKADDKRSRGFFRIGKKSAVENEANDKKFD------SKTVKE----------------------EDNYIPEKIQLIRVVNSESE------------TPIYVPVEFDPE--
SSDDTVDEDEKRASGFFRIGKSAENVDKRKGFFRIGKSVDQNPMNKKASGFFRIGRTP---------IDKRGKGFFRIGKSLNEMDEKRASGFFRIGKSALN-----------DKRSRGFFRI--GRSK-------------
GFFRIGKAFPLDGEKRASGFFRIGRNSPE-EMRKKASKFFRIGKSVNSKEEND---KRASGFFRIGKKCSGDSDKAGDNLTEDKSQSNPENEDTSESFNRNSDEPVRRASQFFRIGKSSSNKVTKRSSGVNSSPEQNLNLNKRAFFRI----
GKVPTSAFMRIGRQH--LLQSLVSDPLYRNGRIQQSS---FIRIGKRSMSDNHL---IDDEQSDSSL---
>KAI8790360.1 sodium-dependent neutral amino acid transporter B(0)AT3 Biomphalaria glabrata
-------------------------------------------------------------------------------------------------------------------------------------------------------
-----------------------------------------------MANGFNGVADMFLVASLLVSGCG-------------
KCAFLIPYAILLAIEGIPLFFLELAIGQRLRKGAIGAWNQVSPYLVGVGICSSIVSFYLSLYYNTIMSWCFVYLVQSFQSPLPWATCPHHFGENDTYI--LEE--
ECRKSSPPSYFWYRETLNTAPSIESTPVFNWKIALGLVIAWTVVFLCIIKGIKSTGKVVYVTATFPYFVLVIFFFRGITLEGFDKGLEHLFVPEWHRVLDPEV--
WLDAATQIFYSFGLGFGCLIALASYNPVRSDFLRDTFIVTICDFFTSIFTAIVVFSILGFKATLQFKDC--LKKHNVINQTED---SSHP------------------------
ACDFQKIISESASGTGLAFIAFTEAINQFPLAPLWAVLFFLMLLTLGLDSMFGTLEGAITSFNDMMLLPRIRKE--
FVCGAVCLICLLLSMCFATSTGPYVFALFDSFCANIPLLLVGFIECVAVSFVYGLKQISDDIELMLGRRPSYFWLICWRYVMPVAVIIIILLSSVIEIMVKGVYYEAWDADKAIAVMLPWPWWCKVLAAFLILSSVLCIPGLALLHHLGVSILPHETPAFF
PSEELKEEYGLNTHKINHFERVVLGFKE----------------------
>CAE1141792.1 SLC6A15S Sepia pharaonis
MFFRCQTKSTLSLCVHHLRLPVVPSPVT----------------------------------
GILVRGLLRRKQLSDSRKQRRHHTKFSSSFDTEETVEDTNFLIAAMKRSGSYSSALDEITNGSDKLSIDDRVSMGDRLSTSDRRSLDERMPARSYNSTPTGPSANGGPVYTIDMGKQHSSTNTPSIGRSTDDLISSSIGTVCNISFRDPDRESWDNKIQY
LLAAIGLAVGLGNVWRFPYLAQKNGGGAFLLPYIIMLFLEGLPIFYMELAMGQRMRNGPIGTWSQISPYLGGIGFSCVMVSYIVGIYYNTIIAWCLYYFGQSFRSKLLWSSCPTIPVPGARYKNETQPVKECELAGPTAYFWYRSTLDVSSSIEENGHML
WHIELCILFAWIIVFLCMIHGIASSGKVVYFTATFPYIVLVIFFFRGVTLKNFHVGLVHMFLPKFDRLADPQV--WLEAANQIFYSLGLAFGGLMAMSSYNPVNNNCLKDAYTVVFINCGTSIFAGIVIFSILGFKASVSYESC----LQTNIETGRNI-
----------------------------
TCNLQEELESTASGPGLAFIAFTEAINQFPAPPVWAILFFLMLITLGLDSMFGSLECVTTCIMDSGRFPDLKKKKFFVPAVLCLTSFLISFAFANQSGSYTFLLFDEFTGGFPLLVIALFEIISVSYIYGLNKFAEDVELMVGTRPNYYWLFMWKYISPL
VIIIMFASILKMMIQGLTYEIWDKETATKIDMPWPGWSLFIASMLIILILIWIPLVYIVKRFNLTGWKRDPPKHFPRDELRREKNLKKYILKDWEKKYLFIFEDLS-------------------
>NP 001284839.1 uncharacterized protein Dmel CG10804 isoform D Drosophila melanogaster
-------------------------------------------------------------------------------------MSDAFDES----------------------------SELEQDPPPFGSEASTSKAKAELVA--I-------
AT------ASADQQDIVEGE-----KE---G---------------------
EERESWDSKIMFLLATIGYAVGLGNVWRFPYLAQKNGGGAFLVPYFIMLCIQGIPIFYLELAIGQRLRKGAIGVWSQVSPYLGGIGISSAVVSYIVALYYNTIIAWCLIYLLHSFESPLPWADCPTRLYKNFTYD--HEP--
ECVASSPTQFYWYRTTLQCSESVDMPENFNYHMAIALIVSWFLVYICMVQGITSSGKIVYMTAIFPYVVLIIFFFRGITLKGAADGVAHLFTPRWETLLDPVV--
WLEAGTQIFFSLGLAFGGLIAFSSYNPANNNCYRDAILVSLTNCGTSMFAGVVVFSVIGFKATATFDRC--TEERNGLVAQNK--THNLP------------------------
VCDLQTELANSASGTGLAFIIFTEAINQFPGAQLWAVLFFLMLFTLGIDSQFGTLEGVVTSLVDMKLFPNLPKE--
YIVGALCFSCCTISMCFANGAGSYIFQLMDSFAGNFPLLIIALFECLSISYIYGVRRFSDDIEMMTGSRPNFYWMFCWKYLSPCAMVTILLASFYQLLTEGSSYPAWIGSKGATEGMEWPHWCIVVAFFLILSSILWIPIVAVLRLCGIKVVEDSDPAWF
PEAELREVHGIVPHEPTELERSIFCFNMDGTEGMCCPKYGLPEKSLEEEE-

SLC6a15/18
>ObockiSlC
-----------------------------------------------------------------------------------MERIEMNHLNIP----------------------------SPHSDENGDGN----SNTLYGSTMT--L-------
GASSPGVGASTANLLKIDLE-----KK---IDRHTG-----------------
EERETWDNRVQFILSLAGFAVGLGNVWRFPYLTQKNGGGAFLIPYAVMLFVEGIPIFYLELAIGQRLRKGAIGCWNQVSPYLGGLGIAAAVVSFNVALYYNTIMAWCMYYLVESFQSPLPWSSCPSETVGNITTF---NQ--
ECEKSGSTTYFWFRVALNSAPAIDSTPIISWKISAALICAWLLVFCCMIKGIKSSGKVVYVTATFPYFVLVIFFFRGVTLHGFEEGVKYLFIPKWDKLFEPAV--
WLDAATQIFYSFGLAFGCLVALASYNPVRSNFLSEALMVSVGDFFTSLLTATVVFSVIGFKATMAYEKC--LEKFGSTNHTKNPNITAEK------------------------
ECDFEKLMTESASGTGLAFIAFTEAIDQFPVAPLWAVLFFLMLLTLGLDTMFGTLEGAITSINDMLLFPAIRKE--
VVCGIVCFVSMLISFCFATSTGPYVFDLFDMYCANIPLLVIAFIEVIAVSYIYGLKQFSDDVEMMVGIRPSYFWMFIWRYLAPLVMIVIFVASLFEILENGSYYMAWDADKGHTVMQPRPWWAQFIAALLILSSVAWIPFVAITRYFNLVKWEPETPAFF
PAEELREERNIKPHQATFIEKYFFGFKG----------------------
>XP 005108791.1 sodium-dependent neutral amino acid transporter B(0)AT3-like Aplysia californica
MMDVGDDEKDIGRNINVAKSPST-----------------------------------------------------DVDERKLPIEMDRLMPP----------------------------NPADGTPGEADSMTGGSASYGSTLT------------
--IGADASETNLLRIDLE-----RK---FSRHTG-----------------
EERVTWDNRAQFVLSLIGYAVGLGNVWRFPYLTQKNGGGAFLIPYAVMLAIEGIPLFFLELAIGQRLRKGAIGAWNEVSPYLGGLGICSAIVSFYLALYYNTIMSWCLVYLFQSFQSPLPWAACPHTFGDNDSYV--IEP--
ECEKSSPPSYFWYRSTLNTAPTIESIPEFNWKIALGLLMAWVIVFLCILKGIQTTGKVVYVTATFPYLVLVAFFFRGITLEGFDRGLEHLFVPEWHRLLDPAV--
WLDAATQIFYSFGLAFGCLIALASYNPVRSDFLRDTLVVTVCDFLTSIFTAVVVFCVLGFKATMHYKDC--LSSHGLDNTTDPHTLDVNH------------------------
TCNFKKIIEESGSGTGLAFIAFTEAINQFPLAPIWAVLFFLMLMTLGLDSMFGTLEGAITSLNDLMLFPRARKE--
VVCGSVCLFCLVVSMSFATSTGPYVFALFDSFSANIPLLVVGFMECVAISYVYGLKQISEDIELMLGKRPSYFWLISWRYVMPLAVLVIFVSSLIEIFSEGIVYEAWDADKGIPVVLPWPWWCRLLAAALILSSILWIPGVALARRLGYVPLADETPAFF
PTDELREEHSITPHKDTQFERVVLGFRE----------------------
>XP 009059532.1 hypothetical protein LOTGIDRAFT 164761 Lottia gigantea
-------------------------------------------------------------------------------------------------------------------------------------------------------
MDIGPGASTANLLRIDLD-----KK---ISRHTGEGA-------------
EQRETWDNRAQFILSLVGYAVGLGNVWRFPYLTQKNGGGAFLIPYAVMLAVEGIPLFYLELAIGQRLRKGAYGAWNQVSPYLGGIGIASSIVSFNVALYYNTIMAWCLVYLVQSFQSPLPWASCPHDLGINDSYI--IVP--
ECEKSSPPSYFWYRDTLQTADSVESTPYLNWKIVVGLIGAWIIVYLCMIKGIKSSGKVVYVTATFPYIVLVVFFFRGVTLRGMEKGIEHLFVPEWNKLFDPAVCIWLS---------------
IALASYNPVRSNFLKETLLVTVCDFFTSIFTAVVVFSIL--------------------------------------------------------------ESASGTGLAFIAFTEAINQFPVAPLWAVLFFLMLFTLGLDS---------------
---------------VVCFFSLIVALCFATNTGPYVFSLFDSFSANIPLLIIAFMECVSISYVYGLQQ------------------------------------------------------------------------------------
-----------------------------------------------------------
>XP 011417303.2 sodium-dependent neutral amino acid transporter B(0)AT3-like Crassostrea gigas

----------------------------------------MYH-------------------GKARSRDHDPPR---RQMGRRNQYSYDNPALSESNSTPSIFTVESSSSLVHNNGFSPDLFPPIRRDSFESAVETGSRKSWRESTE--R-------
MSEDPAKVAGTSKDVLIGVE-----KE---GSSQES--L-------------
EERDSWDNKVQYLLAVVGYAVGLGNVWRFPYLAQKNGGGAFLIPYTIMLAVEGIPIFYLELAVGQRLRKGAIGAWNQISPFLGGIGIASAMVSFWVGLYYNTIISWCLYYLVYSFRSTLPWSECP----GNVT---------
ECTMSSPTTYFWYRETLNISPSVDERGSLNWWVVVSLLAAWIIVFLCMVKGIASSGKVVYVTATFPYLVLVIFFFRGVTLEGFEEGLKHFFIPDFTRLGDPQV--
WLEAATQIFFSLGVAFGGLIAFSSYMPVRNNCYKDAILVSVINCGTSVFAGVVIFSILGFKATQSFKAC--NAHNDLLWSQNK--TVGFL------------------------
TCDIKTEIEKSGSGTGLAFIAFTEAINQLPAAPVWSVLFFLMLLTLGLDSMFGMLEGVVTSIIDMNLIKNLRKD--
VVAAVLCLVSLLLSFCFADSAGPYIFVLFDEYSGNIPLLIIALGELLGLTYMYGLKRFSDDIELMTGQRPNYYWLIMWKYVSPIVIIIIFFASLIKSMMSAATYEAWNPSIADNITLEWPAWCKFIASLLIITSMGWIPLVAIVKKFNIIKWKPETPAEF
PEEELKAENEIKPYIPSDRERKWLYWLEKL---------------------
>XP 011424182.2 sodium-dependent neutral amino acid transporter B(0)AT3 Crassostrea gigas
-------------------------------------------------------------------------M---ERPKKKDKIEMDSLVESGENSVKN---------------------SPSGDDEKQNLQIPVSGQTYGSTLS------------
--IGPEASTANLLKIDLE-----KK---ISRHTG-----------------
EERESWDSRAEFLLSLIGYAVGLGNVWRFPYLTQKNGGGAFLIPYAVMLAIEGIPLFYLELAIGQRLRKGSVGSWNQVSPYLQGVGLASAAVSFNVALYYNTIMAWCIIYLVQSFQSPLPWSECPLVEGFNSTVK---EP--
ECERSGPTTYFWYRETLNIAPSIESSSGLNWKLAVSLLVAWIIVYLCMVKGIKSSGKVVYVTATFPYLVLIIFFFRGITLPGFERGLEHLFVPEWSKLFEPEV--
WLDAATQIFYSFGLAFGCLIALSSYNPIRNNFAKEALIVTVSDFFTSIFTACVVFSILGFKATMAYENC--LTQKENQTLGNS----TDT------------------------
VCDFKTIISESGSGTGLAFIAFTEAINQFPIAPLWSVLFFLMLFTLGLDSMFGTLEGALTSINDMMLFPKLRKE--
ILCGIVCLVSFLISLCFATRTGSYVFSLFDSFSANIPLLIIAFIELLGMSYIYGLQRLCDDIELMIGSRPGYFWLLCWRFIGPLAMVVIFVASLIEIFSKGVSYEVWNSLKGYTELAPWPWWCQVLAVLLILSSVLWIPGVALVKYYRLIDWKEEVPAYF
PEDELMEERQFQPHISNQFEKLLFGFKD-----------------------
>XP 014773791.1 sodium-dependent neutral amino acid transporter B(0)AT3 isoform X1 Octopus bimaculoides
----------------------------------------------------------------------MIA---RAHTNERELHVTTKMDTNKLPPKLQRSNSYLV-------------
SISDNEDGLSQISGSPRQLSISTIDGIGKKHQSTNGTCPSVASDTTNKSTEVLME------A---LRMEEEQNAPT-----------
KSRGAWDSKFQYMLAVIGYAVGLGNVWRFPYLVQKNGGGAFLIPYTLMLAIEGIPIFFLELALGQRMRKGAIGAWNQISPFMGGIGICCAIVSFIVGLYYNTIIAWCLYYLVLSFRSNLPWEKCPTVQVGKYKNESRLVE--
ECRRSGPTAYFWYRETLNISDSVDKSGVMSWWILVCLVFAWLLVFICIVKGIKSSGKVVYVTAIFPYIVLIIFFFRGVTLPGFQNGLYHLFVPEFSRLSDPQV--
WVDAASQIFYSLGLGFGSLIALSSYNPTRNNCHRDAIFIALTNCCTSVFAGIVVFSILGFKATKRLELCQEIRDHELMALFNDTNVIPPPAGTLVNISSSPNGEVTSVVMPNITTCSFKEEILKSASGTGLTFIAFTEAINQFSVPPIWSILFFLMLLTI
GLDSMFATLEGVVTPILDLDIIPDLRKYPPMLSACLCLSSFLLSFVFANQAGPYIFVLFDEYVSGLPLLIIVFAEIITISYIYGLYRFSNDIELMTGHRPNYYLMFCWKFICPAIIIFLIVILIINLTTDEKEYDVWHRETGSLSKSIWPGWCLFVAALI
IILSILGIPLIALIRWLKPSSWREEVPAYFPRELIQLERKLTTYIPKEWEKKILFRFEKHLPTTESEFTNKNKSQMDLVFI
>XP 036358020.1 sodium-dependent neutral amino acid transporter B(0)AT3-like isoform X2 Octopus sinensis
MAAEDSDKINLPENITIANEDITEIAETSFKEKSVKSKRKTPYY-------------------GKLVRGLLNRKR---QRRLHRRQLIYKKFMNSDDFRSKMDNE------------------KPDSNDTNSESFSMRRSASYSEALN--
DISFNVSEQSSSVFPNPNVSYELKISSEMRASAAT---LRQSTDELLPT-----------
DETRDSWDNKVQYLLAAIGLAVGLGNVWRFPYLAQKNGGGAFLLPYVIMLFVEGLPIFFLEVALGQRMRRGPVGTWDQISPYLTGIGYSCIMVSYIVCIYYNTIISWCLYYFALSFRSELLWAKCP-----NETAA---SK--
ECNLAGPSAYFWYRSTLDVSSSIEESGSLQLHLELCLFFAWLIVFLCMIKGIQSSGK----------------------------------------FEKLANPQV--
WLEAATQIFYSLSLGFGGLMAMSSYNPLRNNCYKDTFIVVFVNCGTSVFAGIVIFSILGFKAYKSFEDC----QSRLMEVTKNMTKEAPF------------------------
TCDLDKELENTAGGPGLAFIAFTEAINQFPVSPLWAILFFLMLLALGLDSMFGTLECITTLVTDSGKYPFLRQNRFFIPAVLCLTSFLISLSFAAQSGSYTFVLFDEFTGGLPLLFIAFAEVMSVSYIYGLSKFADDIHVMVGTRPSYFWLIMWKYISPL
VMMIILLASLIKMIITGMTYEVWDKDTATKIKTKWPGWCLFIGSSFVILILIWIPLIFIIKRFNIINWKPDTKVSFPKEELRAEKNLQPYEMKDWEKKYLFIFEDL---------------------
>XP 036360806.1 sodium-dependent neutral amino acid transporter B(0)AT3-like isoform X2 Octopus sinensis
-------------------------------------------------------------------------------MERIEMNHLNIP-----------------------------SPHSDENGDGNS----NTLYGSTMT--L-------
GASSPGVGASTANLLKIDLE-----KK---IDRHTG-----------------
EERETWDNRVQFILSLAGFAVGLGNVWRFPYLTQKNGGGAFLIPYAVMLFVEGIPIFYLELALGQRLRKGAIGCWNQVSPYLGGLGIAAAVVSFNVALYYNTIMAWCMYYLVESFQSPLPWSSCPSETVGNITTY---NQ--
ECEKSGSTTYFWFRVALNSAPSIDTAPIISWKISAALVCAWLLVFCCMIKGIKSSGKVVYVTATFPYFVLVIFFFRGVTLHGFEEGVKYLFIPKWDKLFEPAV--
WLDAATQIFYSFGLAFGCLVALASYNPVRSNFLSEALMVSIGDFFTSLLTATVVFSVLGFKATMAYEKC--LEKFGSTNHTKNPNTTVEK------------------------
ECDFEKLMTESASGTGLAFIAFTEAIDQFPVAPLWAVLFFLMLLTLGLDTMFGTLEGAITSINDMLLFPAIRKE--VVCGVVCFVSMLISFCFATSTGPYVFDLFDMYCANIPLLVIAFLEVIAVSYIYGLKQVTL------------------------
-------------------------------------------------------------------------------------------------------------------------------------
>XP 052823091.1 sodium- and chloride-dependent transporter XTRP3 Octopus bimaculoides
-------------------------------------------------------------------------------MERIEMNHLNIP-----------------------------SPHSDENGDGN----SNTLYGSTMT--L-------
GASSPGVGASTANLLKIDLE-----KK---IDRHTG-----------------
EERETWDNRVQFILSLAGFAVGLGNVWRFPYLTQKNGGGAFLIPYAVMLFVEGIPIFYLELAIGQRLRKGAIGCWNQVSPYLGGLGIAAAVVSFNVALYYNTIMAWCMYYLVESFQSPLPWSSCPSETVGNITTF---NQ--
ECEKSGSTTYFWFRVALNSAPAIDSTPIISWKISAALVCAWLLVFCCMIKGIKSSGKVVYVTATFPYFVLVIFFFRGVTLHGFEEGVKYLFIPKWDKLFEPAV--
WLDAATQIFYSFGLAFGCLVALASYNPVRSNFLSEALMVSVGDFFTSLLTATVVFSVLGFKATMAYEKC--LEKFGSTNHTKNPNITAEK------------------------
ECDFEKLMTESASGTGLAFIAFTEAIDQFPVAPLWAVLFFLMLLTLGLDTMFGTLEGAITSINDMLLFPAIRKE--
VVCGIVCFVSMLVSFCFATSTGPYVFDLFDMYCANIPLLVIAFIEVIAVSYIYGLKQFSDDVEMMVGIRPSYFWMFIWRYLAPLVMIVIFVASLFEILENGSYYMAWDAEKGHTVMQPRPWWAQFVAALLILSSVAWIPFIAIIRYFNLVKWEPETPAFF
PAEELREERNIKPHQATFIEKYFFGFKG-----------------------
>XP 055892413.1 sodium- and chloride-dependent transporter XTRP3-like Biomphalaria glabrata
MEQGGDSRIS-----------------------------------------------------SANGNLSPSS---DSLNRRFQIEMNCLAPPET-------------------------KSGEQSPAESEKMTPVASTYGSIMS------------
--IGPDASTVNLLKIDLE-----KK---INRHTG-----------------
EERVTWDNRAQFVLSLIGYAVGLGNVWRFPYLTQKNGGGAFLIPYAILLAIEGIPLFFLELAIGQRLRKGAIGAWNQVSPYLVGVGICSSIVSFYLSLYYNTIMSWCFVYLVQSFQSPLPWATCPHHFGENDTYI--LEE--
ECRKSSPPSYFWYRETLNTAPSIESTPVFNWKIALGLVIAWTVVFLCIIKGIKSTGKVVYVTATFPYFVLVIFFFRGITLEGFDKGLEHLFVPEWHRVLDPEV--
WLDAATQIFYSFGLGFGCLIALASYNPVRSDFLRDTFIVTICDFFTSIFTAIVVFSILGFKATLQFKDC--LKKHNVINQTED---SSHP------------------------
ACDFQKIISESASGTGLAFIAFTEAINQFPLAPLWAVLFFLMLLTLGLDSMFGTLEGAITSFNDMMLLPRIRKE--
FVCGAVCLICLLLSMCFATSTGPYVFALFDSFCANIPLLLVGFIECVAVSFVYGLKQISDDIELMLGRRPSYFWLICWRYVMPVAVIIIILLSSVIEIMVKGVYYEAWDADKAIAVMLPWPWWCKVLAAFLILSSVLCIPGLALLHHLGVSILPHETPAFF
PSEELKKEEYGLNTHKINHFERVVLGFKE-----------------------
>CAI9715349.1 sodium-dependent neutral amino acid transporter B0AT3-like Octopus vulgaris
MAAEDSDKINLPENITIANEDITEIAETSFKEKSVKSKRKTPYYADPILKLKLKRSSCANYYIGKLVRGLLNRKR---QRRLHRRQLIYKKFMNSDDFRSKMDNE------------------KPDSNDTNSESFSMRRSASYSEALN--
DISFNVSEQSSSVFPNPNVSYELKISSEMRASAAT---LRQSTDELLPT-----------
DETRDSWDNKVQYLLAAIGLAVGLGNVWRFPYLAQKNGGGAFLLPYVIMLFVEGLPIFFLEVALGQRMRRGPVGTWDQISPYLTGIGYSCIMVSYIVCIYYNTIISWCLYYFALSFRSELLWAKCP-----NETAA---SK--

ECNLAGPSAYFWYRSTLDVSSSIEESGSLQLHLELCLFFAWLIVFLCMIKGIQSSGKVIYFTATFPYVVLIIFFFRGVTLSNFYVGLKHLFIPKFEKLANPQV--
WLEAATQIFYSLSLGFGGLMAMSSYNPLRNNCYKDTFIVVFVNCGTSVFAGIVIFSILGFKAYKSFEDC----QSRLMEVTKNMTKEAPF------------------------
TCDLDKELENTAGGPGLAFIAFTEAINQFPVSPLWAILFFLMLLALGLDSMFGTLECITTLVTDSGKYPFLRQNRFFIPAVLCLTSFLISLSFAAQSGSYTFVLFDEFTGGLPLLFIAFAEVMSVSYIYGLSKFADDIHVMVGTRPSYFWLIMWKYISPL
VMMIILLASLIKMIITGMTYEVWDKDTATKIKTKWPGWCLFIGSSFVILILIWIPLIFIIKRFNIINWKPDTKVSFPKEELRAEKNLQPYEMKDWEKKYLFIFEDL---------------------
>XP 052825139.1 sodium-dependent neutral amino acid transporter B(0)AT3 isoform X1 Octopus bimaculoides
MAAEDSDKINLPGNIIIADENITAITETSFKEKPVKSKRKTPYY-------------------GKLVRGLLNRKR---QRRLHRRQLIYKKFMNFDDFQSKMDNE------------------KPDLNDTDSESFSMRRSTSYSEALN--I-
SVSDSVQSSNVFSDPRGPYELRISSEMYTSAVT---LRQSTDELIFT-----------
DETRDSWDNKVQYLLAAIGLAVGLGNVWRFPYLAQKNGGGAFLLPYVIMLFIEGLPIFYLEVALGQRMRRGPVGTWDQISPYLTGIGYSCIMVSYIVCIYYNTIISWCLYYFALSFRSELLWENCP-----NETAA---SK--
ECNLAGPTAYFWYRSTLDVSSSIEESGSMQLHLELCLFFAWLIVFLCMIKGIQSSGKVIYFTATFPYVVLIIFFFRGVTLSNFYVGLKHLFVPKFEKLANPQV--
WLEAATQIFYSLSLGFGGLMAMSSYNPLRNNCYKDTFIVVFVNCGTSVFAGIVIFSILGFKAYKSFEDC----QSRLMEVTKNMTKEAPF------------------------
TCDLDKELENTAGGPGLAFIAFTEAINQFPVSPLWAILFFLMLLALGLDSMFGTLECITTIITDSGKYPFLRQNRFFIPAVVCLTSFLISLSFAAQSGSYTFVLFDEFTGGLPLLFIAFAEVMSVSYIYGLSKFADDIHAMVGTRPGYFWLIMWKYISPL
VMLIIILFASLIKMIITSMTYEVWDKDTATKIKTKWPGWCLFIGSSFVILILIWIPLIFIIKRFNIINWKPDPKVSFPKDELKREKNLQPYVIKDWEKKYLFIFEDLYDS-----------------
>XP 029641283.2 sodium-dependent neutral amino acid transporter B(0)AT3-like isoform X1 Octopus sinensis
MAAEDSDKINLPENITIANEDITEIAETSFKEKSVKSKRKTPYY-------------------GKLVRGLLNRKR---QRRLHRRQLIYKKFMNSDDFRSKMDNE------------------KPDSNDTNSESFSMRRSASYSEALN--
DISFNVSEQSSSVFPNPNVSYELKISSEMRASAAT---LRQSTDELLPT-----------
DETRDSWDNKVQYLLAAIGLAVGLGNVWRFPYLAQKNGGGAFLLPYVIMLFVEGLPIFFLEVALGQRMRRGPVGTWDQISPYLTGIGYSCIMVSYIVCIYYNTIISWCLYYFALSFRSELLWAKCP-----NETAA---SK--
ECNLAGPSAYFWYRSTLDVSSSIEESGSLQLHLELCLFFAWLIVFLCMIKGIQSSGKVIYFTATFPYVVLIIFFFRGVTLSNFYVGLKHLFIPKFEKLANPQV--
WLEAATQIFYSLSLGFGGLMAMSSYNPLRNNCYKDTFIVVFVNCGTSVFAGIVIFSILGFKAYKSFEDC----QSRLMEVTKNMTKEAPF------------------------
TCDLDKELENTAGGPGLAFIAFTEAINQFPVSPLWAILFFLMLLALGLDSMFGTLECITTLVTDSGKYPFLRQNRFFIPAVLCLTSFLISLSFAAQSGSYTFVLFDEFTGGLPLLFIAFAEVMSVSYIYGLSKFADDIHVMVGTRPSYFWLIMWKYISPL
VMMIILLASLIKMIITGMTYEVWDKDTATKIKTKWPGWCLFIGSSFVILILIWIPLIFIIKRFNIINWKPDTKVSFPKEELRAEKNLQPYEMKDWEKKYLFIFEDL---------------------
>AAI14500.1 Tryptophan hydroxylase 2 Homo sapiens
------------------------------------------------------------------MQPAMMMFSSKYWARRGFSLD---SAVPEEHQLLGSSTLNKPNSGKNDDKGNKGSNKREA------ATESGKTAVVFSLKNEV--
GGLVKALRLFQEKRVNMVHIESRKSRR---RSSEVEIFVDCECGKTEFNELIQLLKFQTTIVTLNPPENIWTEEEE------LE---DVPWFPRKISELDKCSHRVLMYGSELDADHPGFKDNVYRQ--
RRKYFVDVAMGYKYGQPIPRVEYTEEETKTWGVVFRELSKLYPTHACREYLKNFPLLTKYCGYREDNVPQLEDVSMFLKERSGFTVRPVAGYLSPRDFLAGLAYRVFHCTQYIRHGSDPLYTPEPDTCHELLGHVPLLADPKFAQFSQEIGLASLGASDE
DVQKLATCYFFTIEFGLCKQEGQLRAYGAGLLSSIGELKHALSDKACVKAFDPKTTCLQECLITTFQEAYFVSESFEEAKEKMRDFAKSITRPFSVYFNPYTQSIEILKDTRSIENVVQDLRSDLNTVCDALNKMNQYLGI-------------------
----------
>AAM08923.1 neuronal tryptophan hydroxylase Mus musculus
------------------------------------------------------------------MQPAMMMFSSKYWARRGLSLD---SAVPEDHQLLGSLTQNKA--IKSEDKKSGKEPGKGD------TTESSKTAVVFSLKNEV--
GGLVKALRLFQEKHVNMLHIESRRSRR---RSSEVEIFVDCECGKTEFNELIQLLKFQTTIVTLNPPESIWTEED------LE---DVPWFPRKISELDRCSHRVLMYGTELDADHPGFKDNVYRQ--
RRKYFVDVAMGYKYGQPIPRVEYTEEETKTWGVVFRELSKLYPTHACREYLKNLPLLTKYCGYREDNVPQLEDVSMFLKERSGFTVRPVAGYLSPRDFLAGLAYRVFHCTQYVRHGSDPLYTPEPDTCHELLGHVPLLADPKFAQFSQEIGLASLGASDE
DVQKLATCYFFTIEFGLCKQEGQLRAYGAGLLSSIGELKHALSDKACVKSFDPKTTCLQECLITTFQDAYFVSDSFEEAKEKMRDFAKSITRPFSVYFNRYTQSIEILKDTRSIENVVQDLRSDLNTVCDALNKMNQYLGI-------------------
----------
>CAI9729217.1 tyrosine 3-monooxygenase-like Octopus vulgaris
-----------------------------------------------MNCSNNGDVENDTTKRRLAFQKCYSLEHGGNWRRKSLIEDARFETVTNAEFERSEERRL-GECISEETEQFNGEGKSRL------EVETARVSLLVILKEQM--
TSLADIMKVFESTKLQVMHIETRVSKQ---AGSKFDVFIQCMGKKVSINQCIELLDKNSKVHSTT---ILDESAKE------KV---REIWYPKHISDLDSCTHLLTKFEPELDSDHPGFTDKEYRR--
RRKLIADMAFTYRHNKSIPRVEYSEDEIKTWKHVYVNIKNLFPTHACSKHRHFFDLLEKKGIYSENFIPQLEDVSNFLKQQTGFQLRPVAGLLSARDFLSSLAFRVFQCTQYIRHGSKPDHTPEPDCVHELLGHVPMLADPEFAEFSQELGLASIGTSDE
DIEKFATLYWFTVEFGLVKEPCGIRAYGAATLSSYGELEHALSNNCEKKIFETGTASLQEYDDNDLQPIYFVAESFEDMKDKMRKYVAQIKRPYEFHYNAFTQSLVELKDKQSLQSVMRVIKGNVDCVQKALKHLDVHTVQKCIERSGEINL--------
----------
>NP 775489.2 tryptophan 5-hydroxylase 2 Homo sapiens
------------------------------------------------------------------MQPAMMMFSSKYWARRGFSLD---SAVPEEHQLLGSSTLNKPNSGKNDDKGNKGSSKREA------ATESGKTAVVFSLKNEV--
GGLVKALRLFQEKRVNMVHIESRKSRR---RSSEVEIFVDCECGKTEFNELIQLLKFQTTIVTLNPPENIWTEEEE------LE---DVPWFPRKISELDKCSHRVLMYGSELDADHPGFKDNVYRQ--
RRKYFVDVAMGYKYGQPIPRVEYTEEETKTWGVVFRELSKLYPTHACREYLKNFPLLTKYCGYREDNVPQLEDVSMFLKERSGFTVRPVAGYLSPRDFLAGLAYRVFHCTQYIRHGSDPLYTPEPDTCHELLGHVPLLADPKFAQFSQEIGLASLGASDE
DVQKLATCYFFTIEFGLCKQEGQLRAYGAGLLSSIGELKHALSDKACVKAFDPKTTCLQECLITTFQEAYFVSESFEEAKEKMRDFAKSITRPFSVYFNPYTQSIEILKDTRSIENVVQDLRSDLNTVCDALNKMNQYLGI-------------------
----------
>EAX02494.1 tyrosine hydroxylase isoform CRA e partial Homo sapiens
--------------------------------------------------------------------------------------DLHTEPCPPPTPPRHRPRASAGPCLSWTPSRQRPSWATKP-------------------------
SALSRAVKVFETFEAKIHHLETRPAQRPRAGGPHLEYFVRLEVRRGDLAALLSGVRQVSEDVR--------SPAGP-----------KVPWFPRKVSELDKCHHLVTKFDPDLDLDHPGFSDQVYRQ--
RRKLIAEIAFQYRHGDPIPRVEYTAEEIATWKEVYTTLKGLYATHACGEHLEAFALLERFSGYREDNIPQLEDVSRFLKERTGFQLRPVAGLLSARDFLASLAFRVFQCTQYIRHASSPMHSPEPDCCHELLGHVPMLADRTFAQFSQDIGLASLGASDE
EIEKLSTLYWFTVEFGLCKQNGEVKAYGAGLLSSYGELLHCLSEEPEIRAFDPEAAAVQPYQDQTYQSVYFVSESFSDAKDKLRSYASRIQRPFSVKFDPYTLAIDVLDSPQAVRRSLEGVQDELDTLAHALSAIG------------------------
----------
>KAI8791548.1 tyrosine 3-monooxygenase partial Biomphalaria glabrata
----------------------------------------------------------------------------------------------------------------------------------------------------------------
--NGKVFITHVESRKSLA---SVHQYQLFLQVVCSHDVFLNVCAEAKQSPLITELK----LLEEKEP------EK---KDVWFPKHISDLDRCTHLITRFEPDLDYTHPGFADKNYRL--
RRKEIADIAFGYKYGQSIPRVQYTEEENNTWAHVYRNLKNLFPTHACQEHIEVFNLLEKEGGFCEEKIPQLEDVSEFIKRKTGFQLRPVAGLLSARDFLASLAFRTFQCTQYVRHGAKPDHSPEPDCIHELLGHVPMLADPKFAQFAQELGLATLGVSDE
EIEKFATLFWFTVEFGLCRQNGEIRAYGAGMLSSYGELENCLSGAPVVKEFDPAVTAVQEYKDDDFQPILFVVDSFDEMMFKMRLVR-------------------------------------------------------------------------
----------
>NP 001191619.1 tryptophan 5-monoxygenase Aplysia californica
---------------------------------------------MGNCMEMYKVTLRNTRGIGGPPPQEDRKPFPICNHLPERRSMGLREGLARKRLCGDIRPPSESQLKALEESLRNLDGMNSP------SGLGTTTTVVFSVNNHV--
GHLASALKIFQENKINVVHIESRKSRR---ADSEYDIYVDVETDHIRLEELVNRLKREVASITFNQLNIPMTPPALTKAAC-ME---NVPWFPRMVYDLDQCAHNVLMYGTELNAEHPGFKDTVYRE--
RRKVFADLAMNYKHGLEIPYIQYTPTEVETWGTVFRELMKLYPTHACREYLANIPLLVEHCGYREDNVPQLEDISRFLKARTGFTLRPVAGYLSSRDFLAGLAFRVFHCTQYIRHDTDPFYTPEPDCCHELMGHVPLLADPSFAQFSQEIGLASLGATDN
DVAKLSTCYFFSVEFGLCKQDGELRAYGAGLLSSISELGHALSDKADKKVFEPLHMCKQECIITTFQDVYFYTDSFEEAKEKMRQFASTIKRPFAVRYNPYTESVDVLDSTRCIATVVSELKGDLCIVSDALKRLQIMERFKFQD--------------
----------
>NP 571224.1 tyrosine 3-monooxygenase Danio rerio

```
-------------------------------------------------------------------
MPNSSSSTSSTKSIRRAASELERSDSITSQKFVGRRQSLIEDARKEREAAAAAAEAAGLSEQIVFEESDGKALISLFFTLRCSKSPALSRTLKVFETFEAKIHHLETRPSRKPKDGLEDLEYYVQCEVHLSDVSTLVSSLKRSAEDVKTT----------
K------EV---KFHWFPRKIAELDKCHHLVTKYDPDLDQDHPGFTDPVYRK--
RRKMIGDIAFKYRHGEPIPRVDYTEEEIGTWREVYSTLRDLYTTHACSEHLEAFRLLEKHCGYSPDNIPQLEDVSCFLKERTGFQLRPVAGLLSARDFLASLAFRVFQCTQYIRHASSPMHSPEPDCVHELLGHVPILSDRTFAQFSQSIGLASLGASDE
DIEKLSTMYWFTVEFGLCKQGGVIKAYGAGLLSSYGELVHSLSDEPERREFDPDIVAVQPYQDQTYQPVYFVSESFVDATEKLRTYVTRIKRPFSVRFDPYTDSIEVLDNPLKIQKGLETIKDELKILTDALNVLA--------------------
----------
>NP 775567.2 tryptophan 5-hydroxylase 2 Mus musculus
-------------------------------------------------------------------MQPAMMMFSSKYWARRGLSLD---SAVPEDHQLLGSLTQNKA--IKSEDKKSGKEPGKGD------TTESSKTAVVFSLKNEV--
GGLVKALRLFQEKHVNMLHIESRRSRR---RSSEVEIFVDCECGKTEFNELIQLLKFQTTIVTLNPPESIWTEEED------LE---DVPWFPRKISELDRCSHRVLMYGTELDADHPGFKDNVYRQ--
RRKYFVDVAMGYKYGQPIPRVEYTEEETKTWGVVFRELSKLYPTHACREYLKNLPLLTKYCGYREDNVPQLEDVSMFLKERSGFTVRPVAGYLSPRDFLAGLAYRVFHCTQYVRHGSDPLYTPEPDTCHELLGHVPLLADPKFAQFSQEIGLASLGASDE
DVQKLATCYFFTIEFGLCKQEGQLRAYGAGLLSSIGELKHALSDKACVKSFDPKTTCLQECLITTFQDAYFVSDSFEEAKEKMRDFAKSITRPFSVYFNPYTQSIEILKDTRSIENVVQDLRSDLNTVCDALNKMNQYLGI-------------------
----------
>NP 840091.2 tryptophan 5-hydroxylase 1a Danio rerio
------------------------------------------------------------------------MYSSKSDGPRRGRSFDSMNLGMTLEEKQLNNEMNKSAFTKIEENKDNKTE-----------SSETGRAAVVFSLKNEV--
GGLVKALKLFQENHVNLVHIESRKSKR---RNSEFEIFVDCDSNREQLHEIIQLLRKHVNVVEMDAPDNRLPEESE------ME---NVPWFPKKISDLDKCANRVLMYGSDLDADHPGFKDNVYRK--
RRKYFADLAMSYKHGDPIPRIEFTEEEVKTWGVVFRELNKLYPSHACREYLKNLPLLIKHCDSREDNIPQLEDVSRFLKERTGFTIRPVAGYLSPRDFLAGLAFRVFHCTQYVRHSSDPLYTPEPDTCHELLGHVPLLAEPSFAQFSQEIGLASLGASDD
SIQKLATCYFFTVEFGLCKQEGKLRAYGAGLLSSISELKHALSGNARILPFDPNVTCKQECIITTFQDVYFMSDSFEEAKVKMREFAKTIKRPFSVRYNPYTQSVDVLKDTTSINNVVEELRHELDIIGDALSRLNKQLGV-------------------
----------



Tryptophan Hydroxylase and Tyrosine Hydroxylase
>ObockiTpH
------------------------------------------------------------------------------MDARSISFE-------RAEVMRRQQSTDFDDHIAVVPEETENGRGCSSR------EKVG---XILLSLKNQV--
GCLSSAXKVFQDNKVNVTHIESRRSQM---EDSEYDIFINVEXSKLAIEDLLEKLQLQIPDDVGNVKAFPTCPEEP------DD---DIVWFPRKIAELDKFANRVLMYGRELDADHPGFKDPVYRQ--
RRIKFAEIAYNYKHGQPIPRVEYTAEEVNTWGTVFRELNKLYPTYSCKQHLKNLPLLMKYCGYRENNIPQLEDVSKFLKERTGFTVRPVAGYLSSRDFLAGLAFRVFHCTQYIRHSSDPFYTPEPDCCHELLGHVPLFADPDFAHFSHELGLASLGASDF
EVAKLASCYFFTIEFGLCRENGQLRVYGAGLLSSIGELKHVLTDAAIKKPFDPDKVIEQECQVSTYQDAYFVSESFKEAQNQIREFASNIKRPFTVRYDPYSQSVMVLESLSTISQVVSFVRADLCLISDAINKLHQSKMKKTRRARTRLLNDK------
----------
>ObockiTyH
--------------------------------------------------------------------------------------------------------------------------------------------------------
MTSLADIMKAFESTKLQVMHIETRVSKQ---AGSKFDVFIQCVGKKVSINQCIELLDKNSKVHSTT---ILDESAKE------KV---REIWYPKHISDLDSCTHLLTKFEPELDSDHXGFTDKEYRR--
RRKLIXDMAFTYRHNMNIPRVEYXEDEIXTWKHVYVNIKNLFPTHACSKHRHFFDLLEKKGIYSENFIPQLEDVSNFLKQQTGFQLRPVAGLLSARDFLSSLAFRVFQCTQYIRHGSKPDHTPEPDCVHELLGHVPMLADPEFAEFSQELGLASIGTSDE
DIEKFATLYWFTVEFGLVREPCGIRAYGAATLSSYGELEHALSNNCEKKPFETETASLQEYDDNDLQPIYFVTESFEDMKDKMRKYVAQIKRPCEFHYNAFTQSLVELKDKQSLQSMMRIIKGDVDSVQKALKYLDVRTVQVTA----------------
----------
>XP 009060391.1 hypothetical protein LOTGIDRAFT 106041 Lottia gigantea
-------------------------------------------------------------------------------------------------------------------------------------------------------------------
----------------------MYQFLIFFYFVPDT---------------------------------------------WFPTHISELDKCTHLVTKFEPELDSDHPGFTDKAYRA--
RRQKIADIAFDYRHGDPIPRVDYTGEDIATWGHVYRQLKVLFPTHACKTHRDIFQLLEDECGYSPDRIPQLEDVSNFLKRKTGFQLRPVSGLLSARDFLASLAFRVFQCTQYVRHGSKPDHSPEPDCVHELLGHVPMLADPGFAQFSHELGLNSLGASDE
DIEKFATLYWFTVEFGICKQAGQLRAYGAGILSSYGELLHALGDKPERRPFDPLKTAVQEYTDEDLQPLYFVVESFEDMMDKMRNYSATIKRPYEVRYCSYTQTVQKLNSKEMLYSVARDLKGELEYLERAIDRIQCLTMTGR-----------------
----------
>XP 009062206.1 hypothetical protein LOTGIDRAFT 229367 Lottia gigantea
----------------------------------------------------------------------------------------------------------MVVYNKIHEDSLRNIDGISSP------SGFGKTTSIVFSLRNQI--
GHLASALQIFQENNINVVHIESRKSRM---KDAQYDIYVDVETDNIRLEELINRLKKEVANITFNDITVPLSPPPNVEDPCDMS---NVPWFPRNIEDLDFSAHRVLMYGTELDADHPGFKDDIYRA--
RRNYFSRIAMQYKHGEPIPHIEYSGEEIKTWGRVFRELNSLYPSHACREYLRNIPLLVEHCGYREDNVPQLEDVSNFLKARTGFQLRPVAGYLSSRDFLAGLAFRLFHCTQYIRHGSDPFYTPEPDCCHELMGHMPLLADASFAQFSQEIGLASLGASDE
EVSKLATCYFFTVEFGLCKQDGELRAYGAGLLSSIGELKHALTDNAKKIRFEPMRTSQQECLITTFQDVYFYTDSFEDAKERMRQFASTIKRPFAVRYNPYTNSVDVLDNTQCIATVVSELKGDLCIVSDALRRLQMLEKQLNECSPANQLDSPAASGYV
----------
>XP 011438860.1 tryptophan 5-hydroxylase 1 Crassostrea gigas
--------------------------------------------------------------------MRNSAICGPDLDNPEEKRDMGKAEQFRQKRRCKSIRAPTEAEIASLEENLKQLDGIFSP------SGIGKTTSVVFTLQNRV--
GNLANALKIFQDNHINVVHIESRKSRS---SDTEAEIYVELETDNIRLQELIKRLKKQVAGISYNDIPVPPSPATAARVPATAEALANAPWFPQKMADLDKAANRVLMYGSELDADHPGFKDKVYRQ--
RRTIFADIAMAYKNGQPIPRVDYTREEINTWGVVFRELNKLYPSHACREYLRNLPLLIDHCGYRDDNIPQLQDVSDFLKERTGFQLRPVAGYLSSRDFLAGLAFRVFHCTQYIRHSSDPFYTPEPDCCHELLGHMPLLADPSFAQFSQELGLASLGASDE
EVQKLATCYFFTVEFGLCKQDGHLRAYGAGLLSSIGELSHALTDKARKIPFEPTRTCKQECLITTFQDVYFFTESFEEAKERMRQFACTIKRPFAVRYNPYTQTVDVLNNTRCIAYAVSDLRGDLCIVSDALRRLQQLEFQKEDEEREMDPSTPR-----
----------
>XP 011440999.1 tyrosine 3-monooxygenase Crassostrea gigas
--------------------------------------------MFATEASVISPTQDPATVKRRLAFQKSYSQEHGGSWRRKSLIEDARFETVTNIEFAKRERTISNRGSISEEEEVFTQNGDTPSPV----
ETDPSRLYVILVTLKEGMTASLSRIVRVFETTKVFLDHIESRKSHK---AGAIFDVLIQFVCSRQKVTSLMNAIRQNPSVEEVT----VIGDKDA------DI---QDAWFPRHISELDNCTHLVTKFEPELDCDHPGFTDEEYKK--
RRKQIADIAFEYKHGHPIPRVQYTREEIATWGHVYKQLKELFPTHACREHIEAFKGLEKDCGYRETNIPQLEDVSNFLKRKTGFQLRPVSGLLSARDFLASLAFRVFQCTQYVRHGSKPDHSPEPDCVHELLGHVPMLSVPAFAQFSQEIGLASLGASDQ
EIEKLATLYWFTVEFGLVRQDGKLRAYGAGTLSSYGELKHALSDSPKQLPFSPTVTSVQEYTDEDLQPIYFYVESFEDMMQKMRDHVSTIKRSVDLRYDPITQSIQVIEHKDTLETLASSLRTEVRNLEKLVKRMDLVFV--------------------
----------
>XP 013086300.2 tryptophan 5-hydroxylase 1-like Biomphalaria glabrata
----------------------------------------------MGNCLEIYKVTLRNTPGIYGPPSQEEKKAFPMMNHHQDRKPMGLREGLARKRLCDDIRPPSESQLKALEESLRNLEGMNSP------SGLGTTTTVVFSVNNHV--
GHLASALNIFQENNINVVHIESRKSRR---SDALFDIYVDFETDHIRLEELVNRLKREVASLTFNQLNIPMTPPALTKKAC-LE---LVPWFPKKVSELDQAAINVLMYGSELDADHPGFKDTVYRQ--
RRKFYTDLAMQYTHGTDIPVIEYTKEEVETWGTVFRELMKLYPTHACREYLANIPLLVEHCGYREDNVPQLADVSRFLKERTGFTLRPVAGYLSSRDFLAGLAFRVFHCTQYIRHRSDPFYTPEPDCCHELMGHMPLLADPSFAQFSQEIGLASLGASDN
```

DIAKLSTCYFFSVEFGLCKQDNELRAYGAGLLSSISELAHALSDKAVKKVFEPLHMCTQECLITTFQDVYFYTDSFEEAKEKMRQFASTIKRPFAVRYNPYTESVDVLDSTRCIASVVSELKGDLCIVSDALKRLQLIENFRFQESHKQGSPQSLTDICK
EEVNSLQIKN
>XP 014778918.1 tryptophan 5-hydroxylase 1 Octopus bimaculoides
----------------------------------------------------------------------------------------------------------------------------------------------------------------
----------------------------------------MVLQRMKG-----------------------------------PRILSEV-IPHHTVTLCRSCMFSE--GFKDPVYRQ--
RRIKFAEIANNYKHGQPIPRVEYTAEEVNTWSTVFRELNKLYPTYSCKQHLKNLPLLMKYCGYRENNIPQLEDVSKFLKERTGFTVRPVAGYLSSRDFLAGLAFRVFHCTQYIRHSSDPFYTPEPDCCHELLGHVPLFADSDFAHFSHELGLASLGASDF
EVAKLASCYFFTIEFGLCRENGQLRVYGAGLLSSIGELKHVLTDAAIKKPFDPDKVIEQECQVSTYQDAYFVSESFKEAQNQIREFASNIKRPFTVRYDPYSQSVMVLESLSTISQVVNFVRADLCLISDAINKLHQSKMKKTRRTRTRLLNDK------
----------
>XP 017209547.1 tryptophan 5-hydroxylase 1 isoform X1 Danio rerio
------------------------------------------------------------------------------MLSGILLSGRRGQGRLLSHMRDRRNMMNKSAFTKIEENKDNKTE----------SSETGRAAVVFSLKNEV--
GGLVKALKLFQENHVNLVHIESRKSKR---RNSEFEIFVDCDSNREQLHEIIQLLRKHVNVVEMDAPDNRLPEESE------ME---NVPWFPKKISDLDKCANRVLMYGSDLDADHPGFKDNVYRK--
RRKYFADLAMSYKHGDPIPRIEFTEEEVKTWGVVFRELNKLYPSHACREYLKNLPLLIKHCDSREDNIPQLEDVSRFLKERTGFTIRPVAGYLSPRDFLAGLAFRVFHCTQYVRHSSDPLYTPEPDTCHELLGHVPLLAEPSFAQFSQEIGLASLGASDD
SIQKLATCYFFTVEFGLCKQEGKLRAYGAGLLSSISELKHALSGNARILPFDPNVTCKQECIITTFQDVYFMSDSFEEAKVKMREFAKTIKRPFSVRYNPYTQSVDVLKDTTSINNVVEELRHELDIIGDALSRLNKQLGV-------------------
----------
>XP 036361567.1 tryptophan 5-hydroxylase 1 Octopus sinensis
MSGQSKRYVWTVQKGDGRKSRELDSMQRSSDSKPNFSKIPMCHKIYRNCGERLSFSVRVYIFEVCPLLKKAFHKHGGNWRRKSLIEDARFETVTNAEFERSEERRL-GECISEETEQFNGEGKSRL------EVETARVSLLVILKEQM--
TSLADIMKVFESTKLQVMHIETRVSKQ---AGSKFDVFIQCMGKKVSINQCIELLDKNSKVHSTT---ILDESAKE------KV---REIWYPKHISDLDSCTHLLTKFEPELDSDHPGFTDKEYRR--
RRKLIADMAFTYRHNMSIPRVEYSEDEIKTWKHVYVNIKNLFPTHACSKHRHFFDLLEKKGIYSENFIPQLEDVSNFLKQQTGFQLRPVAGLLSARDFLSSLAFRVFQCTQYIRHGSKPDHTPEPDCVHELLGHVPMLADPEFAEFSQELGLASIGTSDE
DIEKFATLYWFTVEFGLVKEPCGIRAYGAATLSSYGELEHALSNNCEKKIFETGTASLQEYDDNDLQPIYFVAESFEDMKDKMRKYVAQIKRPYEFHYNAFTQSLVELKDKQSLQSVMRVIKGNVDCVQKALKHLDVRTVQVTA----------------
----------
>XP 036361997.1 tryptophan 5-hydroxylase 1-like Octopus sinensis
-----------------------------------------------------------------------------MDARSISFE-------RAEVMRRQQSTDFDDHIAVVPEETENGRGCSSR------EKVG---SILLSLKNQV--
GCLSSALKVFQDNKVNVTHIESRRSQM---EDSEYDIFINVESSKLAIEDLLEKLQLQIPDDVGNVKAFPTCPEEP------DD---DIVWFPRKIAELDKFANRVLMYGRELDADHPGFKDPVYRQ--
RRIKFAEIAYNYKHGQPIPRVEYTAEEVSTWSTVFRELNKLYPTYSCKQHLKNLPLLMKYCGYRENNIPQLEDVSKFLKERTGFTVRPVAGYLSSRDFLAGLAFRVFHCTQYIRHSSDPFYTPEPDCCHELLGHVPLFADSDFAHFSHELGLASLGASDF
EVAKLASCYFFTIEFGLCRENGQLRVYGAGLLSSIGELKHVLTDAAIKKPFDPDKVIEQECQVSTYQDAYFVSESFKEAQNQIREFASNIKRPFTVRYDPYSQSVMVLESLSAISQVVSFVRADLCLISDAINKLHQSKMKKTRRARTRLLNDK------
----------
>XP 052827174.1 tyrosine 3-monooxygenase Octopus bimaculoides
----------------------MELFSKMFIKCKDIIAQFPEEQRPQLTGKGTPPPRRGKGTPPPPPRVRPPRPHGGNWRRKSLIEDARFETVTNAEFERSEERRL-GECISEESEQFNGERKSRL------EVETARVSLLVILKEQM--
TSLADIMKVFESTKLQVMHIETRVSKQ---AGSKFDVFIQCVGKKVNINQCIELLDKNSKVHSTT---ILDESAKE------KV---
REIYYPKHISDLDSCTHLLTKFEPELDSDHPGFTDKEYRRVANNHDMSNFHFLICSNMSIPRIDYTEDEIKTWKHVYVNIKNLFPTHACSKHRHFFDLLEKKGIYSENFIPQLEDVSNFLKQQTGFQLRPVAGLLSARDFLSSLAFRVFQCTQYIRHGSK
PDHTPEPDCVHELLGHVPMLADPEFAEFSQELGLASIGTSDEDIEKFATLYWFTVEFGLVKEPCGIRAYGAATLSSYGELEHALSNNCEKKLFQTETASLQEYDDNDLQPIYFVAESFEDMKDKMRKYVAQINRPCEFHYNAFTQSLVELKDKQSLQSVM
RIIKGNVDSVQKALKYLDVRTVQVTA--------------------------
>XP 786206.2 tyrosine 3-monooxygenase Strongylocentrotus purpuratus
-----------------------------MAEDSPKTAKNGDFSLPELAPVKPPPRFTAGKPKRTPFEKLSKSFDDMSFVYPTRKKSLIEDARRESTRSSLDITSMSLVLSSPEEKDDVFDDAFEPQQ----
ESAIRRFTVTFSSKEDMGFGSLSEALRVFQKRKVTLTHVESRPSNK---IDGQIEFLMQCETKGSSSKNVLTALQKVADNVRLE-------KEEI------TK---RGPWFPTRVHELDRCTHLLSNYEPDLDDEHPGFTDKDYRE--
RRQRIADVAFKYKHGQPIPRVEYTDDELRTWGLIYRQLKALFPTHACKEHIDAFNILEKEGLYSESFIPQHEDVSNFLKGKTGFQLRPVAGLLSARDFLASLAFRVFQATQYVRHSSAPMHTPEPDCCHELLGHVPMLADPTFAQFSQEIGLASLGVADE
DITRLATLYWFTVEFGLCRQNGETRACGAGLLSAFGELQYALSDKPEHRPFEPNKTAIQEYQDKNYQPIYFVADSFSDAQSKLRLYAMKMARPYNVRYDPYTQSIQVIDKVDKLRDAIRDLNGQMVVLTSAIEKLM------------------------
----------

Vesicular GABA Transporter
>Obocki vGAT
------MT----AKSIYYIEKLKSFIRRSS-----------TDETVHFVKYSQFE-NEKE-----LTEIFDDKKK---FGSITDESNGSA---------------SISDV------
KDSKQTLANPHQNPVMDAVDTKLERNRITSWQAGWNVTNAIQGMFIVSFPYTVVQGGYWALVSMIVVAYICCHTGKILVECLYEVDS-NGYLVRVRHSYVEIAEEAMGKRF---
GGKIVNVAQLIELLMTCILYVLLSGELILGSFPESPISLTFWIMIATTPLLSYAFXKSLRGVSWLSFWCTMAHMAINAFILIFCFTRATEWQWKAVQIKVDIWTFPISIGIVVFSYTSQIFLPTLEENMEDRSRFHCMMHWTHIAAAFFKVLFSYICFLT
WGNGTQEVITNNLPTQSLKIIVNLILVAKSTLSYPLPYYAAVEMIQTTYFNGKPSTYFPSCFDENCRLKIWAGCLRIGLVLATMAMAIFIPHFAILMGLIGSXTGTMLSFVWPCFFHIKIKGKILPWYTKVLDISIIIIGVVVGSIGIYYSFRAL-----
-------------------------------------------------------------------------------SLAFRGIPTTPFHARDFHKSL------------------------------------------------------------
-----------------------------------------
>BAC44888.1 vesicular GABA transporter a form Mus musculus
MATLLRSK----LTNVATSVSNKSQAKVSGMFARMGFQAATDEEAVGFAHCDDLDFEHRQ-----GLQMDILKSE---GEPCGDEGAEAP---------------VEGDIHYQRG-
GAPLPPSGSKDQAVGAGGEFGGHDKPKITAWEAGWNVTNAIQGMFVLGLPYAILHGGYLGLFLIIFAAVVCCYTGKILIACLYEENE-
DGEVVRVRDSYVAIANACCAPRFPTLGGRVVNVAQIIELVMTCILYVVVSGNLMYNSFPGLPVSQKSWSIIATAVLLPCAFLKNLKAVSKFSLLCTLAHFVINILVIAYCLSRARDWAWEKVKFYIDVKKFPISIGIIVFSYTSQIFLPSLEGNMQQPSE
FHCMMNWTHIAACVLKGLFALVAYLTWADETKEVITDNLPG-
SIRAVVNLFLVAKALLSYPLPFFAAVEVLEKSLFQEGSRAFFPACYGGDGRLKSWGLTLRCALVVFTLLMAIYVPHFALLMGLTGSLTGAGLCFLLPSLFHLRLLWRKLLWHQVFFDVAIFVIGGICSVSGFVHSLEG----------------------
-----------------------------------------------------------------KFAGLET----------------------------------------------------------------------------------------
-------------------------
>NP 001074170.1 vesicular inhibitory amino acid transporter Danio rerio
MATLIRSKISNKLSNAATTVTNKSQAKVSGMFARLGFQAATDEEALGFAACDDLDYDHRQ-----GLQMDILKNDEMGGEGGGEMGEDGM---------------AEGDSHYQRD-GTGPPPSASKDG--
GLCSEIGNPDKPRITAWEAGWNVTNAIQGMFVLGLPYAILHGGYLGLFLIIFAAVVCCYTGKILIACLYEENE-
DGQLVRVRDSYVDIANACCAPRFPALGGHVVNVAQIIELVMTCILYVVVSGNLMYNSFPTLPVSQRSWAIIATAALLPCAFLKNLKAVSKFSLLCTLAHFVINILVIAYCLSRARDWAWDKVKFYIDVKKFPISIGIIVFSYTSQIFLPSLEGNMQKPSE
FHCMMNWTHIAACILKGLFALVAYLTWADETKEVITDNLPS-
SIRAVVNLFLVSKALLSYPLPFFAAVEVLEKTFFQDGGRAFFPDCYGGDGRLKSWGLSLRCALVVFTMLMAIYVPHFALLMGLTGSLTGAGLCFLLPSLFHLKLLWRKLLWHQVFFDVAIFVIGGICSISGFIHSVEGL---------------------
---------------------------------------------------------------IEAYKYNLPD---------------------------------------------------------------------------------------
-------------------------

>NP 001408116.1 vesicular inhibitory amino acid transporter Mus musculus
MATLLRSK----LTNVATSVSNKSQAKVSGMFARMGFQAATDEEAVGFAHCDDLDFEHRQ-----GLQMDILKSE---GEPCGDEGAEAP---------------VEGDIHYQRG-
GAPLPPSGSKDQAVGAGGEFGGHDKPKITAWEAGWNVTNAIQGMFVLGLPYAILHGGYLGLFLIIFAAVVCCYTGKILIACLYEENE-
DGEVVRVRDSYVAIANACCAPRFPTLGGRVVNVAQIIELVMTCILYVVVSGNLMYNSFPGLPVSQKSWSIIATAVLLPCAFLKNLKAVSKFSLLCTLAHFVINILVIAYCLSRARDWAWEKVKFYIDVKKFPISIGIIVFSYTSQIFLPSLEGNMQQPSE
FHCMMNWTHIAACVLKGLFALVAYLTWADETKEVITDNLPG-
SIRAVVNLFLVAKALLSYPLPFFAAVEVLEKSLFQEGSRAFFPACYGGDGRLKSWGLTLRCALVVFTLLMAIYVPHFALLMGLTGSLTGAGLCFLLPSLFHLRLLWRKLLWHQVFFDVAIFVIGGICSVSGFVHSLEGL---------------------
---------------------------------------------------------------IEAYRTNAED---------------------------------------------------------------------------------------
-------------------------

>NP 542119.1 vesicular inhibitory amino acid transporter Homo sapiens
MATLLRSK----LSNVATSVSNKSQAKMSGMFARMGFQAATDEEAVGFAHCDDLDFEHRQ-----GLQMDILKAE---GEPCGDEGAEAP---------------VEGDIHYQRGSGAPLPPSGSKDQ-
VGGGGEFGGHDKPKITAWEAGWNVTNAIQGMFVLGLPYAILHGGYLGLFLIIFAAVVCCYTGKILIACLYEENE-
DGEVVRVRDSYVAIANACCAPRFPTLGGRVVNVAQIIELVMTCILYVVVSGNLMYNSFPGLPVSQKSWSIIATAVLLPCAFLKNLKAVSKFSLLCTLAHFVINILVIAYCLSRARDWAWEKVKFYIDVKKFPISIGIIVFSYTSQIFLPSLEGNMQQPSE
FHCMMNWTHIAACVLKGLFALVAYLTWADETKEVITDNLPG-
SIRAVVNIFLVAKALLSYPLPFFAAVEVLEKSLFQEGSRAFFPACYSGDGRLKSWGLTLRCALVVFTLLMAIYVPHFALLMGLTGSLTGAGLCFLLPSLFHLRLLWRKLLWHQVFFDVAIFVIGGICSVSGFVHSLEGL---------------------
---------------------------------------------------------------IEAYRTNAED---------------------------------------------------------------------------------------
-------------------------

>NP 610938.1 vesicular GABA transporter Drosophila melanogaster
MSFIAKLK-ATPLPPLRNILNVAVQTARQQIPERKDYEQPPGSTAQQHHHSQQAQHKAMEAGMDGGDTTEMSSNPFRNAGSWTNDGEGGGDGDGEYRNEYQSTSFNEYDGRYQQTDGFRQGSIASEGSSFVCEGEGGG--
GCKIDEFQAAWNVTNAIQGMFIVSLPFAVLHGGYWAIVAMVGIAHICCYTGKVLVQCLYEPDPATGQMVRVRDSYVAIAKVCFGPKL---
GARAVSIAQLIELLMTCILYVVVCGDLLAGTYPQGSFDSRSWMLFVGIFLLPMGFLKSLKMVSTLSFWCTMSHIVINAVILGYCLLQIGDWGWSKVRFSIDMENFPISLGVIVFSYTSQIFLPTLEGNMIDRSKFNWMLDWSHIAAAVFKAGFGYICFLT
FQNDTQQVITNNLHSQGFKGMVNFFLVIKALLSYPLPYYAACELLERNFFRGPPKTKFPTIWNLDGELKVWGLGFRVGVIVSTILMAIFIPHFSILMGFIGSFTGTMLSFIWPCYFHIKIKGHLLDQKEIAKDYLIIGLGVLFGVIGIYDSGNAL-----
-------------------------------------------------------------------------------INAFEIGLPF----------------------------------------------------------------------
-----------------------------------------

>XP 014787249.1 vesicular inhibitory amino acid transporter isoform X1 Octopus bimaculoides
------MT----AKSIYYIEKLKSLIRQSS-----------TDETVHFVKYSQFE-NEKE-----LTEIFDDKKK---FGSITEESNGTA---------------SISDV------
KDGKQTLGSPHQTPVMDAVDAKLERNRITSWQAGWNVTNAIQGMFIVSFPYTVVQGGYWALVSMIVVAYICCHTGKILVECLYEVDS-NGYLVRVRHSYVEIAEEAMGKRF---
GGKIVNVAQLIELLMTCILYVLLSGELILGSFPESPISLTFWIMIATTPLLSYAFLKSLRGVSWLSFWCTMAHMAINAFILIFCFTRASEWQWKTVQIKVDIWTFPISIGIVVFSYTSQIFLPTLEENMEDRSRFHCMMHWTHIAAAFFKVLFSYICFLT
WGNGTQEVITNNLPTQGLKIIVNLILVAKSTLSYPLPYYAAVEMIQTTYFNGKPSTYFPSCFDENCRLKIWAGCLRIGLVLATMVMAIFIPHFAILMGLIGSFTGTMLSFVWPCFFHIKIKGKILPWYTKVLDISIIIIGVVVGSIGIYYSFRAL-----
-------------------------------------------------------------------------------SLAFRGIPTTPFHARDFHKSL------------------------------------------------------------
-----------------------------------------

>XP 019925215.2 vesicular inhibitory amino acid transporter-like Crassostrea gigas
---------------------------------------------------------------------------------------------------------------------------------------------------------------
MFLVSFPYALVQGGYWTLLVISVTAAICAHTSQLIVECLYDEDA-SGQKVRVRNSYVDIANRVWGPRV---
GKVLLTAAQIIELSFTCILYILVSGELLYGCFRSRDVSLAAWTVISTVPLLPCAFLQSIRRVSSLSFWCTMAHVVINAVIIVYCFTKGSDWHWDQMPVEINIFEFPVSLGIVVFSYTSQIFAPTLEGKMRNPGRFSRMTFCTHICAAVFKSVFAYVCFLT
WGKETKEVITNNLTIPSLKTAVDLVLVIKALLSYPLPYFATLEIIEQEFFTLFNNSCCTPCFDDKNKLDTWAAAMRIAFVLATMLLAVFLPHFSILMGLVGSFTGTMLSLVWPCHFYLQIHGQRLSWHKKFVNWIIIVLGLVVCCIGMFYSGLAL-----
-------------------------------------------------------------------------------HHAVHGQGVAMSMTFNLTQL-------------------------------------------------------------
-----------------------------------------

>XP 029641102.1 vesicular inhibitory amino acid transporter isoform X1 Octopus sinensis
------MT----AKSIYYIEKLKSLIRQSS-----------TDETVHFVKYSQFE-NEKE-----LTEIFDDKKK---FGSITDESNGSA---------------SMSDV------
KDSKQTLGSPHQTPVMDAVDTKLERNRITSWQAGWNVTNAIQGMFIVSFPYTVVQGGYWALVSMIVVAYICCHTGKILVECLYEVDS-NGYLVRVRHSYVEIAEEAMGKRF---
GGKIVNVAQLIELLMTCILYVLLSGELILGSFPESPISLTFWIMIATTPLLSYAFLKSLRGVSWLSFWCTMAHMAINAFILIFCFTRATEWQWKSVQIKVDVWTFPISIGIVVFSYTSQIFLPTLEENMEDRSRFHCMMHWTHIAAAFFKVLFSYICFLT
WGNGTQEVITNNLPTQSLKIVVNLILVAKSTLSYPLPYYAAVEMIQTTYFNGKPSTYFPSCFDENCRLKIWAGCLRIGLVLATMAMAIFIPHFAILMGLIGSFTGTMLSFVWPCFFHIRIKGKILPWYTKVLDISIIIIGVVVGSIGIYYSFRAL-----
-------------------------------------------------------------------------------SLAFRGIPTTPFHARDFHKSL------------------------------------------------------------
-----------------------------------------

>XP 009061033.1 hypothetical protein LOTGIDRAFT 193611 Lottia gigantea
---------------------------------------------------------------------------------------------------------------------------------------------------------------
MFIVSFPYAVVQGGYWAILSMIVVAYICCHTGKILVQCLYEENE-HGRKVRVRNSYVEIAEHVWGKRF---
GGRIVDCAQIIELLMTCILYVLLCGDLIKGSFPDSPLGLSSWIMVSTIPLLACAFLQSLQRVSMLSFWCTVAHMLINAIIIIYCFTRVTEWHWSDVQVRINIWTFPISLGIVVFSYTSQIFLPTLECNLEDKSKFTCMMHWTHLAAAVFKAGFSYIGFLT
WGFATKEVISNNLPTQVFKIIVNMILVAKAQLSYPLPYFAAVELLETSLFAGRPETVFPACIDNTHRLKIWALICRVALVVFTTLLAIIIPHFAILMGLIGSFTGTMLSFVWPCYFHLKIKWHTMSITYKILDIAIIVIGLACGSIGIYYSGHAL-----
-------------------------------------------------------------------------------TRAFQGLPPSPFHGSKHD---------------------------------------------------------------
-----------------------------------------

>XP 055885872.1 vesicular inhibitory amino acid transporter-like Biomphalaria glabrata
----MSWR----ERLAYLALNVKLQWRYRS-----------PEYNEEKLGFAKWQSQSQI-----IESHPMKNRL---FNHTGKNKSPSE---------------HA---------
ENPGEFSPFQDDLEVEEAEICGDPRDKITEWQAGWNVTNAIQGMFIVSFPFTVLQGGYWAVVAMVLVAYICCHTGNILVDCLYDLDP-MGHRVRVRSSYVEIATAVWGARY---
GARIVHCAQLIELLMTCILYVLLCGDLIQGSFPNTPFSLTSWILMCTTPLLACAFLTSLRRVSTLSMWCTVAHMLINAIILVYCFTKAGQWKWTDVQIRIDIWTFPISLGIIVFSYTSQIFLPTLEGKMRDRSKFKCMMTWTHVVAAVFKAFFSYVGFLT
WGLDTLEVVTNNLPYLSLKLIVNLILVCKAMLSYPLPYFASVELLESAFFKGKPSTCFPPCLDDSRQLKWWGLSLRLALVLFTCAMAIIVPHFALLMALIGSFTGTMLSFVWPCYFHLRLRWYVMSRTTRVLNIFIILLGLACGGIGIYYSAHAL-----
-------------------------------------------------------------------------------TRAFQGLPPEPLHL-------------------------------------------------------------------
-----------------------------------------

>XP 791315.4 vesicular inhibitory amino acid transporter Strongylocentrotus purpuratus
------------MMNSRYVNRFLDLFRLSG-----------GDESVPFARQDATELTVIP-----GASSSGDSDD----SPTVENGKPPG------------------GLRDSSNGGAVKQHGVDDK-
PSGEGDNSNIMRKQITAWDAGWNVTNAIQGMFLVALPYAVMHGGYWTVLSLVLAAIITCYTGLILVDCLYDTNAITGERVRVRETYVSIAEEVWGKRF---
ASRVVHTAQFIELIMTCILYLVLCGDLLYNTIRHTPLRESAWTLIACFLVLPCAFLRNLKAVSRSSFGNAIAHVIINVIILGFCFAQAGHWHWKDTSLRIHIHYFPVSLGIIVFSYTSHIFLPSLEGNMVDRRYFKRMMLWTHGLAGFFKAFFGYVAYLT
FGLSTQEVISDNLPTHSFRSIVNLVLVAKALLSFPLPYFAAVELLERAFFQGRPTTVLPSCYSHDGMLTVWSIPLRLLLICVSVLLAVFIPHFAILMGLIGSVTGTMLSFIWPCWFHLRLKWHELKLWNKVIDILIMLAGAGCGCIGIIYSFEAL-----

--------------------------------------------------------------------------------VKTYVES--------------------------------------------------------------------------------------------------------------------------------------------
>CAE1313291.1 SLC32A Sepia pharaonis
----------------------------------------------------------------------------------------------------------------------------------------------------------------------
MFIVSFPYTVLQGGYWAIVSIIVVAYICCHTGKILVECLYEVDS-SGYLVRVRRSYVEIAEEAMGKRF---
GGKIVNVAQLIELLMTCILYVLLSGELILGSFPESPITLTSWIMIATVPLLSYAFLKSLRGVSWLSFWCTMAHMAINAFILIFCFTNATEWHWKEVQVKVDIWTFPISMGIVVFSYTSQIFLPTLEGNMKDRSRFHCMMHWTHLTAAFFKALFAYICFLT
WGFNTQEVITNNLPTQGLKIVVNLILVAKSILSYPLPYYAAVEMIQSTYFSGKPATLFPSCFDESYRLKVWALCLRIGLVLATMMLAIFIPHFAILMGLIGSFTGTMLSFVWPCYFHIKIKWKVLPWYTRALDICIILIGATVGSIGMYYSFRALDFLSF
IFSFFHFFLSFIFFFLSAKSHIHTLSVCLSVCLSLSLSIYLSMLHGVLSIYLSIYLSIYLSIYLSITIFSFLSRSFHFFLSIYLSISIFSFLSRSFHFCLDLFISIYLSIYLSIYLSGSFHIYLSIYLSIYLDLFISVSIFSFVSLYLFIDLNLFISICF
DLCISIYLSIYLSIYLSIYLSRAVLQECATRNTQMFGVTNS

Vesicular glutamate transporter
>Obocki vGlut
MTARFVESIPRIKAALPSFNTDIPMKPINKIKGLIKKGQDEKTLVNYDELDETYGDDESPP--GPRKTLPMPGPLPPVPAPKCMCFQ--SRQRYLIALLSSLGFLISFGIRCNMGVAIVMMVNNDTDSDSSQNT---------------TEVRM----
PEFSWTPETIGIVDSSFFWGYIITQIPGGYLASKLPATRIFGAAIXISSFLNXFLPGAASVHYGLVMAVRILQGLVEGVTYPACHGIWRHWAPPLERSQLATISFCGSYAGAVLGMPLSGLLTQYLGWRSGFYCFGMMGIIWGICWYLLSHERPSTHPHI
TKEERHYIESSIGEPAV---
AVTTHNTPWKKFFTSKPVYAIMVANFCRSWTFYLLIIEQPTYFKEAFRFNVSQSGIISALPHLVMAIIVPFGGQLADFLRRNXYLXTTNVRKIFNCGGFGMEAVFLLGVGYTADRVTAIVCLTLAVGFSGFAISGFNVNHLDIAPRYASILMGLSNGVGT
LSGMLCPPITEMLTKIPDVS------SWEHVFLIASMVHFGGVIFYAIFASGELQPWAEPPPEDE----WKPEDTLKGDSY---------------------DKMPDYGTV---------------NEN----------G-------
PVYETKEEMVQRYARQ----PSDDXE-QENY-
>XP 014776384.1 vesicular glutamate transporter 1 isoform X1 Octopus bimaculoides
MTARFVESIPRIKAALPSFNTDIPMKPINKIKGLIKKSQDQKTLVNYDELDETYGDDESPP--GPRKTLPMPGPLPPVPAPKCMCFQ--SRQRYLIALLSSLGFLISFGIRCNMGVAIVMMVNNDTDSDSSQNT---------------TEVRM----
PEFSWTPETIGIVDSSFFWGYIITQIPGGYLASKLPATRIFGTAITISSFLNLFLPGAASVHYGLVMAVRILQGLVEGVTYPACHGIWRHWAPPLERSQLATISFCGSYAGAVLGMPLSGLLTQYLGWRSGFYCFGMMGIIWGICWYLLSHERPSTHPHI
TKEERHYIESSIGEPAV---
AITTHNTPWKKFFTSKPVYAIMVANFCRSWTFYLLIIEQPTYFKEAFRFNVSQSGIISALPHLVMAIIVPFGGQLADFLRRNGYLSTTNVRKIFNCGGFGMEAVFLLGVGYTADRVTAIVCLTLAVGFSGFAISGFNVNHLDIAPRYASILMGLSNGVGT
LSGMLCPPITEMLTKIPDVS------SWEHVFLIASMVHFGGVIFYAIFASGELQPWAEPPPEDE----WKPEDTLKGDSY---------------------DKMPDYGTV---------------NEN----------G-------
PVYETKEEMVQRYARQ----PSDDSE-QENY-
>XP 029646438.1 vesicular glutamate transporter 1-like isoform X1 Octopus sinensis
MTARFVESIPRIKAALPSFNTDIPMRPINKIKGLIKKGQDEKTLVNYDELDETYGDDESPP--GPRKTLPMPGPLPPVPAPKCMCFQ--SRQRYLIALLSSLGFLISFGIRCNMGVAIVMMVNNDTDIDSSQNT---------------TEVRM----
PEFSWTPETIGIVDSSFFWGYIITQIPGGYLASKVPATRIFGAAITISSFLNLFLPGAASVHYGLVMAVRILQGLVEGVTYPACHGIWRHWAPPLERSQLATISFCGSYAGAVLGMPLSGLLTQYLGWRSGFYCFGMMGIIWGIFWYLLSHERPSTHPHI
TKEERHYIESSIGEPAG---
AITTYNTPWKKFFTSKPVYAIMVANFCRSWTFYLLIIEQPTYFKEAFRFNVSQSGIVSALPHLVMAIIVPFGGQLADFLRRNGYLSTTNVRKIFNCGGFGMEAVFLLGVGYTADRVTAIICLTLAVGFSGFAISGFNVNHLDIAPRYASILMGLSNGVGT
LSGMLCPPITEMLTKIPDVS------SWEHVFLIASMVHFGGVIFYAIFASGELQPWAEPPPEDE----WKPEDTLKGDSY---------------------DKMPDYGTV---------------NEN----------G-------
PVYETKEEMVQRYARQ----PSDDSE-QENY-
>NP 001297420.1 vesicular glutamate transporter 2-like Aplysia californica
---MPFGAFNNLKDRVLQPSKEAAQLAKDSIHRLVHKQTDEKSLVKYDELDEAYTEADELPTKSEKSFIPSKSYLP----TSCPCMRCNVSKRYQIALLSSVGFVLSFGIRCNLGVAVLDMTKNNTVDLDGDGI---------------
IRLNDSIQEPEFNWTPETVGVVDSSFFWGYIVTQIPGGYLASRVSATRLFGLAIGISACLNLLLPGAAEVHYGLVISVRILQGLVEGVTYPACHGIWRHWAPPLERSMLATISFCGSYAGAVLGMPISGILTRYLGWQSGFYAFGIAGAIWSAAWWFLSY
EKPSTHPTITEEERVYIENSIGESTSV--ISKSIPTPWLKFFTSTPVWAIMVANFCRSWTFYLLIISQPAYFEEVFGFKIDESGTLSALPHLVMAIIVPIGGMIADFLRRR-
ILTTTVVRKMFNCGGFGMEAIFLLGVSFTTTTAPAIVCLTLAVGFSGFAISGFNVNHLDIAPRYASILMGLSNSVGTLAGMLCPIVVQAITKDAREKPENAQQEWQYVFLIASMIHFAGVIFYGIFASGEKQPWADPPGEET----
WRPEHTLKPDDNSWKFSNHNGHQY---------GGSTNYGAT----------------DKGYDDYIPTQYS-------PVYETKETFVQKPAKE----KYYSDDSEKDF-
>XP 055900160.1 vesicular glutamate transporter 2-like Biomphalaria glabrata
---MPFGAFTNLKDRVLQPSKDATQLAKDSIHRLVSKQNDEKSLVKYDELDDAYTEADELPSKSEKSFIPRKSYIP----RNCPCFRCNVAKRYQIALLSSVGFLLSFGIRCNMGVAVLDMTKNSTRDLDGDGV---------------
ISLNDSIQEPEFNWTPETVGVVDSSFFWGYIVTQIPGGYLASRISATRLFGLAIGISGCLNLLLPGAAEIHYGLVISVRILQGLVEGVTYPACHGIWRHWAPPLERSMLATISFCGSYAGAVLGMPLSGILTRYLGWQSGFYVFGVLGAIWSVAWWFLSY
EKPSTHPTITEEERVYIETSIGETTSM--SMKNIGTPWVAFFTSMPVWAIMVANFCRSWTFYLLIISQPAYFEEVFGFKIDESGTLSALPHLVMAIIVPIGGQIADFLRRR-
VLTTTVVRKIFNCGGFGMEAVFLLGVSFTRDTAPAVVCLTLAVGFSGFAISGFNVNHLDIAPRYASILMGLSNSVGTLAGMLCPIVVQAITKDARNNVEVAKTEWQYVFLIASLIHFAGVIFYGIFASGEKQPWADPPGEET----
WRPEHTLRPNDESWKFSNHGNPHYD--------GGKTNYGAT---------------TDKGYDDFVPTQYS-------PVYETKESFVQKPAKE----KYYSDDSDRGF-
>XP 011431416.2 vesicular glutamate transporter 1 isoform X2 Crassostrea gigas
------------MNKLDGLIEVLRLRDI----RVFKKDAEEKSFVKYDELDEAYSEEEDK----KSRFLGYC--------PSCTCCRC-
LAQRYLVAILSCVGFLISFGIRCNMGVAIVMMTKNNTYDDDDKPTPAPILVKDGNSTYYNRTVPTLPHLPQFLWTPETIGIVDSSFFWGYIVTQIPGGYLASRVPANRIFGLAIGISSFLNILLPGAAQVHYGLVMAVRILQGLVEGVTYPACHGIWRHW
APPLERSRLATISFCGSYAGAVLGMPLSGILSKTLGWQSGFYVFGIIGMIWCIFWMIFSFEKPSTNPHITQEERIFIETSIAESGCL--
VDRDMKTPWFKFITSMPVYAIIVANFCRSWTFYLLIISQPMYFSEVFHFNVSKSGVLSALPHLVMAIIVPIGGFMADKLRQR-
LLTTTTVRKIFNCGGFGLEACFLLGVAITRNTTTAIVCLTLAVGSSGFAISGFNVNHLDIAPRFASILMGLSNGIGTLSGMLCPVVTELLTKKGTAD------EWQTVFIIASVVHFLGIIFYGIFASGEKQPWAEPPEEDR----
WRPEDTLKPEGSGKLFSYGAFHGLDSQADDKMNGGVISNGHVGNSSTGNGEYGGFDKSEEYYNNGYGGDYGKRMSFDAPLYSTKEELVQVQSKD----SYLND--KREL-
>XP 009062336.1 hypothetical protein LOTGIDRAFT 128007 Lottia gigantea
------------------------------------------------------------MSSYIPS----------RCPCIKCNISKRYQIALLSSVGFILSFGIRCNLGVAILSMTTNETIQFTGDTK---------------TKAQF---
NLTFSWTPETVGIVDSSFFWGYIITQIPGGYLASRLPANRLFGLAIGISSFLNLFLPAAAKVHYGMVITVRILQGLVEGVTYPATHGIWRHWAPPLERSKLATISFCGSYAGAVLGMPVSGVLTQYLGWQSGFYFFGALGLLWTVIWWFFSFERPSTHPT
ISEAEKIYIETSIGENTSV--ILKDVKTPWLKFFTSMPVYAIMVANFCRSWTFYLLIISQPTYFLKVFRFEVSKSGTLSALPHLVMAIIVPIGGQIADFLRMR-
ILTTTVVRKIFNCGGFGMEAVFLLGVAFARDTATSIACLTLAVGFSGFAISGFNVNHLDIAPRYASILMGLSNSVGTLSGMLCPIVVQRITKGQERDFETATREWEYVFLIASMIHFAGVIFYAIFASGEKQPWADPPSEEE----
WRPEHSLKPNTNNFTYKKYGSTQ----------NESIDYNTS---------------PNQ----------S-------PVYETREEFVQKESKD----SYMRDG-GGDF-
>XP 006540664.1 vesicular glutamate transporter 2 isoform X1 Mus musculus
------------------------------MLVLEKKQDNRETI---ELTE-----DGKPLEVPEKKAPLC---------DCTCFG--LPRRYIIAIMSGLGFCISFGIRCNLGVAIVDMVNNSTIHRGGKVI---------------KEKAK------
FNWDPETVGMIHGSFFWGYIITQIPGGYIASRLAANRVFGAAILLTSTLNMLIPSAARVHYGCVIFVRILQGLVEGVTYPACHGIWSKWAPPLERSRLATTSFCGSYAGAVIAMPLAGILVQYTGWSSVFYVYGSFGMVWYMFWLLVSYESPAKHPTITD
EERRYIEESIGESANLLGAMEKFKTPWRKFFTSMPVYAIIVANFCRSWTFYLLLISQPAYFEEVFGFEISKVGMLSAVPHLVMTIIVPIGGQIADFLRSKQILSTTTVRKIMNCGGFGMEATLLLVVGYSHTRGVAISFLVLAVGFSGFAISGFNVNHLD

```
IAPRYASILMGISNGVGTLSGMVCPIIVGAMTKNKSRE------EWQYVFLIAALVHYGGVIFYALFASGEKQPWADPEETSEEKCGFIHEDELDEETGDITQNYINY------------GTTKSYGAT---------------SQE----NGGWPNG--
-------WEKKEEFVQEGAQD----AYTYKD-RDDYS
>NP 065079.1 vesicular glutamate transporter 2 Homo sapiens
----MESVKQRILAPGKEGLKNFAGKSLGQIYRVLEKKQDTGETI---ELTE-----DGKPLEVPERKAPLC---------DCTCFG--LPRRYIIAIMSGLGFCISFGIRCNLGVAIVDMVNNSTIHRGGKVI---------------KEKAK------
FNWDPETVGMIHGSFFWGYIITQIPGGYIASRLAANRVFGAAILLTSTLNMLIPSAARVHYGCVIFVRILQGLVEGVTYPACHGIWSKWAPPLERSRLATTSFCGSYAGAVIAMPLAGILVQYTGWSSVFYVYGSFGMVWYMFWLLVSYESPAKHPTITD
EERRYIEESIGESANLLGAMEKFKTPWRKFFTSMPVYAIIVANFCRSWTFYLLLISQPAYFEEVFGFEISKVGMLSAVPHLVMTIIVPIGGQIADFLRSKQILSTTTVRKIMNCGGFGMEATLLLVVGYSHTRGVAISFLVLAVGFSGFAISGFNVNHLD
IAPRYASILMGISNGVGTLSGMVCPIIVGAMTKNKSRE------EWQYVFLIAALVHYGGVIFYAIFASGEKQPWADPEETSEEKCGFIHEDELDEETGDITQNYINY------------GTTKSYGAT---------------TQA----NGGWPSG--
-------WEKKEEFVQGEVQD----SHSYKD-RVDYS
>NP 001076304.1 vesicular glutamate transporter 3 Danio rerio
---MPLGGFAGLKEKLNPGKEELK-NNVGDSLGNLQKKIDGSNVTEEDNIELT---EDGRPVAAPKRSPPLL---------DCGCFG--LPKRYIIAMLSGLGFCISFGIRCNLGVAIVEMVNNNTVYINGTAV---------------MQPAQ------
FNWDPETVGLIHGSFFWGYIVTQIPGGFISNKLAANRVFGAAIFLTSVLNMFIPSAARVHYGCVMFVRILQGLVEGVTYPACHGMWSKWAPPLERSRLATTSFCGSYAGAVIAMPLAGILVQYVGWPSVFYIYGVFGIIWYIFWILLAYNSPAVHPTISE
EERNYIETSIGEGANLMSSTEKFKTPWREFFTSMPVYAIIVANFCRSWTFYLLLISQPAYFEEVFGFPISKVGILSAVPHMVMTIIVPIGGQLADFLRSRKILSTTTVRKIMNCGGFGMEATLLLVVGFSHTRAVAISFLILAVGFSGFAISGFNVNHLD
IAPRYASILMGISNGVGTLSGMVCPLIVGALTKHKTRL------EWQHVFVIASMVHYTGVIFYAIFASGEKQDWADPENTSDEKCGIIGEDELADETEPSSDSGLA-------------TRQKTYGTT---------------DNS-----SGRKQG--
-------WKKKRGVTMQAEDDHESNHYENGEYQTQYQ
```

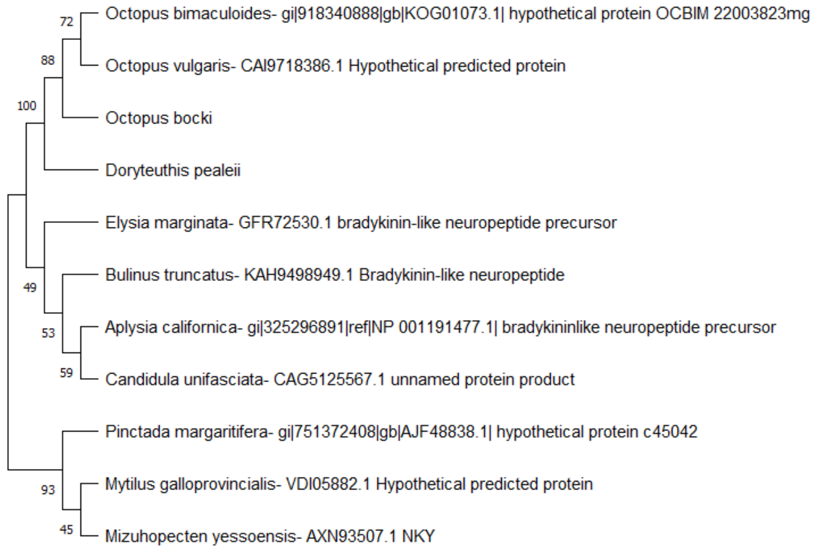


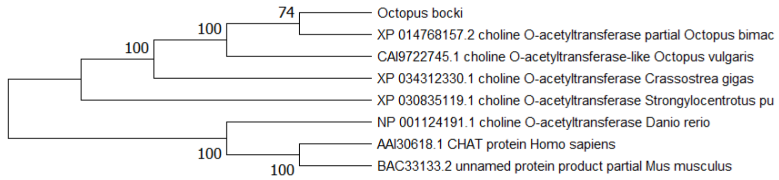

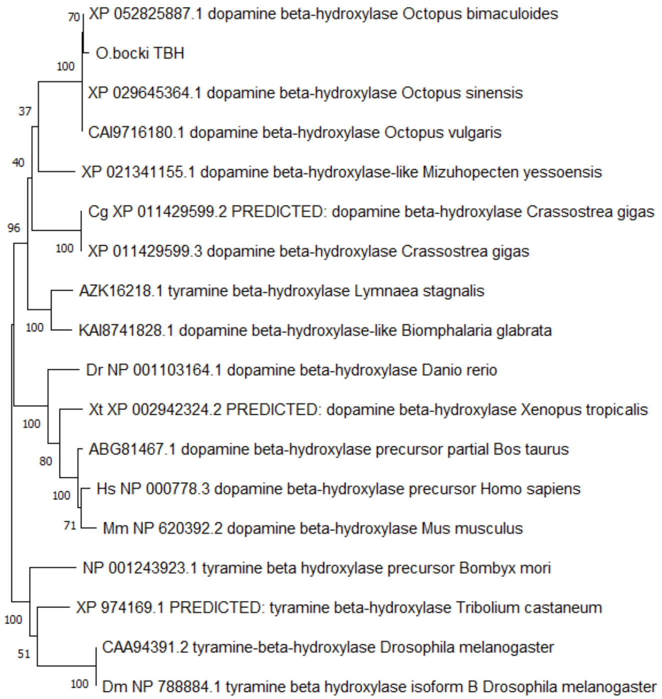

70
XP 052825887.1 dopamine beta-hydroxylase Octopus bimaculoides
O.bocki TBH
100
XP 029645364.1 dopamine beta-hydroxylase Octopus sinensis
37
CAI9716180.1 dopamine beta-hydroxylase Octopus vulgaris
40
XP 021341155.1 dopamine beta-hydroxylase-like Mizuhopecten yessoensis
Cg XP 011429599.2 PREDICTED: dopamine beta-hydroxylase Crassostrea gigas
96
100
XP 011429599.3 dopamine beta-hydroxylase Crassostrea gigas
AZK16218.1 tyramine beta-hydroxylase Lymnaea stagnalis
100
KAI8741828.1 dopamine beta-hydroxylase-like Biomphalaria glabrata
Dr NP 001103164.1 dopamine beta-hydroxylase Danio rerio
Xt XP 002942324.2 PREDICTED: dopamine beta-hydroxylase Xenopus tropicalis
100
80
ABG81467.1 dopamine beta-hydroxylase precursor partial Bos taurus
100
Hs NP 000778.3 dopamine beta-hydroxylase precursor Homo sapiens
71
Mm NP 620392.2 dopamine beta-hydroxylase Mus musculus
NP 001243923.1 tyramine beta hydroxylase precursor Bombyx mori
100
XP 974169.1 PREDICTED: tyramine beta-hydroxylase Tribolium castaneum
51
CAA94391.2 tyramine-beta-hydroxylase Drosophila melanogaster
100
Dm NP 788884.1 tyramine beta hydroxylase isoform B Drosophila melanogaster

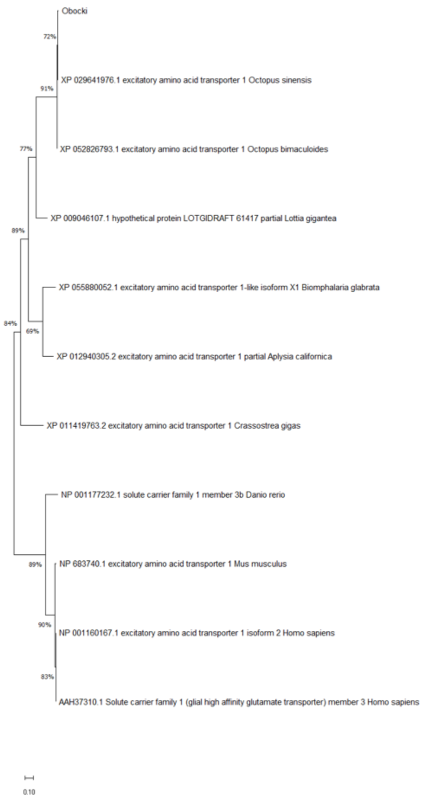
Obocki
72%
XP 029641976.1 excitatory amino acid transporter 1 Octopus sinensis
91%
XP 052826793.1 excitatory amino acid transporter 1 Octopus bimaculoides
77%
XP 009046107.1 hypothetical protein LOTGIDRAFT 61417 partial Lottia gigantea
89%
XP 055880052.1 excitatory amino acid transporter 1-like isoform X1 Biomphalaria glabrata
84%
69%
XP 012940305.2 excitatory amino acid transporter 1 partial Aplysia californica
XP 011419763.2 excitatory amino acid transporter 1 Crassostrea gigas
NP 001177232.1 solute carrier family 1 member 3b Danio rerio
89%
NP 683740.1 excitatory amino acid transporter 1 Mus musculus
90%
NP 001160167.1 excitatory amino acid transporter 1 isoform 2 Homo sapiens
83%
AAH37310.1 Solute carrier family 1 (glial high affinity glutamate transporter) member 3 Homo sapiens
0.10

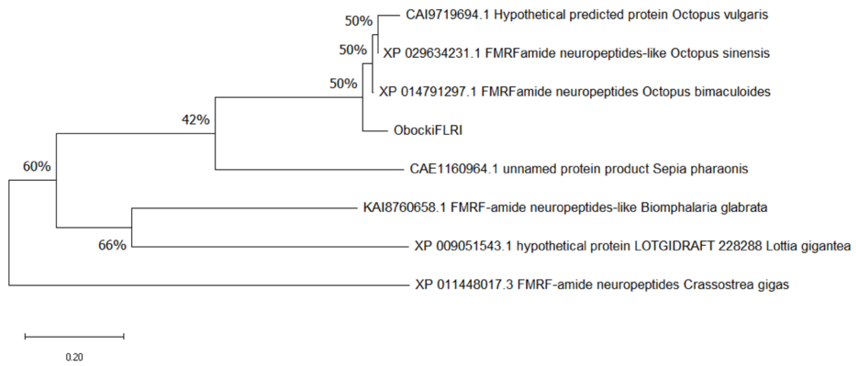
50%
CAI9719694.1 Hypothetical predicted protein Octopus vulgaris
50%
XP 029634231.1 FMRFamide neuropeptides-like Octopus sinensis
50%
XP 014791297.1 FMRFamide neuropeptides Octopus bimaculoides
42%
ObockiFLRI
60%
CAE1160964.1 unnamed protein product Sepia pharaonis
KAI8760658.1 FMRF-amide neuropeptides-like Biomphalaria glabrata
66%
XP 009051543.1 hypothetical protein LOTGIDRAFT 228288 Lottia gigantea
XP 011448017.3 FMRF-amide neuropeptides Crassostrea gigas
0.20

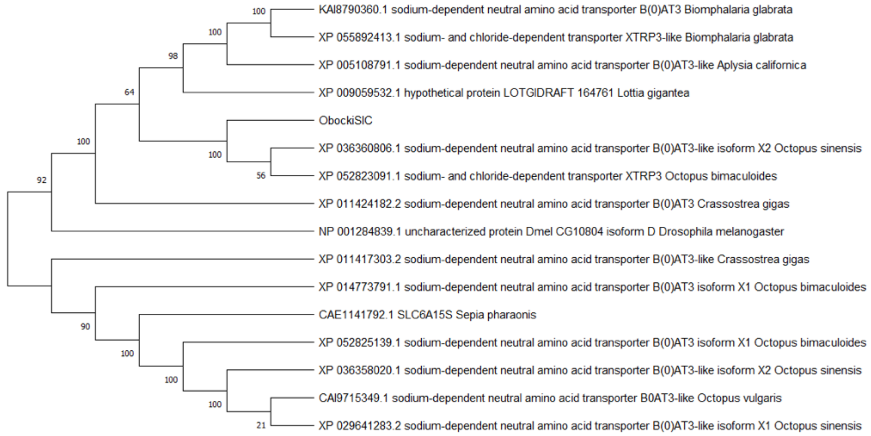
KAI8790360.1 sodium-dependent neutral amino acid transporter B(0)AT3 Biomphalaria glabrata
XP 055892413.1 sodium- and chloride-dependent transporter XTRP3-like Biomphalaria glabrata
XP 005108791.1 sodium-dependent neutral amino acid transporter B(0)AT3-like Aplysia californica
XP 009059532.1 hypothetical protein LOTGIDRAFT 164761 Lottia gigantea
ObockiSIC
XP 036360806.1 sodium-dependent neutral amino acid transporter B(0)AT3-like isoform X2 Octopus sinensis
XP 052823091.1 sodium- and chloride-dependent transporter XTRP3 Octopus bimaculoides
XP 011424182.2 sodium-dependent neutral amino acid transporter B(0)AT3 Crassostrea gigas
NP 001284839.1 uncharacterized protein Dmel CG10804 isoform D Drosophila melanogaster
XP 011417303.2 sodium-dependent neutral amino acid transporter B(0)AT3-like Crassostrea gigas
XP 014773791.1 sodium-dependent neutral amino acid transporter B(0)AT3 isoform X1 Octopus bimaculoides
CAE1141792.1 SLC6A15S Sepia pharaonis
XP 052825139.1 sodium-dependent neutral amino acid transporter B(0)AT3 isoform X1 Octopus bimaculoides
XP 036358020.1 sodium-dependent neutral amino acid transporter B(0)AT3-like isoform X2 Octopus sinensis
CAI9715349.1 sodium-dependent neutral amino acid transporter B0AT3-like Octopus vulgaris
XP 029641283.2 sodium-dependent neutral amino acid transporter B(0)AT3-like isoform X1 Octopus sinensis
100
100
98
64
100
100
56
92
90
100
100
100
21

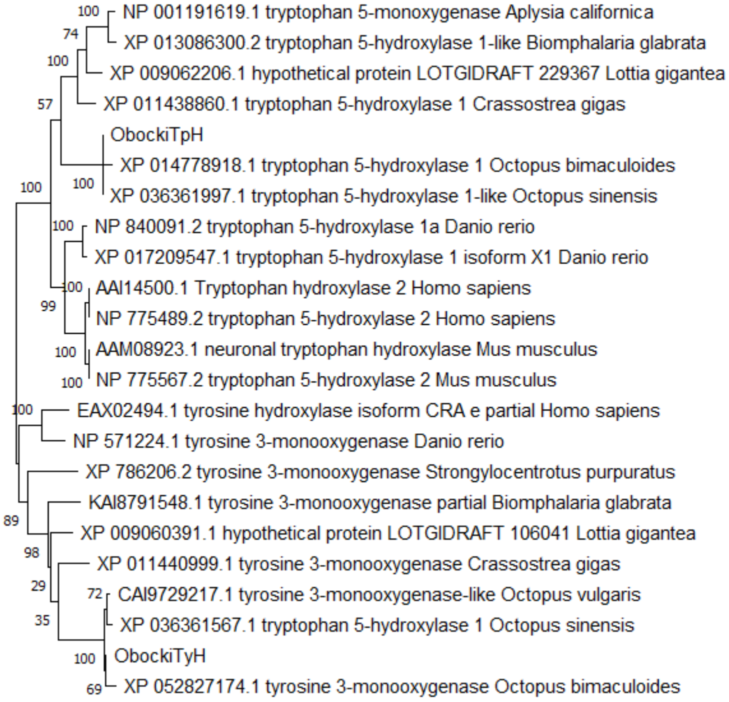
NP 001191619.1 tryptophan 5-monoxygenase Aplysia californica
XP 013086300.2 tryptophan 5-hydroxylase 1-like Biomphalaria glabrata
XP 009062206.1 hypothetical protein LOTGIDRAFT 229367 Lottia gigantea
XP 011438860.1 tryptophan 5-hydroxylase 1 Crassostrea gigas
ObockiTpH
XP 014778918.1 tryptophan 5-hydroxylase 1 Octopus bimaculoides
XP 036361997.1 tryptophan 5-hydroxylase 1-like Octopus sinensis
NP 840091.2 tryptophan 5-hydroxylase 1a Danio rerio
XP 017209547.1 tryptophan 5-hydroxylase 1 isoform X1 Danio rerio
AAI14500.1 Tryptophan hydroxylase 2 Homo sapiens
NP 775489.2 tryptophan 5-hydroxylase 2 Homo sapiens
AAM08923.1 neuronal tryptophan hydroxylase Mus musculus
NP 775567.2 tryptophan 5-hydroxylase 2 Mus musculus
EAX02494.1 tyrosine hydroxylase isoform CRA e partial Homo sapiens
NP 571224.1 tyrosine 3-monooxygenase Danio rerio
XP 786206.2 tyrosine 3-monooxygenase Strongylocentrotus purpuratus
KAI8791548.1 tyrosine 3-monooxygenase partial Biomphalaria glabrata
XP 009060391.1 hypothetical protein LOTGIDRAFT 106041 Lottia gigantea
XP 011440999.1 tyrosine 3-monooxygenase Crassostrea gigas
CAI9729217.1 tyrosine 3-monooxygenase-like Octopus vulgaris
XP 036361567.1 tryptophan 5-hydroxylase 1 Octopus sinensis
ObockiTyH
XP 052827174.1 tyrosine 3-monooxygenase Octopus bimaculoides
100
74
100
57
100
100
100
100
99
100
100
100
89
98
29
35
72
100
69

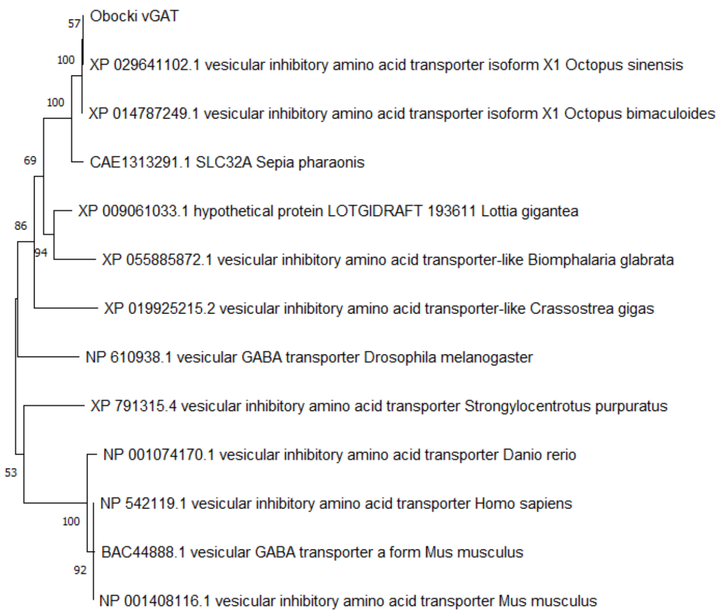
Obocki vGAT
XP 029641102.1 vesicular inhibitory amino acid transporter isoform X1 Octopus sinensis
XP 014787249.1 vesicular inhibitory amino acid transporter isoform X1 Octopus bimaculoides
CAE1313291.1 SLC32A Sepia pharaonis
XP 009061033.1 hypothetical protein LOTGIDRAFT 193611 Lottia gigantea
XP 055885872.1 vesicular inhibitory amino acid transporter-like Biomphalaria glabrata
XP 019925215.2 vesicular inhibitory amino acid transporter-like Crassostrea gigas
NP 610938.1 vesicular GABA transporter Drosophila melanogaster
XP 791315.4 vesicular inhibitory amino acid transporter Strongylocentrotus purpuratus
NP 001074170.1 vesicular inhibitory amino acid transporter Danio rerio
NP 542119.1 vesicular inhibitory amino acid transporter Homo sapiens
BAC44888.1 vesicular GABA transporter a form Mus musculus
NP 001408116.1 vesicular inhibitory amino acid transporter Mus musculus
57
100
100
69
86
94
53
100
92
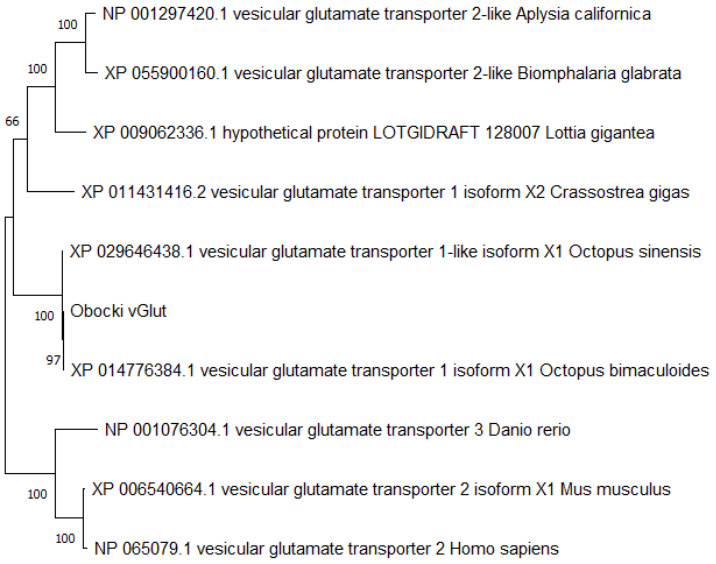
NP 001297420.1 vesicular glutamate transporter 2-like Aplysia californica
XP 055900160.1 vesicular glutamate transporter 2-like Biomphalaria glabrata
XP 009062336.1 hypothetical protein LOTGIDRAFT 128007 Lottia gigantea
XP 011431416.2 vesicular glutamate transporter 1 isoform X2 Crassostrea gigas
XP 029646438.1 vesicular glutamate transporter 1-like isoform X1 Octopus sinensis
Obocki vGlut
XP 014776384.1 vesicular glutamate transporter 1 isoform X1 Octopus bimaculoides
NP 001076304.1 vesicular glutamate transporter 3 Danio rerio
XP 006540664.1 vesicular glutamate transporter 2 isoform X1 Mus musculus
NP 065079.1 vesicular glutamate transporter 2 Homo sapiens
100
100
66
100
97
100
100
